# Signatures of archaic adaptive introgression in present-day human populations

**DOI:** 10.1101/045237

**Authors:** Fernando Racimo, Davide Marnetto, Emilia Huerta-Sánchez

## Abstract

Comparisons of DNA from archaic and modern humans show that these groups interbred, and in some cases received an evolutionary advantage from doing so. This process - adaptive introgression - may lead to a faster rate of adaptation than is predicted from models with mutation and selection alone. Within the last couple of years, a series of studies have identified regions of the genome that are likely examples of adaptive introgression. In many cases, once a region was ascertained as being introgressed, commonly used statistics based on both haplotype as well as allele frequency information were employed to test for positive selection. Introgression by itself, however, changes both the haplotype structure and the distribution of allele frequencies, thus confounding traditional tests for detecting positive selection. Therefore, patterns generated by introgression alone may lead to false inferences of positive selection. Here we explore models involving both introgression and positive selection to investigate the behavior of various statistics under adaptive introgression. In particular, we find that the number and allelic frequencies of sites that are uniquely shared between archaic humans and specific present-day populations are particularly useful for detecting adaptive introgression. We then examine the 1000 Genomes dataset to characterize the landscape of uniquely shared archaic alleles in human populations. Finally, we identify regions that were likely subject to adaptive introgression and discuss some of the most promising candidate genes located in these regions.

## 1. Introduction

There is now a large body of evidence supporting the idea that certain modern human populations admixed with archaic groups of humans after expanding out of Africa. In particular, non-African populations have 1 − 2% Neanderthal ancestry [1, 2], while Melanesians and East Asians have 3% and 0.2% ancestry, respectively, from Denisovans [3, 4, 2].

Recently, it has become possible to identify the fragments of the human genome that were introgressed and survive in present-5 day individuals [5, 6, 2, 7, 8, 7]. Researchers have also detected which of these introgressed regions are present at high frequencies in certain present-day non-African populations. Some of these regions are likely to have undergone positive selection in those populations after they were introgressed, a phenomenon known as adaptive introgression (AI). One particularly striking example of AI is the gene *EPAS1* [9] which confers a selective advantage in Tibetans by making them less prone to hypoxia at high altitudes [10, 11, 12, 13, 14, 15, 16, 17]. The selected Tibetan haplotype is likely to have been introduced in the human gene pool by Denisovans or a population closely related to them [18, 19].

In this study, we first use simulations to assess the power to detect AI using different exploratory summary statistics that do not require the introgressed fragments to be identified *a priori*. Some of these are inspired by the signatures observed in *EPAS1*, which contains an elevated number of sites with alleles uniquely shared between the Denisovan genome and Tibetans. We then apply these statistics to real human genomic data from phase 3 of the 1000 Genomes Project [20], to detect AI in human populations, and find candidate genes. While these statistics are sensitive to adaptive introgression, they may also be sensitive to other phenomena that generate genomic patterns similar to those generated by AI, like ancestral population structure and incomplete lineage sorting. These processes, however, should not generate long regions of the genome where haplotypes from the source and the recipient population are highly similar. As additional confirmation that the candidates we found with our statistics are generated by AI, we explored the haplotype structure of some of the most promising candidates, and used a probabilistic method [21] that infers introgressed segments along the genome by looking at the spatial arrangement of SNPs that are consistent with introgression. This allows us to verify that the candidate regions contain introgressed haplotypes at high frequencies: a hallmark of AI.

## 2. Results

### 2.1. Statistics for detecting AI

We began by evaluating the performance of various statistics at detecting AI. In addition to testing statistics that have already 25 been previously defined in the literature (*D*, *f_D_*, *D*′, *r*^2^, *π*), we define three new types of statistics that we find are particularly powerful at detecting AI (Table 1). We briefly describe the new statistics here, but more extensive descriptions of all tested statistics can be found in the Methods section below.

**Table 1:** Summary statistics mentioned in the main text.

| Statistic | Explanation | Reference |
| --- | --- | --- |
| $D$ | D-statistic: measures excess allele sharing between a test population and an outgroup using a sister population that is more closely related to the test than the outgroup | [1][100] |
| $f_D$ | Similar to the D-statistic, but serves to better control for local variation in diversity patterns if one is interested in finding loci with excess ancestry from an admixing population. | [103] |
| $R_D$ | Average ratio of the sequence divergence between an individual from the source population and an individual from the admixed population, and the sequence divergence between an individual from the source population and an individual from the non-admixed population. This is computed by taking the average over all pairs of admixed and non-admixed individuals. | This study |
| $U_{A,B,C}(w, x, y)$ | Number of sites in which any allele is at a frequency lower than $w$ in panel $A$, higher than $x$ in panel $B$ and equal to $y$ in panel $C$. | This study |
| $U_{A,B,C,D}(w, x, y, z)$ | Number of sites in which any allele is at a frequency lower than $w$ in panel $A$, higher than $x$ in panel $B$, equal to $y$ in panel $C$ and equal to $z$ in panel $D$. | This study |
| $Q95_{A,B,C}(w, y)$ | 95% quantile of the distribution of derived allele frequencies in panel $B$, for sites where the derived allele is at a frequency lower than $w$ in panel $A$ and equal to $y$ in panel $C$. | This study |
| $Q95_{A,B,C,D}(w, y, z)$ | 95% quantile of the distribution of derived allele frequencies in panel $B$, for sites where the derived allele is at a frequency lower than $w$ in panel $A$, equal to $y$ in panel $C$ and equal to $z$ in panel $D$. | This study |
| $\pi$ | Expected heterozygosity, measured as the average of $2p(1-p)$ over all sites in a window, where $p$ is the frequency of an arbitrarily chosen allele. | [109] |
| $D'_{[intro]}$ | A measure of linkage disequilibrium. Computed as $D/D_{max}$ where $D = p_{XY} - p_X p_Y$, $p_{XY}$ is the frequency of haplotype $XY$, $p_X$ is the frequency of allele $X$, $p_Y$ is the frequency of allele $Y$, and $D_{max}$ is the maximum theoretical value that $D$ can take. $D'_{[intro]}$ is computed only using frequencies from the introgressed panel. When we compute this in a window, the value is obtained by taking the average over all pairs of SNPs. | [110] |
| $D'_{[comb]}$ | $D'$ computed using haplotype and allele frequencies from the combination of the introgressed and non-introgressed panels. | [110] |
| $r^2_{[intro]}$ | A measure of linkage disequilibrium. Computed as $D^2/(p_X(1-p_X)p_Y(1-p_Y))$. $r^2_{[intro]}$ is computed only using frequencies from the introgressed panel. When we compute this in a window, the value is obtained by taking the average over all pairs of SNPs. | [111] |
| $r^2_{[comb]}$ | $r^2$ computed using haplotype and allele frequencies from the combination of the introgressed and non-introgressed panels. | [111] |

First, in a region under AI, one would expect the sequence divergence between an individual from the source population and an admixed individual to be smaller than the sequence divergence between an individual from the source population and a non-admixed individual. Thus, we define *R_D_* as the average ratio of the sequence divergence between a panel from the source population and a panel from the admixed population, over the sequence divergence between the source panel and a non-admixed panel, computed in a genomic window of the genome.

Second, under AI we would expect a large number of sites containing archaic alleles at high frequency in the admixed population, but absent or at low frequency in a non-admixed population. Therefore, we define the statistic *U_A,B,C_*(*w, x, y*) to be equal to the number of sites within a genomic window where a sample C from an archaic source population has a particular allele at frequency y, and that allele is at a frequency smaller than w in a panel A of a non-admixed population but larger than x in a panel B of an admixed population (Figure 1). Throughout the text, we denote panels A, B and C as the “outgroup”, “target” and “bait” panels, respectively. If we have samples from two different archaic populations (for example, a Neanderthal genome and a Denisova genome), we can define *U_A,B,C,D_*(*w, x, y, z*) as the number of sites where the archaic sample C has a particular allele at frequency y and the archaic sample D has that allele at frequency z, while the same allele is at a frequency smaller than w in an outgroup panel A and larger than x in a target panel B (Figure S1).

**Figure 1:**
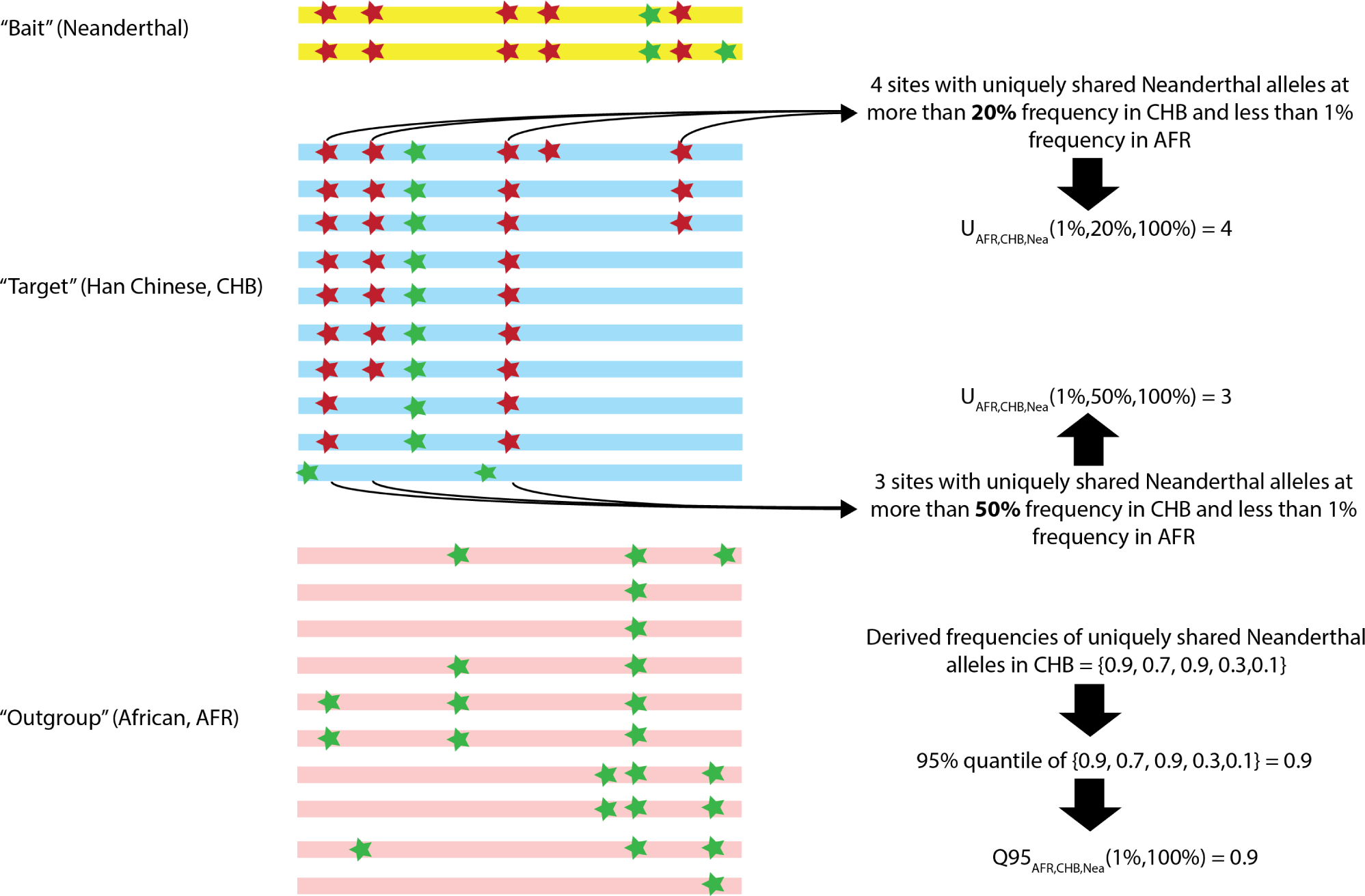
Schematic illustration of the way the *U_A,B,C_* and *Q*95_*A,B,C*_ statistics are calculated.

**Figure 2:**
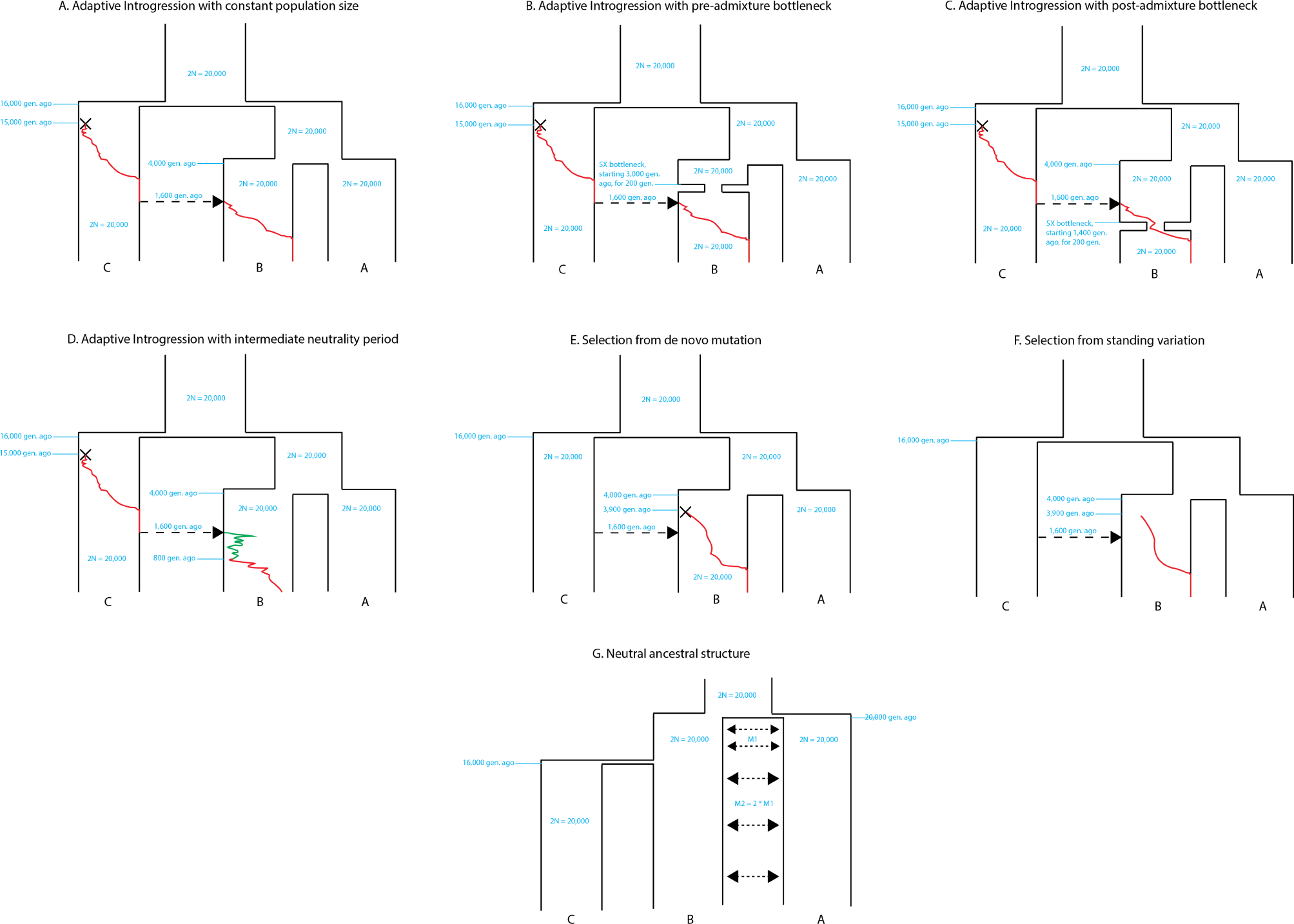
Demographic models described in the main text.

Finally, if we do not want to set a hard cutoff for what we consider “high-frequency” archaic alleles, we can just take a summary statistic of the site-frequency spectrum in the target panel, conditional on the archaic allele being at low frequency in the outgroup. This statistic should be high when a region contains many alleles at especially high frequencies in the target. We therefore define *Q*95_*A,B,C*_ (*w, y*) to be equal to the 95^*th*^ percentile of derived frequencies in a target panel B of all SNPs that have a derived allele frequency y in the bait (archaic) panel C, but where the derived allele is at a frequency smaller than w in an outgroup panel A from a non-admixed population (Figure 1).

### 2.2. Simulations under AI

We use simulations to assess the performance of the statistics mentioned above at detecting AI. Figures S3, S4 and S5 show 50 the distribution of statistics that rely on patterns of shared allele configurations between source and introgressed populations (*π*, *D*, *f_D_*, *U_A,B,C_*, *Q*95_*A,B,C*_ and *R_D_*), for different choices of the selection coefficient s, and under 2%, 10% and 25% admixture rates, respectively. For *Q*95_*A,B,C*_(*w*, 100%) and *U_A,B,C_*(*w, x,* 100%), we tested different choices of the outgroup cutoff w (1%, 10%) and the target cutoff x (0%, 20%, 50% and 80%).

Some statistics, like *Q*95_*A,B,C*_ (1%, 100%) and *f_D_* show strong separation between the selection regimes. For example, with an admixture rate of 2%, *Q*95_*A,B,C*_ (1%, 100%) has 100% sensitivity at a specificity of 99%, for both s=0.1 and s=0.01. Some parameterizations of the U statistic are not as effective, however. For example, *U_A,B,C_* (1%, 0%, 100%) shows some power when the admixture rate is low (2%), but almost no power when the admixture rate is high (25%). This is because setting the minimum frequency of the archaic allele in the test population at *x* = 0% means that any site with some archaic allele in the test panel will be counted, regardless of the allele frequency, so long as the archaic allele is at low frequency in the outgroup panel. At high admixture rates, low- and medium-frequency archaic alleles would naturally occur under neutrality, so they would not be informative about AI.

We also evaluated the effectiveness of LD-based statistics at detecting AI (Figure S6). We tested the performance of *D*′ and *r*^2^ by either computing each of these in the admixed panel only (*D*′[*intro*],*r*^2^[*intro*]) or in the combination of the admixed and non-admixed panels (*D*′[*comb*], *r*^2^[*comb*]). While *D*′[*intro*], *D*′[*comb*] and *r*^2^[*comb*] are modestly increased by AI, this is not the case with *r*^2^[*intro*] under strong selection and admixture regimes. This is because *r*^2^ will tend to decrease if the minor allele frequency is very small, which will occur if this frequency is only measured in the population undergoing AI. In general, these statistics are not as powerful for detecting AI as allele configuration statistics like *U* or *Q*95.

To jointly explore the power and specificity of all these statistics, we generated receiving operating characteristic (ROC) curves under various selection and admixture regimes (Figures 3 and S7). In general, *Q*95_*A,B,C*_ (1%, 100%), *Q*95_*A,B,C*_ (10%, 100%) and *f_D_* are very powerful statistics for detecting AI under strong (s=0.1) and intermediate (s=0.01) selection pressures. The number of uniquely shared sites *U_A,B,C_*(*w, x, y*) is also powerful, so long as the population in the target panel (*B*) is large. Additionally, for different choices of x, using *w* = 1% yields a more powerful statistic than using *w* = 10%. We also tested AI scenarios with weak selection (s=0.001), in which all statistics performed rather poorly, with *Q*95 and *f_D_* performing comparably better than the rest (Figure S8). However, under these conditions, it is very unlikely that the selected allele will reach appreciable frequencies (Figure S2), so the lack of sensitivity of all statistics here is largely a reflection of the fact that in most simulations the selected allele is not successful, especially when the probability of admixture is low. Conditioning on survival of the allele should therefore increase sensitivity.

**Figure 3:**
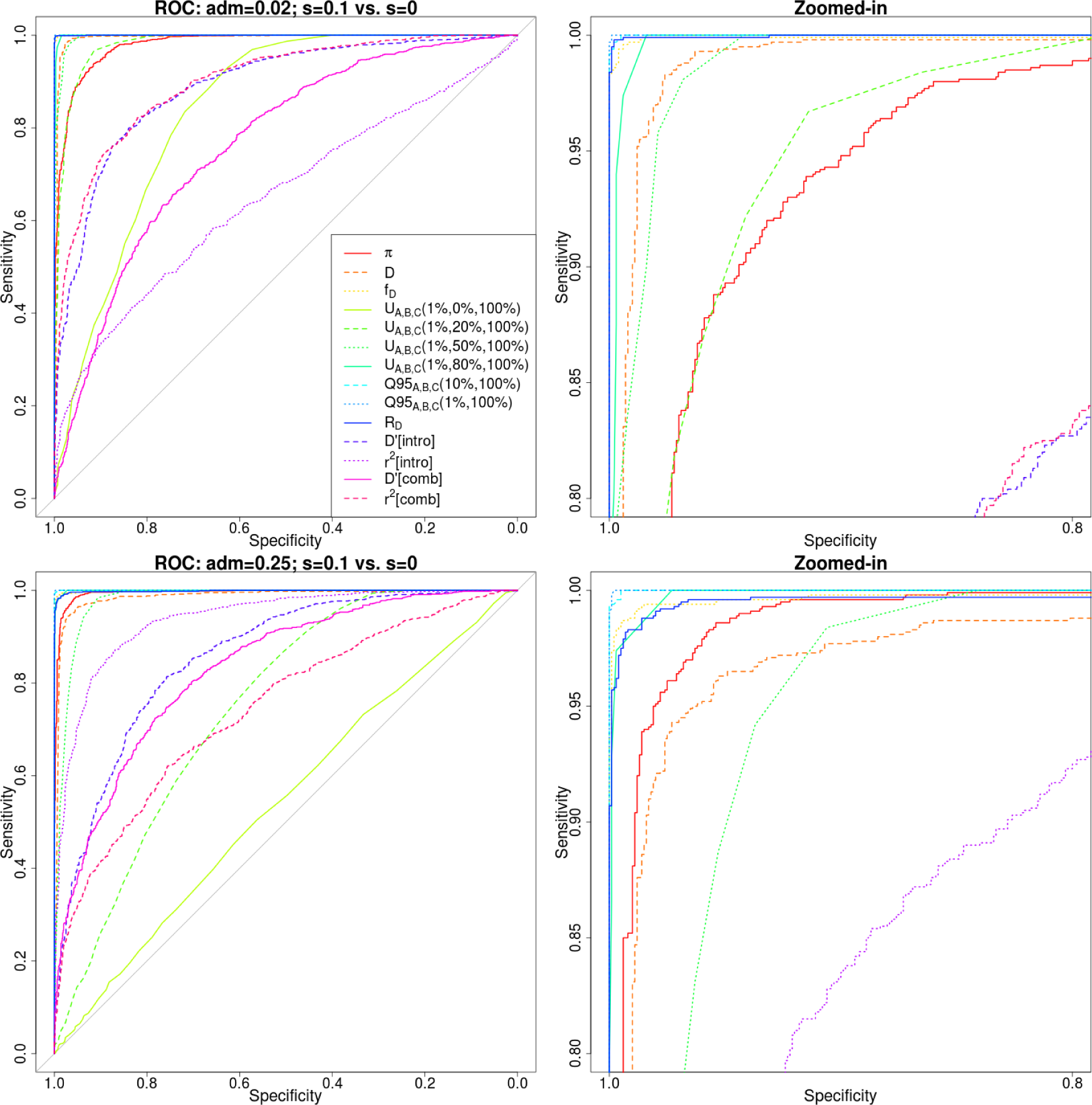
Receiver operating characteristic curves for a scenario of adaptive introgression (s=0.1) compared to a scenario of neutrality (s=0), using 1,000 simulations for each case. Populations A and B split from each other 4,000 generations ago, and their ancestral population split from population C 16,000 generations ago. Population sizes were constant and set at 2*N* = 20, 000. The admixture event occurred 1,600 generations ago from population C to population B, at rate 2% (top panels) or 25% (bottom panels). The right panels are zoomed-in versions of the left panels.

We were also interested in the joint distribution of pairs of these statistics. Figure S9 shows the joint distribution of *Q*95_*A,B,C*_ (1%, 100%) in the y-axis and four other statistics (*R_D_*, *π*, *D* and *f_D_*) in the x-axis, under different admixture and selection regimes. One can observe, for example, that while *Q*95_*A,B,C*_ (1%, 100%) increases with increasing selection intensity and admixture rates, *π* increases with increasing admixture rates, but decreases with increasing selection intensity. Thus, under AI the two forces cancel each other out, and we obtain a similar value of *π* as under neutrality. Furthermore, the joint distributions of *Q*95_*A,B,C*_ (1%, 100%) and *f_D_* or *R_D_* show particularly good separation among the different AI scenarios.

Another joint distribution that is especially good at separating different AI regimes is the combination of *Q*95_*A,B,C*_ (*w*, 100%) and *U_A,B,C_*(*w, x,* 100%). In Figure 4, we show this joint distribution, for different choices of w (1%, 10%) and x (20%, 50%). Here, with increasing intensity of selection and admixture, the number of uniquely shared sites and the quantile statistic increase, but the quantile statistic tends to only reach high values when selection is strong, even if admixture rates are low.

**Figure 4:**
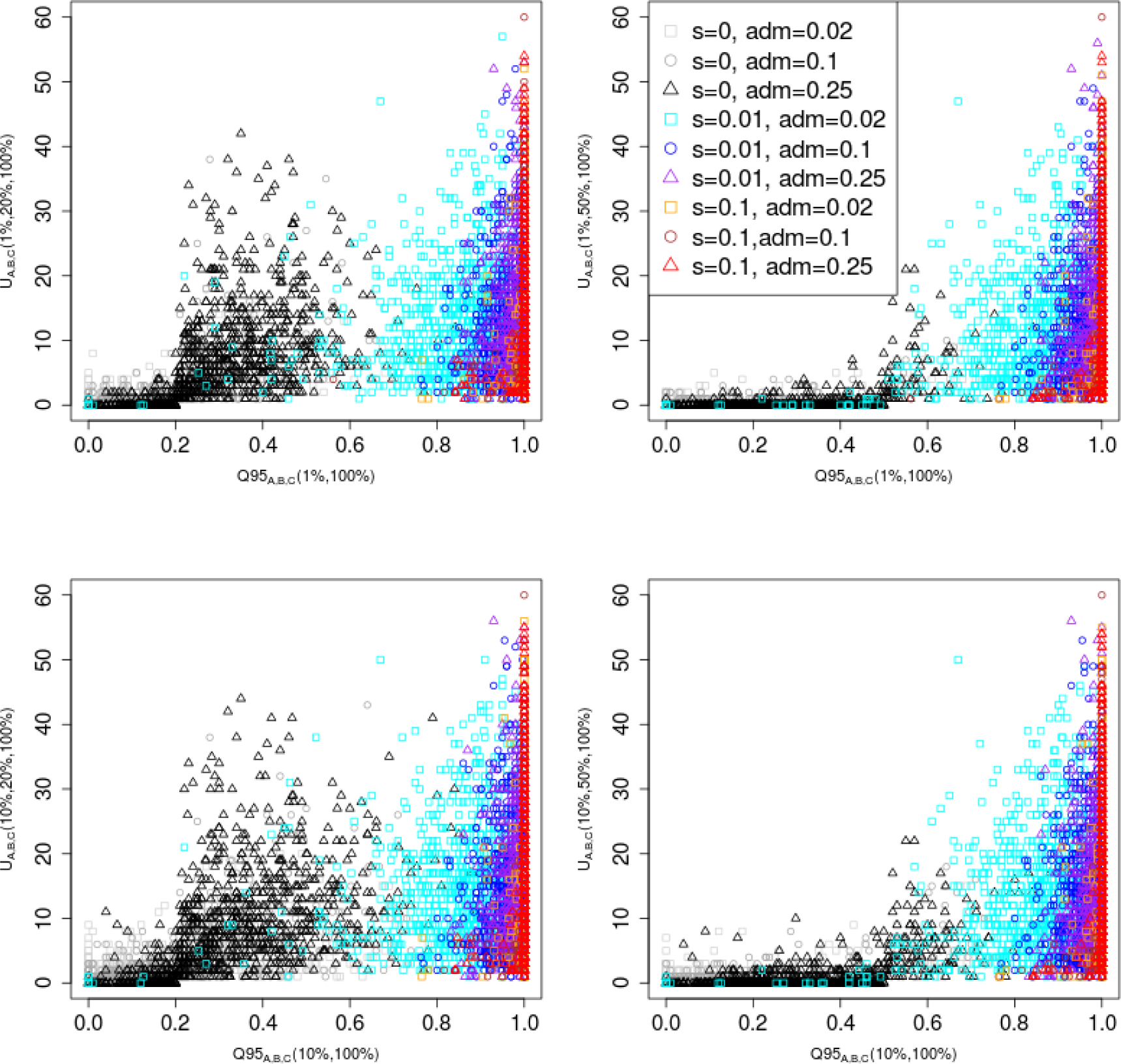
Joint distribution of *Q*95_*A,B,C*_ (*w, y*) and *U_A,B,C_*(*w, x, y*) for different choices of w (1%, 10%) and x (20%, 50%). We set y to 100% in all cases. 100 individuals were sampled from panel A, 100 from panel B and 2 from panel C. The demographic parameters were the same as in Figure 3.

### 2.3. Alternative demographic scenarios

We evaluated the performance of our statistics under various alternative demographic scenarios. First, we simulated a 5X bottleneck occurring in population B 1,600 generations before the admixture event, and lasting 200 generations, to observe its effects on the power of the statistics for detecting AI (Figure 2.B). Though we observe a reduction in power - most evident in the heterozygosity statistics - none of the statistics are very strongly affected by this event (Figure S10). We also simulated a bottleneck of equal size but occurring after the admixture event - starting 1,400 generations ago, and lasting 200 generations (Figure 2.C). In this case, the sensitivity of all the statistics is strongly reduced when the admixture rate is low (Figure S11). For example, when looking at the raw values of the *UA,B,C* and *Q*95*A,B,C* statistics, we observe that for low admixture rates the distribution under selection has a larger overlap with the distribution under neutrality, which explains the low power (Figures S12, S13). Additionally, *UA,B,C* (but not *Q*95) seems to display more elevated values under neutrality in the bottleneck model than in the constant population size model. However, the relative performance of each statistic with respect to all the others does not appear to change substantially (Figure S11).

We next explored a model where the introgressed haplotype was not immediately adaptive in the Eurasian population, but instead underwent an intermediate period of neutral drift, before it becomes advantageous (Figure 2.D). In such a situation, our power to detect AI is reduced, for all statistics (Figure S14). This is particularly an issue when the admixture rate is low, as in those cases the starting frequency of the selected allele in the Eurasian population is low, so it is more likely to drift to extinction during the neutral period, before it can become advantageous.

We also evaluated the performance of our statistics under selective scenarios that did not involve adaptive introgression, to check which of them were sensitive to these models and which were not. Under a model of selection from de novo mutation (SDN, Figure 2.E) - in which a single mutation appears in the receiving population after the split time between it and the non-admixed population the heterozygosity (*π*) and linkage disequilbrium statistics (*r*^2^[*intro*] and *D*′[*intro*]) are the most sensitive ones (Figure S15). This is expected, given that classical selective sweeps are known to strongly affect patterns of heterozygosity and linkage disequilibrium in the neighborhood of the selected site [22, 23, 24]. Since all other statistics have very poor sensitivity to detect SDN, we expect to be able to distinguish signatures generated from SDN and AI. One caveat to this is the scenario in which a *de novo* selected mutation occurs on an introgressed haplotype immediately after an introgression event - before the haplotype has a chance to expand and recombine in the population - in which case our statistics will not be able to distinguish SDN from AI.

We also simulated a model of selection from standing variation (Figure 2.F), by randomly selecting 20% of haplotypes within the introgressed population to be advantageous, after the split time between it and the non-introgressed population. In this case, all statistics perform poorly, especially when admixture is low. Interestingly, when admixture is high (Figure S16), *Q*95_*A,B,C*_ (1%, 100%) and *U_A,B,C_* (1%, 0%, 100%) are the best performing statistics. This is likely because some of the haplotypes that are randomly chosen to be selected also happen to be ancestrally polymorphic and present in the archaic humans.

When we set ancestral structure to be our null model, we observe different behaviors depending on the strength of the migration rates. When the migration rates are strong (Figure S17), we have excellent power to detect AI with several statistics, including *Q*95_*A,B,C*_ (1%, 100%), *D*, *f_D_*, *R_D_* and *U_A,B,C_* (1%, 50%, 100%). When the rates are of medium strength (Figure S18), the power is slightly reduced, but the same statistics are the ones that perform best. When the migration rates are weak - meaning ancestral structure is very strong - *Q*95_*A,B,C*_ (1%, 100%) loses power, and the best-performing statistics are *R_D_*, *D* and *f_D_* (Figure S19). We note, though, that the genome-wide *D* observed under this last ancestral structure model (*D* = 0.24) is much more extreme than the genome-wide D observed empirically between any Eurasian population and Neanderthals or Denisovans, suggesting that if there was ancestral structure between archaic and modern humans, it was likely not of this magnitude.

### 2.4. Global features of uniquely shared archaic alleles

Before identifying candidate genes for adaptive introgression, we investigated the frequency and number of uniquely shared sites at the genome-wide level. Specifically, we wanted to know whether human populations varied in the number of sites with uniquely shared archaic alleles, and whether they also varied in the frequency distribution of these alleles. Therefore, we computed *UA,B,Nea,Den*(1%,x,y,z) and *Q*95*A,B,Nea,Den*(1%,y,z) for different choices of x, y and z. We used different cutoffs for the frequency of the archaic allele (*x*) in the target population B: 0%, 20% and 50%. To look for alleles uniquely shared with the Altai Neanderthal genome only, we set *y* = 100% and *z* = 0%. To look for alleles uniquely shared with the Denisovan genome only, we set *y* = 0% and *z* = 100%. Finally, to look for uniquely shared alleles matching both of the archaic genomes, we set *y* = 100% and *z* = 100%.

We used each of the non-African panels in the 1000 Genomes Project phase 3 data [20] as the “target” panel (B), and chose the outgroup panel (A) to be the combination of all African populations (YRI, LWK, GWD, MSL, ESN), excluding admixed African-Americans. We note that this is a conservative reference panel, as some of the African panels - like LWK - are from populations with a substantial amount of Eurasian ancestry [20], likely preventing the detection of introgressed segments at some loci.

When setting *x* = 0% (i.e. not imposing a frequency cutoff in the target panel B), South Asians as a target population show the largest number of archaic alleles (Figure 5.A). However, East Asians have a larger number of high-frequency uniquely shared archaic alleles than Europeans and South Asians, for both *x* = 20% and *x* = 50% (Figure 5.B-C). Population-specific D-statistics (using YRI as the non-admixed population) also follow this trend (Figure S20) and we observe this pattern when looking only at the X chromosome as well (Figure S21). These results hold in comparisons with both archaic human genomes, but we observe a stronger signal when looking at Neanderthal-specific shared alleles. To correct for the fact that some panels have more segregating sites than others (and may therefore have more archaic-like segregating sites), we also scaled the number of uniquely shared sites by the total number of segregating sites per population panel (Figure 5.D-F), and we see in general the same patterns, with the exception of a Peruvian panel, which we discuss further below. We also observe similar patterns when calculating *Q*95_*A,B,Nea,Den*_(1%, *y*, *z*) genome-wide (Figure S22). The elevation in *U_A,B,Nea,Den_* and *Q*95_*A,B,Nea,Den*_ in East Asians may possibly result from higher levels of archaic ancestry in East Asians than in Europeans [25], which some studies argue could be due to additional admixture events occurring in East Asians [26, 27].

**Figure 5:**
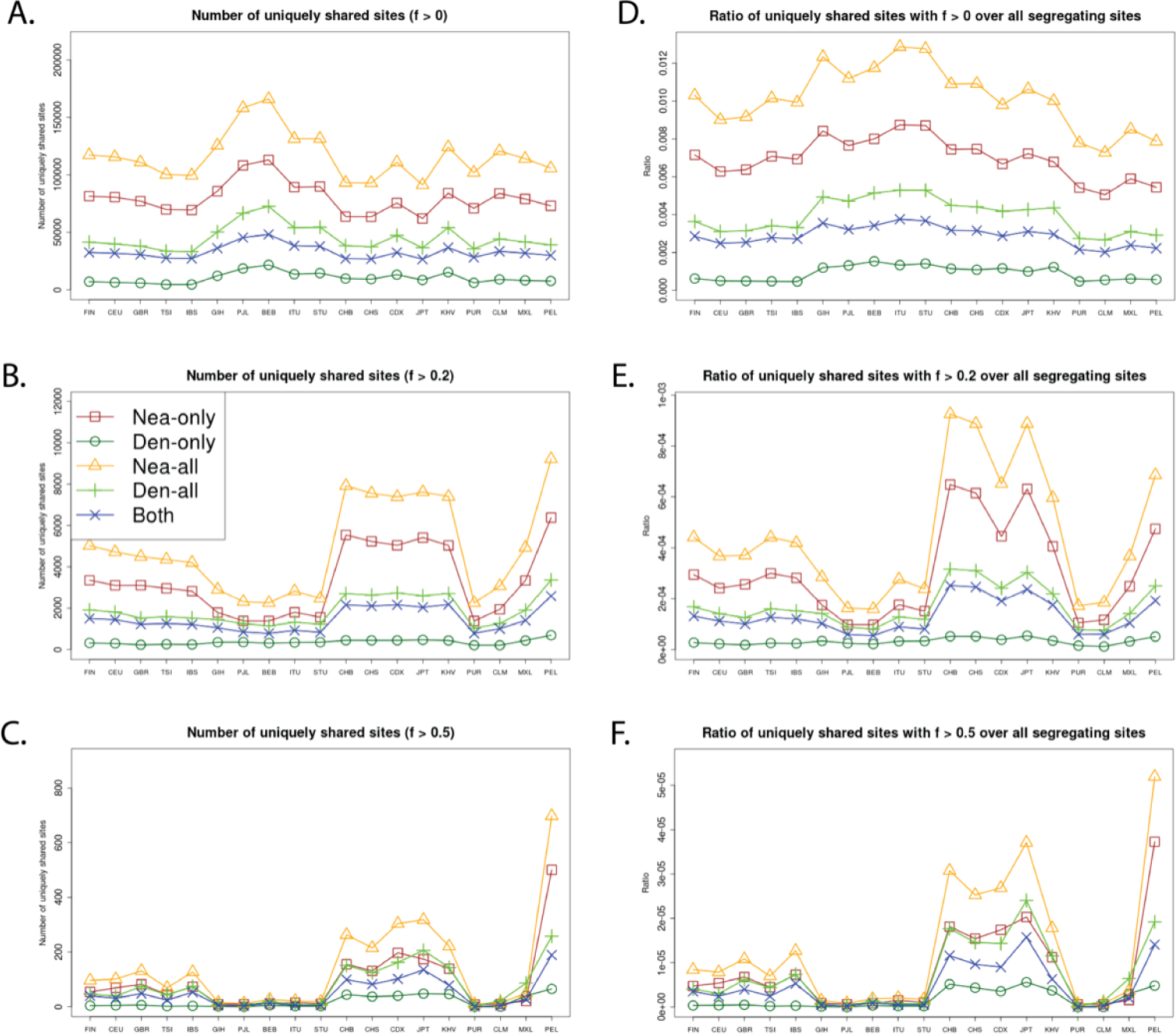
We computed the number of uniquely shared sites in the autosomes and the X chromosome between particular archaic humans and different choices of present-day non-African panels X (x-axis) from phase 3 of the 1000 Genomes Project. We used a shared frequency cutoff of 0% (A), 20% (B) and 50% (C). Nea-only = *U_Af r,X,Nea,Den_*(1%, 20%, 100%, 0%). Den-only = *U_Af r,X,Nea,Den_*(1%, 20%, 0%, 100%). Nea-all = *U_Af r,X,Nea_*(1%, 20%, 100%). Den-all = *U_Af r,X,Den_*(1%, 20%, 100%). Both = *U_Af r,X,Nea,Den_*(1%, 20%, 100%, 100%). Finally, we scaled each of the statistics from panels A-C by the number of segregating sites in each 1000 Genomes population panel, yielding panels D-F.

Surprisingly, the Peruvians (PEL) harbor the largest amount of high frequency mutations of archaic origin than any other single population, especially when using Neanderthals as bait (Figures 5.B-C,S21). It is unclear whether this signal is due to increased drift or selection in this population. Skoglund et al. [28] argue via simulations that if one analyzes a population with high amounts of recent genetic drift and excludes SNPs where the minor allele is at low frequency, some statistics that are meant to detect archaic ancestry - like D - may be artificially inflated. Our filtering procedure to select uniquely shared archaic alleles necessarily excludes sites where the archaic allele is at low frequency in the target panel, and the PEL panel comes from a population with a history of low effective population sizes (high drift) relative to other Non-Africans [20], which could explain this pattern. This could also explain why the effect is not seen when x = 0% (Figure 5.A), or when computing D-statistics (Figure S20), both of which include sites with low-frequency alleles in their computation. Additionally, scaling the uniquely shared sites by the total number of segregating sites per population panel mitigates (but does not completely erase) this pattern. After scaling, PEL shows levels of archaic allele sharing within the range of the East Asian populations at *x* = 20% (Figure 5.E), but is still the panel with the largest number of archaic sites at *x* = 50% (Figure 5.F).

Furthermore, we plotted the values of *U_AF R,X,Nea,Den_*(1%, *x, y, z*) and *Q*95_*AF R,X,Nea,Den*_(1%, *y, z*) jointly for each population X, under different frequency cutoffs *x*. When *x* = 0%, there is a generally inversely proportional relationship between the two scores (Figure S23), but this becomes a directly proportional relationship when *x* = 20% (Figure 6) or *x* = 50% (Figure S27). Here, we also clearly observe that PEL is an extreme panel with respect to both the number and frequency of archaic shared derived alleles, and that East Asian and American populations have more high-frequency archaic shared alleles than Europeans.

**Figure 6:**
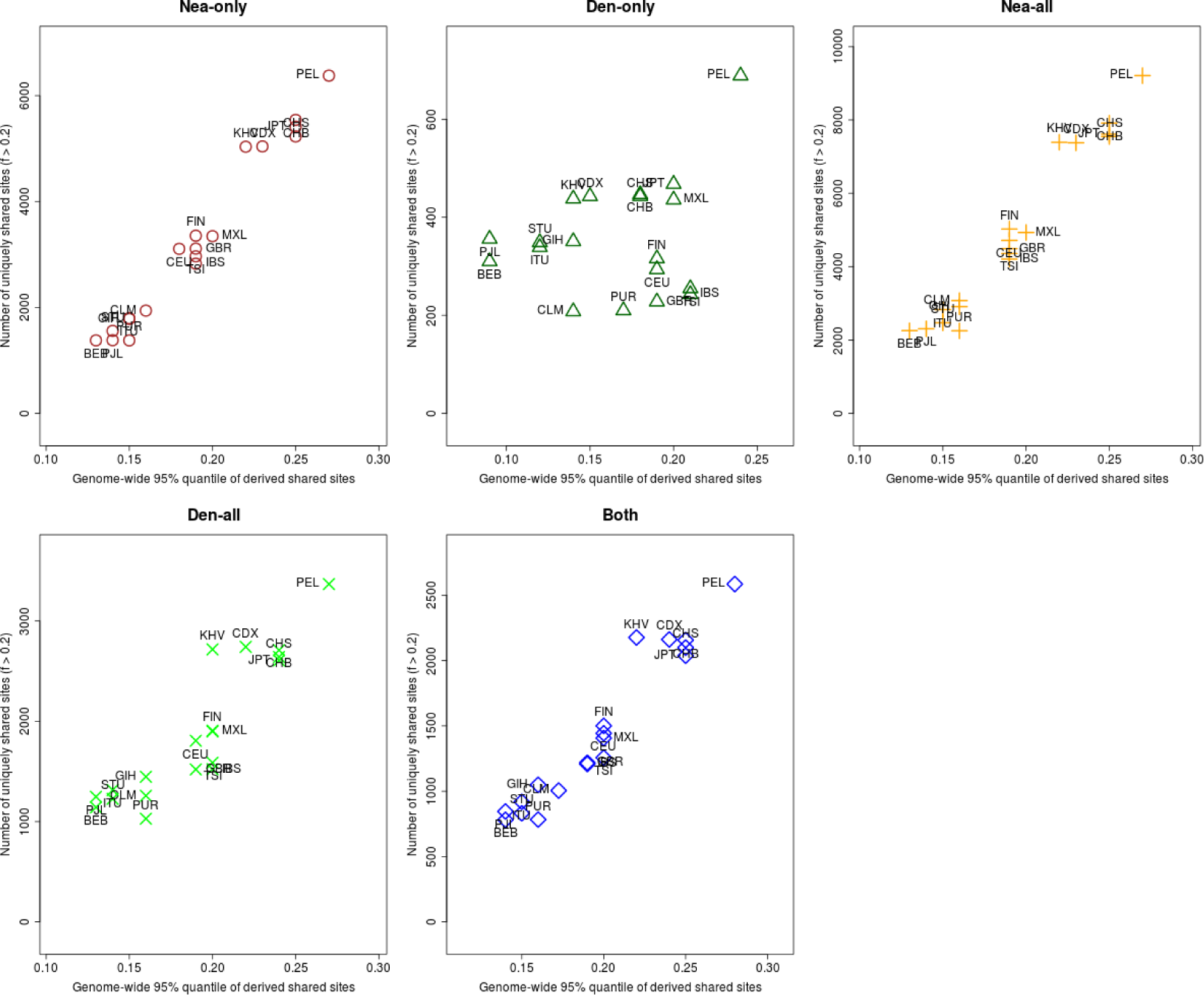
For each population panel from the 1000 Genomes Project, we jointly plotted the *U* and *Q*95 statistics with an archaic frequency cutoff of > 20% within each population. Nea-only = *U_Af r,X,Nea,Den_*(1%, 20%, 100%, 0%) and *Q*95_*Af r,X,Nea,Den*_(1%, 100%, 0%). Den-only = *U_Af r,X,Nea,Den_*(1%, 20%, 0%, 100%) and *Q*95_*Af r,X,Nea,Den*_ (1%, 0%, 100%). Nea-all = *U_Af r,X,Nea_* (1%, 20%, 100%) and *Q*95_*Af r,X,Nea*_ (1%, 100%). Den-all = *U_Af r,X,Den_* (1%, 20%, 100%) and *Q*95_*Af r,X,Den*_ (1%, 100%). Both = *U_Af r,X,Nea,Den_* (1%, 20%, 100%, 100%) and *Q*95_*Af r,X,Nea,Den*_ (1%, 100%, 100%).

We checked via simulations if the observed excess of high frequency archaic derived mutations in Americans and especially Peruvians could be caused by genetic drift, as a consequence of the bottleneck that occurred in the ancestors of Native Americans as they crossed Beringia. We observe that if the introgressed population B undergoes a bottleneck, this can lead to a larger number of *U_A,B,C_*(*w, x, y, z*) for large values of x (Figures S12,S13,S24). Indeed, population structure analyses of the 1000 Genomes samples suggest that Peruvians have the largest amount of Native American ancestry [20] and show a bottleneck with a lack of recent population growth, which could explain this pattern. We also observe an increase in the variance of the distribution of *U* and *Q*95 in the presence of a bottleneck, especially when long and severe (Figures S25, S26).

### 2.5. Candidate regions for adaptive introgression

To identify adaptively introgressed regions of the genome, we computed *U_A,B,C,D_*(*w, x, y, z*) and *Q*95_*A,B,C,D*_(*w, y, z*) in 40 kb non-overlapping windows along the genome, using the low-coverage sequencing data from phase 3 of the 1000 Genomes Project [20]. We used this window size because the mean length of introgressed haplotypes in ref. [2] was 44,078 bp (Supplementary Information 13). Our motivation was to find regions under AI in a particular panel B, using panel A as a non-introgressed outgroup (generally Africans, unless otherwise stated). We used the high-coverage Altai Neanderthal genome [2] as bait panel C and the high-coverage Denisova genome [4] as bait panel D. We deployed these statistics in three ways: a) to look for Neanderthal-specific AI, we set *y* = 100% and *z* = 0%; b) to look for Denisova-specific AI, we set *y* = 0% and *z* = 100%; c) to look for AI matching both of the archaic genomes, we set *y* = 100% and *z* = 100% (Figure S1, Table S3). To try to determine the adaptive pressure behind the putative AI event, we obtained all the CCDS-verified genes located inside each window [29].

For guidance as to how high a value of *U* and *Q*95 we would expect under neutrality, we used the simulations from Figure 2 to obtain 95% empirical quantiles of the distribution of these scores under neutrality. Tables S1 and S2 show the 95% quantiles for these two statistics under various models of adaptive introgression and ancestral structure, for different choices of parameter values (see Methods Section). When examining our candidates for AI below, we focused on windows whose values for *U_AB,Nea,Den_*(*w, x, y, z*) and *Q*95_*A,B,Nea,Den*_(*w, y, z*) were both in the 99.9% quantile of their respective genome-wide distributions. We verified via simulations that, under a simple model of neutral admixture at a genome-wide rate of 2%, the estimated probability of obtaining values as high as these (or the false positive rate, FPR) was between 10.6% and 0%, depending on the target population chosen. The highest rates correspond to the African-American (ASW) admixed panel, as this panel contains high proportions of ancestry from the outgroup panel (unadmixed Africans) and are therefore not well-suited for our statistics. Excluding ASW, the highest estimated FPR was 5.5%.

We also calculated *D* and *f_D_* along the same windows (using Africans as the non-admixed population), and saw good agreement with the new statistics presented here (Table S3). Finally, we further validated the regions most likely to have been adaptively introgressed by searching for archaic tracts of introgression within them that were at high frequency, using a Hidden Markov Model (see below).

#### 2.5.1. Continental populations

When focusing on adaptive introgression in continental populations, we first looked for uniquely shared archaic alleles specific to Europeans that were absent or almost absent (< 1% frequency) in Africans and East Asians. Conversely, we also looked for uniquely shared archaic alleles in East Asians, which were absent or almost absent in Africans and Europeans. In this continental survey, we ignored Latin American populations as they have high amounts of European and African ancestry, which could confound our analyses. Figure 7 shows the number of sites with uniquely shared alleles for increasing frequency cutoffs in the introgressed population, and for different types of archaic alleles (Neanderthal-specific, Denisova-specific or common to both archaic humans). In other words, we calculated *U_AF R,EU R,Nea,Den_*(1%, *x*, *y*, *z*) and *U_AF R,EAS,Nea,Den_*(1%, *x, y, z*) for different values of x (0%, 20%, 50% and 80%) and different choices of y and z, depending on which type of archaic alleles we were looking for. We observe that the regions in the extreme of the distributions for *x* = 50% corresponded very well to genes that had been previously found to be candidates for adaptive introgression from archaic humans in these populations, using more complex probabilistic methods [6, 5] or gene-centric approaches [30]. These include *BNC2* (involved in skin pigmentation [31, 32]), *POU2F3* (involved in skin keratinocyte differentiation [33, 34]), *HYAL2* (involved in the response to UV radiation on human keratinocytes [35]), *SIPA1L2* (involved in neuronal signaling [36]) and *CHMP1A* (a regulator of cerebellar development [37]). To be more rigorous in our search for adaptive introgression, we looked at the joint distribution of the *U* statistic and the *Q*95 statistic for the same choices of w, y and z, and then selected the regions that were in the 99.9% quantiles of the distributions of both statistics (Figures 8, S28, S29). We find that the strongest candidates here are *BNC2*, *POU2F3*, *SIPA1L2* and the *HYAL2* region.

**Figure 7:**
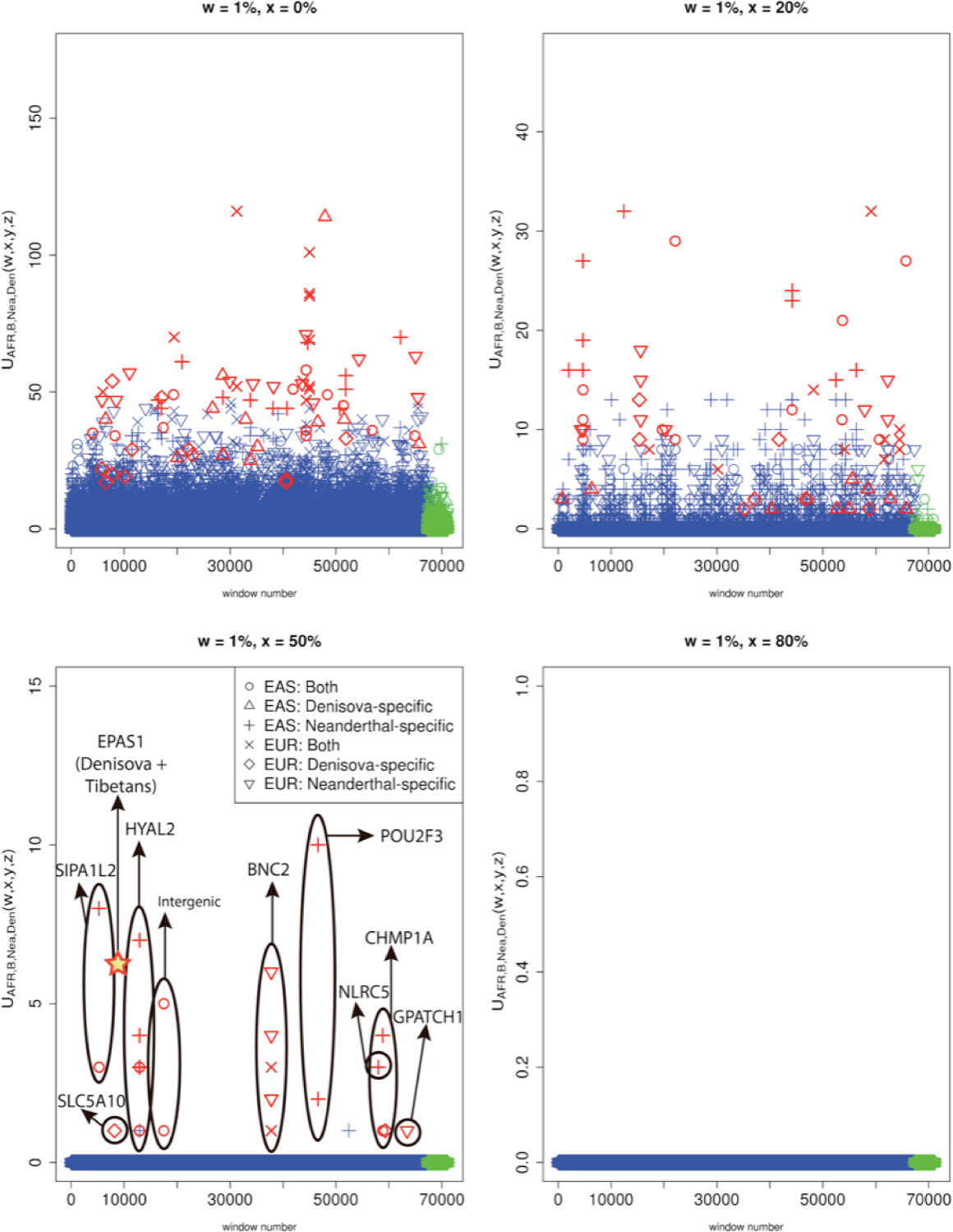
We partitioned the genome into non-overlapping windows of 40kb. Within each window, we computed *U_EU R,Out,Nea,Den_* (1%, *x, y, z*) and *UEAS,Out,Nea,Den* (1%, *x, y, z*), where Out=EAS+AFR for EUR as the target introgressed population, and Out=EUR+AFR for EAS as the target introgressed population. We searched for Neanderthal-specific alleles (*y* = 100%, *z* = 0%), Denisovan-specific alleles (*y* = 0%, *z* = 100%) and alleles present in both archaic genomes (*y* = 100%, *z* = 100%) that were uniquely shared with either EUR or EAS at frequencies above different cutoffs (x=0%, x=20%, x=50% and x=80%). Windows that fall within the upper tail of the distribution for each modern-archaic population pair are colored in red (P < 0.001 / number of pairs tested) and those that do not are colored in blue, except for those in the X chromosome, which are in green. Ovals drawn around multiple points contain multiple windows with uniquely shared alleles that are contiguous. For comparison, the number of high frequency uniquely shared sites between Denisova and Tibetans is also shown [18], although Tibetans are not included in the 1000 Genomes data and the region is 32 kb long, so this may be an underestimate.

We also scanned for regions of the genome where South Asians (SAS) had uniquely shared archaic alleles at high frequency, which were absent or almost absent in Europeans, East Asians and Africans. In this case, we focused on *x* = 20% because we found that *x* = 50% left us with no candidate regions. Among the candidate regions sharing a large number of high-frequency Neanderthal alleles in South Asians, we find genes *ASTN2*, *SFMBT1*, *MUSTN1* and *MAML2* (Figure S30). *ASTN2* is involved in neuronal migration [38] and is associated with schizophrenia [39, 40]. *SFMBT1* is involved in myogenesis [41] and is associated with hydrocephalus [42]. *MUSTN1* plays a role in the regeneration of the muscoskeletal system [43]. Finally, *MAML2* codes for a signaling protein [44, 45], and is associated with cutaneous carcinoma [46] and lacrimal gland cancer [47].

#### 2.5.2. Eurasia

We then looked for AI in all Eurasians (EUA=EUR+SAS+EAS, ignoring American populations) using Africans as the non-admixed population (AFR, ignoring admixed African-Americans). Figure 8 shows the extreme outlier regions that are in the 99.9% quantiles for both *UEU_A,AFR,Nea,Den_*(1%, 20%, *y, z*) and *Q*95_*EU A,AF R,Nea,Den*_(1%, *y, z*), while Figure S31 shows the entire distribution. We focused on *x* = 20% because we found that *x* = 50% left us with almost no candidate regions. In this case, the region with by far the largest number of uniquely shared archaic alleles is the one containing genes *OAS1* and *OAS3*, involved in innate immunity [48, 49, 50, 51]. This region was previously identified as a candidate for AI from Neanderthals in non-Africans [52]. Another region that we recover and was previously identified as a candidate for AI is the one containing genes *TLR1* and *TLR6* [53, 54]. These genes are also involved in innate immunity and have been shown to be under positive selection in some non-African populations [55, 56].

**Figure 8:**
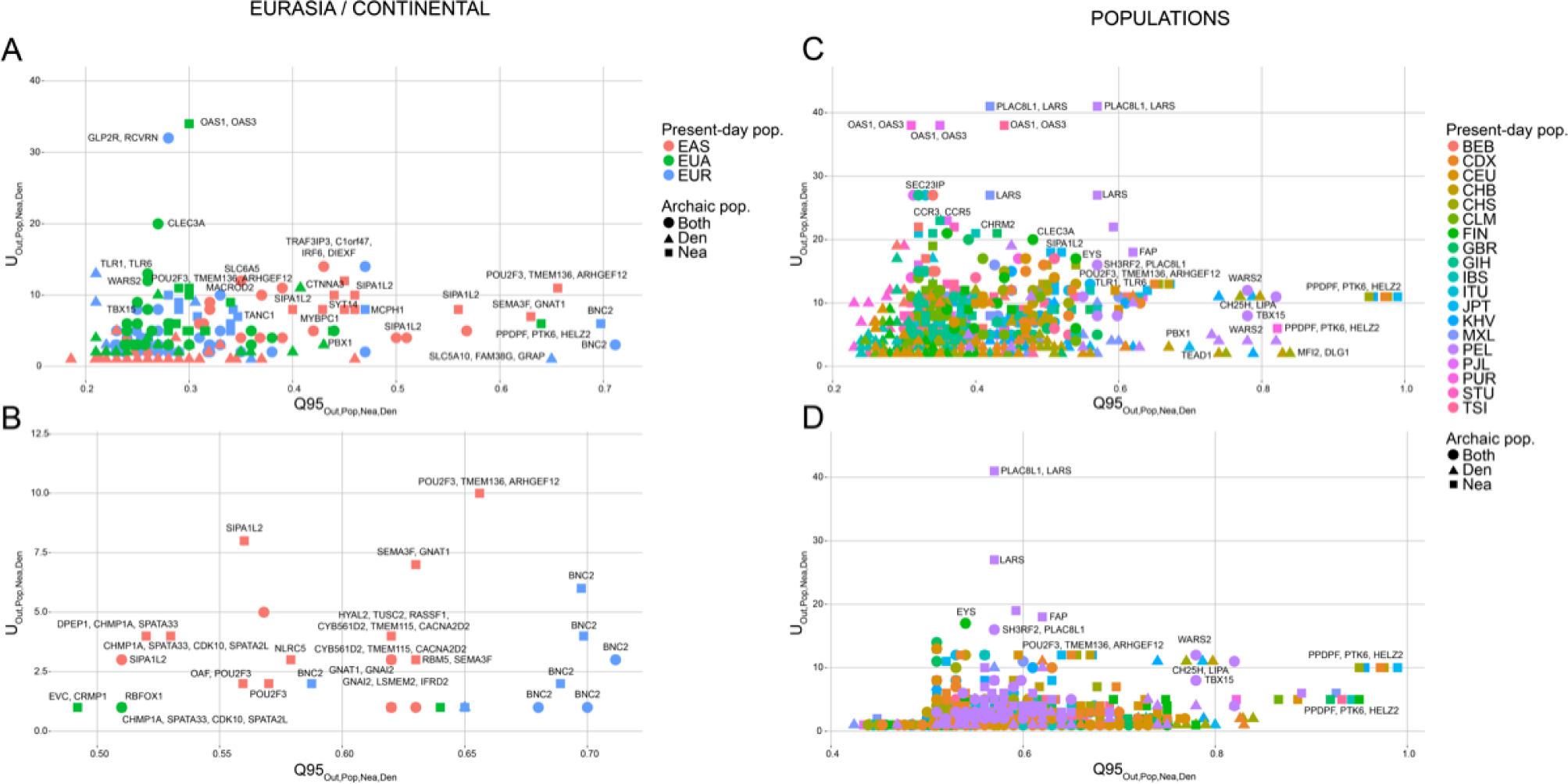
We plotted the 40kb regions in the 99.9% highest quantiles of both the *Q*95_*Out,P op,Nea,Den*_(1%, *y, z*) and *U_Out,P op,Nea,Den_*(1%, *x, y, z*) statistics for different choices of target introgressed population (Pop) and outgroup non-introgressed population (Out), and different archaic allele frequency cutoffs within the target population (x). A) We plotted the extreme regions for continental populations EUR (Out=EAS+AFR), EAS (Out=EUR+AFR) and Eurasians (EUA, Out=AFR), using a target population archaic allele frequency cutoff x of 20%. B) We plotted the extreme regions from the same statistics as in panel A, but with a more stringent target population archaic allele frequency cutoff x of 50%. C) We plotted the extreme regions for individual non-African populations within the 1000 Genomes data, using all African populations (excluding African-Americans) as the outgroup, and a cutoff x of 20%. D) We plotted the extreme regions from the same statistics as in panel C, but with a more stringent target population archaic allele frequency cutoff x of 50%. Nea-only = *U_Out,P op,Nea,Den_*(1%, *x*, 100%, 0%) and *Q*95_*Out,P op,Nea,Den*_ (1%, 100%, 0%). Den-only = *U_Out,P op,Nea,Den_*(1%, *x*, 0%, 100%) and *Q*95_*Out,P op,Nea,Den*_ (1%, 0%, 100%). Both = *U_Out,P op,Nea,Den_* (1%, *x*, 100%, 100%) and *Q*95_*Out,P op,Nea,Den*_ (1%, 100%, 100%).

Interestingly, we find that a very strong candidate region in Eurasia contains genes *TBX15* and *WARS2*. This region has been associated with a variety of traits, including adipose tissue differentiation [57], body fat distribution [58, 59, 60, 61], hair pigmentation [62], facial morphology [63, 64], ear morphology [65], stature [64] and skeletal development [66, 64]. It was previously identified as being under positive selection in Greenlanders [67], and it shows particularly striking signatures of adaptive introgression, so we devote a separate study to its analysis [68].

#### 2.5.3. Population-specific signals of adaptive introgression

To identify population-specific signals of AI, we looked for archaic alleles at high frequency in a particular non-African panel X, which were also at less than 1% frequency in all other non-African and African panels, excluding X (Table S3). This is a very restrictive requirement, and indeed, we only find a few windows in a single panel (PEL) with archaic alleles at more than 20% frequency. One of the regions with the largest number of uniquely shared Neanderthal sites in PEL contains gene *CHD2*, which codes for a DNA helicase [69] involved in myogenesis (UniProtKB by similarity), and that is associated with epilepsy [70, 71]. We note, however, that the presence of these extreme regions could be due to the global elevation in the U statistic caused by higher levels of drift in Peruvians, as explained above.

#### 2.5.4. Shared signals among populations

In the previous section, we focused on regions where archaic alleles were uniquely at high frequencies in particular populations, but at low frequencies in all other populations. This precludes us from detecting AI regions that are shared across more than one non-African population. To address this, we conditioned on observing the archaic allele at less than 1% frequency in a non-admixed outgroup panel composed of all the African panels (YRI, LWK, GWD, MSL, ESN), excluding African-Americans, and then looked for archaic alleles at high frequency in particular non-African populations. Unlike the previous section, we did not condition on the archaic allele being at low frequency in other non-African populations as well. The whole joint distributions of *U* and *Q*95 for this choice of parameters for each non-African panel are shown in figs. S32 to S50, while regions in the 99.9% quantile for both statistics are shown in Figure 8.

Here, we recapitulate many of the findings from our Eurasian and continental-specific analyses above, like *TLR1/TLR6*, *BNC2*, *OAS1/OAS3*, *POU2F3*, *LIPA* and *TBX15/WARS2* (Figure 8). For example, just as we found that *POU2F3* was an extreme region in the East Asian (EAS) continental panel, we separately find that almost all populations composing that panel (CHB, KHV, CHS, CDX, JPT) have archaic alleles in that region at disproportionately high frequency, relative to their frequency in Africans. Additionally though, we can learn things we would not have detected at the continental level. For example, the Bengali from Bangladesh (BEB) - a South Asian population - also have archaic alleles at very high frequencies in this region.

We detected several genes that appear to show signatures of AI across various populations (Figure 8). One of the most extreme examples is a 120 kb region containing the *LARS* gene, with 76 uniquely shared Neanderthal alleles at < 1% frequency in Africans and > 50% frequency in Peruvians, which are also at > 20% frequency in Mexicans. *LARS* codes for a leucin-tRNA synthetase [72], and is associated with liver failure syndrome [73]. Additionally, a region containing gene *ZFHX3* displays an elevated number of uniquely shared Neanderthal sites in PEL, and we also observe this when looking more broadly at East Asians (EAS) and - based on the patterns of inferred introgressed tracts (see below) - in various American (AMR) populations as well. *ZFHX3* is involved in the inhibition of estrogen receptor-mediated transcription [74] and has been associated with prostate cancer [75].

We also find several Neanderthal-specific uniquely shared sites in American panels (PEL, CLM, MXL) in a region previously identified as harboring a risk haplotype for type 2 diabetes (chr17:6880001-6960000) [76]. This is consistent with previous findings suggesting the risk haplotype was introgressed from Neanderthals and is specifically present at high frequencies in Latin Americans [76]. The region contains gene *SLC16A11*, whose expression is known to alter lipid metabolism [76]. We also find that the genes *FAP/IFIH1* have signals consistent with AI, particularly in PEL. This region has been previously associated with type 1 diabetes [77, 78]. A previous analysis of this region has suggested that the divergent haplotypes in it resulted from ancestral structure or balancing selection in Africa, followed by local episodes of positive selection in Europe, Asia and the Americas [79]. A more recent analysis has found this as a region of archaic AI in Melanesians as well [7].

Another interesting candidate region contains two genes involved in lipid metabolism: *LIPA* and *CH25H*. We find a 40 kb region with 11 uniquely shared Denisovan alleles that are at low (< 1%) frequency in Africans and at very high (> 50%) frequency in various South and East Asian populations (JPT, KHV, CHB, CHS, CDX and BEB). The Q95 and D statistics in this region are also high across all of these populations, and we also find this region to have extreme values of these statistics in our broader Eurasian scan. The *LIPA* gene codes for a lipase [80] and is associated with cholesterol ester storage disease [81] and Wolman disease [82]. In turn, the *CH25H* gene codes for a membrane hydroxylase involved in the metabolism of cholesterol [83] and associated with Alzheimer’s disease [84] and antiviral activity [85].

Finally, we find a region harboring between 3 and 10 uniquely shared Neanderthal alleles (depending on the panel used) in various non-African populations. This region was identified earlier by ref. [5] and contains genes *PPDPF*, *PTK6* and *HELZ2*. *PPDPF* codes for a probable regulator of pancreas development (UniProtKB by similarity). *PTK6* codes for an epithelial signal transducer [86] and *HELZ2* codes for a helicase that works as a transcriptional coactivator for nuclear receptors [87, 88].

### 2.6. The X chromosome

Previous studies have observed lower levels of archaic introgression in the X chromosome relative to the autosomes [5, 6]. Here, we observe a similar trend: compared to the autosomes, the X chromosome contains a smaller number of windows with sites that are uniquely shared with archaic humans (Figure 7). For example, for *w* = 1% and *x* = 20%, we observe that, in Europeans, 0.4% of all windows in the autosomes have at least one uniquely shared site with Neanderthals or Denisovans, while only 0.05% of all windows in the X chromosome have at least one uniquely shared site (P = 4.985 × 10^−4^, chi-squared test assuming independence between windows). The same pattern is observed in East Asians (P = 1.852 × 10^−8^).

Nevertheless, we do identify some regions in the X chromosome exhibiting high values for both *U_A,B,C,D_*(*w, x, y, z*) and *Q*95_*A,B,C,D*_(*w, y, z*). For example, a region containing gene DHRSX contains a uniquely shared site where a Neanderthal allele is at < 1% frequency in Africans, but at > 50% frequency in a British panel (GBR). The site is also at high frequency (29% − 47%) in the other European panels, but never as high as in GBR (55%). It is also surrounded by 5 neighboring SNPs that have intermediate Neanderthal allele frequencies (24% − 41%) in GBR. Another region contains gene DMD and harbors two uniquely shared sites where two archaic (Denisovan/Neanderthal) alleles are also at low (< 1%) frequency in Africans but at > 50% frequency in Peruvians. DHRSX codes for an oxidoreductase enzyme [89], while DMD is a well-known gene because mutations in it cause muscular dystrophy [90], and was also previously identified as having signatures of archaic introgression in non-Africans [91]. We note, however, that our simulations do not account for the particular inheritance and recombination patterns of the X chromosome, so caution should be taking when calling these regions as under AI.

### 2.7. Introgressed haplotypes in candidate loci

We inspected the haplotype patterns of candidate loci with support in favor of AI. We displayed the haplotypes for selected populations at seven regions: *POU2F3* (Figure 9.A), *BNC2* (Figure 9.B), *LARS* (Figure 9.C), *FAP/IFIH1* (Figure 9.D), *OAS1* (Figure 9.E), *LIPA* (Figure 9.F) and *SLC16A11* (Figure S51.C). We included continental populations that show a large number of uniquely shared archaic alleles, and included YRI as a representative African population. We then clustered and ordered the haplotypes by similarity to the closest archaic genome (Altai Neanderthal or Denisova) (Figure 9). As can be observed, all these regions tend to show sharp distinctions between the putatively introgressed haplotypes and the non-introgressed ones. This is also evident when looking at the cumulative number of differences of each haplotype to the closest archaic haplotype, where we see a sharp rise in the number of differences, indicating strong differentiation between the two sets of haplotypes. Additionally, the YRI haplotypes tend to predominantly belong to the non-introgressed group, as expected.

**Figure 9:**
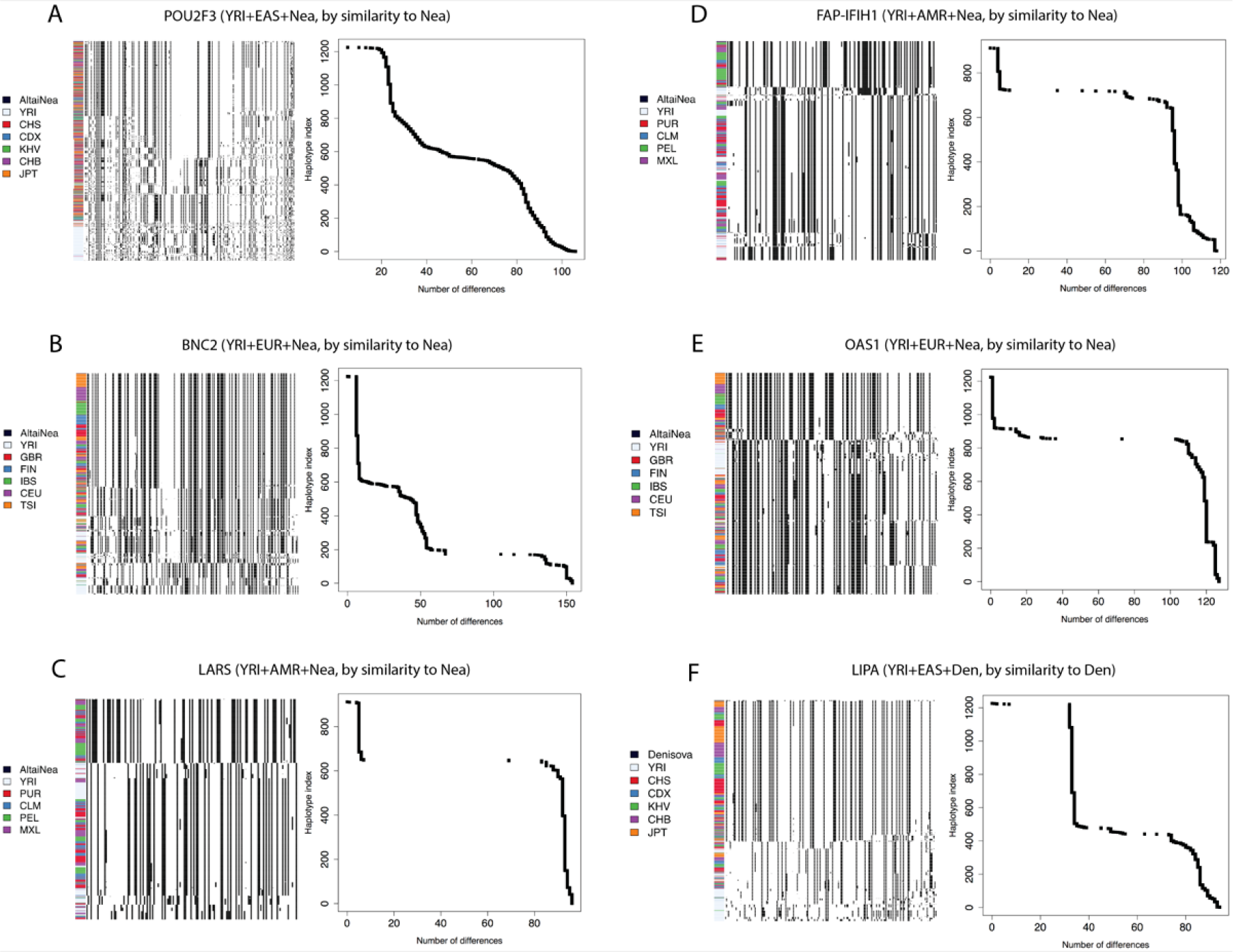
We explored the haplotype structure of six candidate regions with strong evidence for AI. For each region, we applied a clustering algorithm to the haplotypes of particular human populations and then ordered the clusters by decreasing similarity to the archaic human genome with the larger number of uniquely shared sites (see Methods Section). We also plotted the number of differences to the archaic genome for each human haplotype and sorted them simply by decreasing similarity. In the latter case, no clustering was performed, so the rows in the cumulative difference plots do not necessarily correspond to the rows in the adjacent haplotype structure plots. *POU2F3*: chr11:120120001-120200000. *BNC2*: chr9:16720001-16760000. *LARS*: chr5:145480001-145520000. *FAP/IFIH1*: chr2:163040001-163120000. *OAS1*: chr12:113360001-113400000. *LIPA*: chr10:90920001-90980000.

#### 2.7.1. Consequences of relaxing the outgroup frequency cutoff

When using a more lenient cutoff for the outgroup panel (10% maximum frequency, rather than 1%), we find a few genes that display values of the *U* statistic that are suggestive of AI, and that have been previously found to be under strong positive selection in particular human populations [92, 93]. The most striking examples are *TYRP1* in EUR (using EAS+AFR as outgroup) and *OCA2* in EAS (using EUR+AFR as outgroup)(Table S3). Both of these genes are involved in pigmentation. We caution, however, that the reason why they carry archaic alleles at high frequency may simply be because their respective selective sweeps pushed an allele that was segregating in both archaic and modern humans to high frequency in modern humans, but not necessarily via introgression.

In fact, *TYRP1* only stands out as an extreme region for the number of archaic shared alleles in EUR when using the lenient 10% cutoff, but not when using the more stringent 1% cutoff. When looking at these SNPs in more detail, we find that their allele frequency in Africans (∼ 20%) is even higher than in East Asians (∼ 1%), largely reflecting population differentiation across Eurasia due to positive selection [93], rather than adaptive introgression. When exploring the haplotype structure of this gene (Figure S51.B), we find one haplotype that shows similarities to archaic humans but is at low frequency. In the combined YRI+EUR panel, just 6.37% of all haplotypes have 36 or less differences to the Neanderthal genome, and this number is roughly the point of transition between the archaic-like and the non-archaic-like haplotypes (Figures S51.B). There is a second - more frequent - haplotype that is more distinct from archaic humans but present at high frequency in Europeans. The uniquely shared sites obtained using the lenient (< 10%) allele frequency outgroup cutoff are tagging both haplotypes together, rather than just the highly differentiated archaic-like haplotype.

*OCA2* has several sites with uniquely shared alleles in EAS (AFR+EUR as outgroup) when using the lenient 10% cutoff, but only a few (2) shared archaic sites when using the < 1% outgroup frequency cutoff. When exploring the haplotype structure of this gene, we fail to find a clear-cut differentiation between putatively introgressed and non-introgressed haplotypes, so the evidence for adaptive introgression in this region is also weak. A close inspection of its haplotype structure shows that *OCA2* does not show a large number of differences between the haplotype classes that are closer and those that are distant from the archaic humans (Figure S51.A).

Finally, using the lenient outgroup cutoff of < 10% and a target cutoff of > 20%, we find the gene with the highest number of uniquely shared sites among all the populations and cutoffs we tested: *MUC19*. This region is rather impressive in containing 115 sites where the archaic alleles are shared between the Mexican panel (MXL) and the Denisovan genome at more than 20% frequency, when using all populations that are not MXL as the outgroup. However, the actual proportion of individuals that contain a Denisova-like haplotype (though highly differentiated from the rest of present-day human haplotypes) is very small. Only 11.86% of haplotypes in the combined YRI+AMR panel show 69 differences or less to the closest archaic genome (Denisova), and the next closest haplotype has 134 differences (Figure S51.D).

Overall, a finer investigation of these three cases suggests that using a lenient outgroup frequency cutoff may lead to misleading inferences. Nevertheless, the haplotype structure of these genes and their relationship to their archaic human counterparts are quite unusual. It remains to be determined whether these patterns could be caused by either positive selection or introgression alone, or whether a combination of these or other demographic forces is required to explain them.

### 2.8. Inferred introgressed tracts

We used an HMM [21] to verify that the strongest candidate regions effectively contained archaic segments of a length that would be consistent with introgression after the population divergence between archaic and modern humans. For each region, we used the closest archaic genome (Altai Neanderthal or Denisova) as the putative source of introgression. We then plotted the inferred segments in non-African continental populations for genes with strong evidence for AI. Among these, genes with Neanderthal as the closest source (figs. S52 to S59) include: *POU2F3* (EAS,SAS), *BNC2* (EUR), *OAS1* (Eurasians), *LARS* (AMR), *FAP/IFIH1* (PEL), *CHD2* (PEL), *TLR1-6* (EAS) and *ZFHX3* (PEL). Genes with Denisova as the closest source (figs. S60 and S61) include: *LIPA* (EAS, SAS, AMR) and *MUSTN1* (SAS).

### 2.9. Testing for enrichment in genic regions

We aimed to test whether uniquely shared archaic alleles at high frequencies were enriched in genic regions of the genome. We looked at archaic alleles at high frequency in any of the Non-African panels that were also at low frequency (< 1%) in Africans. As background, we used all archaic alleles that were at any frequency larger than 0 in the same Non-African populations, and that were also at low frequency in Africans. We then tested whether the high-frequency archaic alleles tended to occur in genic regions more often than expected.

SNPs in introgressed blocks will tend to cluster together and have similar allele frequencies, which could cause a spurious en-361 richment signal. To correct for the fact that SNPs at similar allele frequencies will cluster together (as they will tend to co-occur in the same haplotypes), we performed linkage disequilibrium (LD) pruning using two methods. In one (called “LD-1”), we downloaded the approximately independent European LD blocks published in ref. [94]. For each set of high frequency derived sites, we randomly sampled one SNP from each block. In a different approach (called “LD-2”), for each set of high frequency derived sites, we subsampled SNPs such that each SNP was at least 200 kb apart from each other. We then tested these two types of LD-pruned SNP sets against 1000 SNP sets of equal length that were also LD-pruned and that were obtained randomizing frequencies and collecting SNPs in the same ways as described above.

Regardless of which LD method we used, we find no significant enrichment in genic regions for high-frequency (> 50%) Nean-369 derthal alleles (LD-1 P=352, LD-2 P=0.161) or Denisovan alleles (LD-1 P=0.348, LD-2 P=0.192). Similarly, we find no enrichment for medium-to-high-frequency (> 20%) Neanderthal alleles (LD-1 P=0.553, LD-2 P=0.874) or Denisovan alleles (LD-1 P=0.838, LD-2 P=0.44).

## 3. Discussion

Here, we carried out one of the first investigations into the joint dynamics of archaic introgression and positive selection, to develop statistics that are informative of AI. We find that one of the most powerful ways to detect AI is to look at both the number and allele frequency of mutations that are uniquely shared between the introgressed and the archaic populations. Such mutations should be abundant and at high-frequencies in the introgressed population if AI occurred. In particular, we identified two novel summaries of the data that capture this pattern quite well: the statistics *Q*95 and *U*. These statistics can recover loci under AI and are easy to compute from genomic data, as they do not require phasing.

We have also studied the general landscape of archaic alleles and their frequencies in present-day human populations. While scanning the present-day human genomes from phase 3 of the 1000 Genomes Project [20] using these and other summary statistics, we were able to recapitulate previous AI findings (like the *TLR* [53, 54] and *OAS* regions [52]) as well as identify new candidate regions for AI in Eurasia (like the *LIPA* gene and the *FAP/IFIH1* region). These mostly include genes involved in lipid metabolism, pigmentation and innate immunity, as observed in previous studies [5, 6, 95]. Phenotypic changes in these systems may have allowed archaic humans to survive in Eurasia during the Pleistocene, and may have been passed on to present-day human populations during their expansion out of Africa.

When using more lenient definitions of what we consider to be “uniquely shared archaic alleles” we find sites containing these alleles in genes that have been previously found to be under positive selection (like *OCA2* and *TYRP1*) but not necessarily under adaptive introgression. While these do not show as strong signatures of adaptive introgression as genes like *BNC2* and *POU2F3*, their curious haplotype patterns and their relationship to archaic genomes warrants further exploration.

We tested whether uniquely shared archaic alleles at high frequencies in non-Africans were significantly more likely to be found genic regions, relative to all shared archaic alleles, but did not find a significant enrichment. Though this suggests archaic haplotypes subject to AI may not be preferentially found near or inside genes, it may also be a product of a lack of power, or of the fact that not all uniquely shared archaic alleles may be truly introgressed. As mentioned before, some of these alleles may be present due to incomplete lineage sorting, which could add noise to the test signal. A more rigorous - and possibly more powerful - test could involve testing whether HMM-inferred introgressed archaic segments at high frequency tend to be found in genic regions, relative to all inferred introgressed archaic segments, while controlling for features like the length of introgressed segments and the sensitivity of the HMM to different regions of the genome. However, in this study, we did not pursue this line of research further.

In this study, we have mostly focused on positive selection for archaic alleles. One should remember, though, that a larger proportion of introgressed genetic material was likely maladaptive to modern humans, and therefore selected against. Indeed, two recent studies have shown that negative selection on archaic haplotypes may have reduced the initial proportion of archaic material present in modern humans immediately after the hybridization event(s) [96, 97].

Another caveat is that some regions of the genome display patterns that could be consistent with multiple introgression events, followed by positive selection on one or more distinct archaic haplotypes [53]. In this study, we have simply focused on models with a single pulse of admixture - followed immediately by selection or with an intermediate neutrality period in the introgressed population. We have not considered complex scenarios with multiple sources of introgression. Additionally, the currently limited availability of high-coverage archaic human genomes may prevent us from detecting AI events for which the source may not have been closely related to the sequenced Denisovan or Altai Neanderthal genomes. This may include other Neanderthal or Denisovan subpopulations, or other (as yet unsampled) archaic groups that may have lived in Africa and Eurasia.

It is also worth noting that positive selection for archaic haplotypes may be due to heterosis, rather than adaptation to particular environments [96]. That is, archaic alleles may not have been intrinsically beneficial, but simply protective against deleterious recessive modern human alleles, and therefore selected after their introduction into the modern human gene pool. The degree of dominance of deleterious alleles in humans remains elusive, so it is unclear how applicable this model would be to archaic admixture in humans.

Many of the statistics we introduced in this study have their drawbacks: notably, they depend on simulations to assess significance and some - like U - may be sensitive to local variation in mutation rates across the genome. Nevertheless they serve as useful exploratory tools, as they highlight a characteristic signature left by AI in present-day human genomes. Future avenues of research could involve developing ways to incorporate uniquely shared sites into a robust test of selection that specifically targets regions under AI. For example, one could think about modifying statistics based on local between-population population differentiation, like PBS [10], so that they are only sensitive to allele frequency differences at sites that show signatures of archaic introgression.

Finally, while this study has largely focused on human AI, several other species also show suggestive signatures of AI [98]. Assessing the extent and prevalence of AI and uniquely shared sites in other biological systems could provide new insights into their biology and evolutionary history. This may also serve to better understand how populations of organisms respond to introgression events, and to derive general principles about the interplay between admixture and natural selection.

## 4. Methods

### 4.1. Summary statistics sensitive to adaptive introgression

Several statistics have been previously deployed to detect AI events (reviewed in Racimo et al. [99]). We briefly describe these below, as well as three new statistics tailored specifically to find this signal (Table 1). One of the simplest approaches consists of applying the *D* statistic [1, 100] locally over windows of the genome. The D statistic was originally applied to compare a single human genome against another human genome, so as to detect excess shared ancestry between one of the genomes and a genome from an outgroup population. Application of this statistic comparing non-Africans and Africans served as one of the pieces of evidence in in support of Neanderthal admixture into non-Africans. However, it can also be computed from large panels of multiple individuals instead of single genomes. This form of the *D* statistic has been applied locally over windows of the genome to detect regions of excess shared ancestry between an admixed population and a source population [101, 102].

The D statistic, however, can be confounded by local patterns of diversity, as regions of low diversity may artificially inflate the statistic even when a region was not adaptively introgressed. To correct for this, Martin et al. [103] developed a similar statistic called *f_D_* which is less sensitive to differences in diversity along the genome. Both of these patterns exploit the excess relatedness between the admixed and the source population.

AI is also expected to increase linkage disequilibrium (LD), as an introgressed fragment that rises in frequency in the population will have several closely linked loci that together will be segregating at different frequencies than they were in the recipient population before admixture. Thus, two well-known statistics that are informative about the amount of LD in a region - *D*′ and *r*^2^ - could also be informative about adaptive introgression. To apply them over regions of the genome, we can take the average of each of the two statistics over all SNP pairs in a window. In the section below, we calculate these statistics in two ways: a) using the introgressed panel only (*D*′[*intro*] and *r*^2^[*intro*]), and b) using the combination of the introgressed and the non-introgressed panels (*D*′[*comb*] and *r*^2^[*comb*]). The first way (intro) should capture patterns of within-population LD in the introgressed population under AI, while the second way (comb) should capture patterns of global LD across both populations. If the introgressed population has a particular set of archaic haplotypes at high frequency that are highly differentiated from the non-archaic haplotypes in the non-introgressed panel, we expect the second way to be more powerful at distinguishing AI from neutrality.

We also introduce three new statistics that one would expect, *a priori*, to be particularly effective at identifying windows of the genome that are likely to have undergone adaptive introgression: *R_D_*, *U* and *Q*95. *R_D_* is computed by calculating - in a window of the genome - the ratio of the sequence divergence between an individual from the source population and an admixed individual, and the sequence divergence between an individual from the source population and a non-admixed individual. One can then take the average of this ratio over all individuals in the admixed and non-admixed panels. This average should be larger if the introgressed haplotype is present in a large number of individuals of the admixed population. We call this statistic *R_D_*.

Second, for a window of arbitrary size, let *U_A,B,C_*(*w, x, y*) be defined as the number of sites where a sample C (the “bait”) from an archaic source population (which could be as small as a single diploid individual) has a particular allele at frequency y, and that allele is at a frequency smaller than w in a sample A (the “outgroup”) of a population but larger than x in a sample B (the “target”) of another population (Figure 1). In other words, we are looking for sites that contain alleles shared between an archaic human genome and a test population, but absent or at very low frequencies in an outgroup (usually non-admixed) population. For example, suppose we are looking for Neanderthal adaptive introgression in the Han Chinese (CHB). In that case, we can consider CHB as our target panel, and use Africans as the outgroup panel and a single Neanderthal genome as the bait. If *U_AF R,CHB,Nea_* (1%, 20%, 100%) = 4 in a window of the genome, that means there are 4 sites in that window where the Neanderthal genome is homozygous for a particular allele and that allele is present at a frequency smaller than 1% in Africans but larger than 20% in Han Chinese. In other words, there are 4 sites that are uniquely shared at more than 20% frequency between Han Chinese and Neanderthal, but not with Africans.

This statistic can be further parametrized if we have samples from two different archaic populations (for example, a Neanderthal genome and a Denisova genome). In that case, we can define *U_A,B,C,D_*(*w, x, y, z*) as the number of sites where the archaic sample C has a particular allele at frequency y and the archaic sample D has that allele at frequency z, while the same allele is at a frequency smaller than w in an outgroup panel A and larger than x in a target panel B (Figure S1). For example, if we were interested in looking for Neanderthal-specific AI, we could set *y* = 100% and *z* = 0%, to find alleles uniquely shared with Neanderthal, but not Denisova. If we were interested in archaic alleles shared with both Neanderthal and Denisova, we could set *y* = 100% and *z* = 100%.

Another statistic that we found to be useful for finding AI events is *Q*95_*A,B,C*_ (*w, y*), and is here defined as the 95^*th*^ percentile of derived frequencies in an admixed sample B of all SNPs that have a derived allele frequency y in the archaic sample C, but where the derived allele is at a frequency smaller than w in a sample A of a non-admixed population (Figure 1). For example, *Q*95_*AFR,CHB,Nea*_ (1%, 100%) = 0.65 means that if one computes the 95% quantile of all the Han Chinese derived allele frequencies of SNPs where the Neanderthal genome is homozygous derived and the derived allele has frequency smaller than 1% in Africans, that quantile will be equal to 0.65. The motivation for this statistic is that AI will produce archaic SNPs at high frequencies in the introgressed population. The 95^*th*^ percentile should be an effective way of summarizing the frequencies of these SNPs while downweighting other SNPs that may also share the same allelic state as the archaic genomes, but that are segregating at low frequencies in the target panel and are therefore not informative about AI. In other words, it is a summary of the allele frequency spectrum in the introgressed population, conditional on only looking at alleles uniquely shared with the source population and at low frequency in the non-admixed population. As before, we can generalize this statistic if we have a sample D from a second archaic population. Then, *Q*95_*A,B,C,D*_ (*w, y, z*) is the 95^*th*^ percentile of derived frequencies in the sample B of all SNPs that have a derived allele frequency y in the archaic sample C and derived allele frequency z in the archaic sample D, but where the derived allele is at a frequency smaller than w in the sample A (Figure S1).

A common statistic that is indicative of population variation - expected heterozygosity (*π*) was previously found to be affected by archaic introgression in a serial founder model of human history [104]. We measured *π* as the average of 2*p*(1-p) over all sites in a window, where p is the sample derived allele frequency in the introgressed population.

### 4.2. Simulations

None of these statistics have been explicitly vetted under scenarios of AI so far, though the performance of *D* and *f_D_* has been previously evaluated for detecting local introgression [103]. Therefore, we aimed to test how each of the statistics described above performed in detecting AI in a 40 kb window. We chose this window size because the mean length of introgressed haplotypes in ref. [2] was 44,078 bp (Supplementary Information 13) and because 40kb is well above the length needed to reject incomplete lineage sorting for regions with moderate recombination rates [18]. We began by simulating a three population tree in Slim [105] with constant *N*_*e*_ = 10, 000, mutation rate equal to 1.5 * 10^−8^ per bp per generation, recombination rate equal to 10^−8^ per bp per generation, and split times emulating the African-Eurasian and Neanderthal-modern human split times (4,000 and 16,000 generations ago, respectively). We allowed for admixture between the most distantly diverged population and one of the closely related sister populations, at different rates: 2%, 10% and 25% (Figure 2.A). We use the lower (2%) rate to represent the Neanderthal genome-wide admixture into Eurasians, with Africans as the non-admixed population. The higher (10% and 25%) rates are meant to represent cases when a researcher is focusing on a particular region of the genome that has some *a priori* evidence for having been introgressed, thus pushing the local probability of introgression to high values, even though the genome-wide rate may be lower. Under each of the three admixture rate scenarios, we simulated regions that were evolving neutrally, regions where the central SNP was under weak positive additive selection (*s* = 0.01) and regions with a central SNP under strong selection (*s* = 0.1). We required the selected allele to be fixed in the archaic population prior to introgression, but allowed the allele to rise or decrease in frequency in the introgressed population, as determined by the strength of selection, its probability of entering the introgressed population and its starting frequency after introgression. Figure S2 shows the distributions of frequencies of the selected alleles in the introgressed population in the present.

We also tested how the statistics perform at detecting adaptive introgression when the alternative model is not a neutral intro-508 gression model, but a neutral model with ancestral structure (Figure 2.G). We followed a model described in Huerta-Sanchez et al (2014) and simulated a population in which an African population splits from archaic humans before Eurasians, but is allowed to exchange migrants with them. Afterwards, we split Eurasians and archaic humans. At that point, we stop the previous migration and only allow for migration between the Eurasian and African populations until the present, at double the previous rate. This is meant to generate loci where Eurasians and archaic humans share a more recent common ancestor with each other than with Africans, but because of ancient shared ancestry, not recent introgression. We simulated 3 scenarios, in which we set the per-generation ancient migration rate to be 0.01, 0.001 and 0.0001, respectively, and the recent migration rate to be 0.02, 0.002 and 0.0002, respectively. We call these the strong-, medium-, and weak-migration scenarios, respectively. The stronger the migration, the weaker the ancestral structure, as archaic-shared segments in Eurasians will tend to be removed by migration with Africans.

### 4.3. Plotting haplotype structure

The *Haplostrips* software (Marnetto et al. in prep.) was used to produce plots of haplotypes at candidate regions for AI. This software displays each SNP within a predefined region as a column, while each row represents a phased haplotype: the result is a heatmap. Each haplotype is labeled with a color that corresponds to the 1000 Genomes panel of its carrier individual. The haplotypes were first hierarchically clustered via the single agglomerative method based on Manhattan distances, using the *stats* library in R. The resulting dendrogram of haplotypes was then re-ordered by decreasing similarity to a reference sequence constructed so that it contains all the derived alleles found in the archaic genome (Altai Neanderthal or Denisova). The reordering is performed using the mininum distance method, so that haplotypes with more derived alleles shared with the archaic population are at the top of the plot. Derived alleles are represented as black spots and ancestral alleles are represented as white spots. Variant positions were filtered out when the site in the archaic genome had mapping quality less than 30 or genotype quality less than 40, or if the minor allele had a population frequency smaller than 5% in each of the present-day human populations included in the plot.

### 4.4. Hidden Markov Model

As haplotypes could look archaic simply because of ancestral structure or incomplete lineage sorting, we used a Hidden Markov Model (HMM) described in ref. [21] (which assumes an exponential distribution of admixture tract lengths [106, 107]), in order to verify that our candidate regions truly had archaic introgressed segments. This procedure also allowed us to confirm which of the archaic genomes was closest to the original source of introgression, as using a distant archaic source as input (for example, the Denisova genome when the true source is closest to the Neanderthal genome) produced shorter or less frequent inferred segments in the HMM output than when using the closer source genome.

The HMM we used requires us to specify a prior for the admixture rate. We tried two priors: 2% and 50%. The first was chosen because it is consistent with the genome-wide admixture rate for Neanderthals into Eurasians. The second, larger, value was chosen because each candidate region should *a priori* have a larger probability of being admixed, as they were found using statistics that are indicative of admixture in the first place. We observe almost no differences in the number of haplotypes inferred using either value. For example, for BNC2 - a well known candidate for AI [5, 6] - the frequency of sequences in EUR with inferred introgressed haplotypes under the 2% prior is 92.5%, while it is 93.1% under the 50% prior. However, the larger prior leads to longer and less fragmented introgressed chunks, as the HMM is less likely to transition into a non-introgressed state between two introgressed states. Therefore, all figures we show below were obtained using a 50% admixture prior. The admixture time was set to 1,900 generations ago and the recombination rate parameter was set to the local recombination rate in each region, following the recombination rate map in ref. [108]. A tract was called as introgressed if the posterior probability for introgression was higher than 90%. Under these parameters, the HMM has a specificity of 99.56%, a sensitivity of 36.07% and a false discovery rate of 1.15%.

## 5. Acknowledgments

We are grateful to Montgomery Slatkin, Rasmus Nielsen, Fergal P. Casey, Kirk Lohmueller and Amy Ko for helpful advice and comments. E.H.S. is supported by UC Merced start-up funds and National Science Foundation grant NSF-DEB-1557151. D.M. is supported by a University of Torino PhD Scholarship. F.R. is supported by National Institutes of Health grant R01HG003229 to Rasmus Nielsen.

---

**Table S1:** 95% quantiles of the *U_A,B,C_* statistic in a 40 kb window, under different demographic scenarios and archaic allele frequency cutoffs in the outgroup (A) and target (B) population panels. The demographic scenarios correspond to scenarios A, B, C and G from Figure 2. The bottlenecks were 5X and lasted 200 generations.

| Max. outgroup freq. | Min. target freq. | Scenario | 95% quantile under neutrality |
| --- | --- | --- | --- |
| 0.01 | 0.8 | Admixture (2%) | 0 |
| 0.01 | 0.8 | Admixture (10%) | 0 |
| 0.01 | 0.8 | Admixture (25%) | 0 |
| 0.01 | 0.8 | Ancestral Structure (strong mig.) | 0 |
| 0.01 | 0.8 | Ancestral Structure (medium mig.) | 1 |
| 0.01 | 0.8 | Ancestral Structure (weak mig.) | 18 |
| 0.01 | 0.8 | Admixture (2%), then bottleneck | 0 |
| 0.01 | 0.8 | Admixture (10%), then bottleneck | 0 |
| 0.01 | 0.8 | Admixture (25%), then bottleneck | 0.05 |
| 0.01 | 0.8 | Bottleneck, then admixture (2%) | 0 |
| 0.01 | 0.8 | Bottleneck, then admixture (10%) | 0 |
| 0.01 | 0.8 | Bottleneck, then admixture (25%) | 0 |
| 0.01 | 0.5 | Admixture (2%) | 2 |
| 0.01 | 0.5 | Admixture (10%) | 2 |
| 0.01 | 0.5 | Admixture (25%) | 5 |
| 0.01 | 0.5 | Ancestral Structure (strong mig.) | 0 |
| 0.01 | 0.5 | Ancestral Structure (medium mig.) | 5 |
| 0.01 | 0.5 | Ancestral Structure (weak mig.) | 22 |
| 0.01 | 0.5 | Admixture (2%), then bottleneck | 2 |
| 0.01 | 0.5 | Admixture (10%), then bottleneck | 2 |
| 0.01 | 0.5 | Admixture (25%), then bottleneck | 8 |
| 0.01 | 0.5 | Bottleneck, then admixture (2%) | 2 |
| 0.01 | 0.5 | Bottleneck, then admixture (10%) | 2 |
| 0.01 | 0.5 | Bottleneck, then admixture (25%) | 6 |
| 0.01 | 0.2 | Admixture (2%) | 6 |
| 0.01 | 0.2 | Admixture (10%) | 13 |
| 0.01 | 0.2 | Admixture (25%) | 29.05 |
| 0.01 | 0.2 | Ancestral Structure (strong mig.) | 0 |
| 0.01 | 0.2 | Ancestral Structure (medium mig.) | 9.05 |
| 0.01 | 0.2 | Ancestral Structure (weak mig.) | 25 |
| 0.01 | 0.2 | Admixture (2%), then bottleneck | 6 |
| 0.01 | 0.2 | Admixture (10%), then bottleneck | 17 |
| 0.01 | 0.2 | Admixture (25%), then bottleneck | 30 |
| 0.01 | 0.2 | Bottleneck, then admixture (2%) | 8 |
| 0.01 | 0.2 | Bottleneck, then admixture (10%) | 13.05 |
| 0.01 | 0.2 | Bottleneck, then admixture (25%) | 29 |
| 0.01 | 0 | Admixture (2%) | 24 |
| 0.01 | 0 | Admixture (10%) | 37 |
| 0.01 | 0 | Admixture (25%) | 39 |
| 0.01 | 0 | Ancestral Structure (strong mig.) | 3 |
| 0.01 | 0 | Ancestral Structure (medium mig.) | 12.05 |
| 0.01 | 0 | Ancestral Structure (weak mig.) | 27 |
| 0.01 | 0 | Admixture (2%), then bottleneck | 21 |
| 0.01 | 0 | Admixture (10%), then bottleneck | 34 |
| 0.01 | 0 | Admixture (25%), then bottleneck | 38 |
| 0.01 | 0 | Bottleneck, then admixture (2%) | 28 |
| 0.01 | 0 | Bottleneck, then admixture (10%) | 34.05 |
| 0.01 | 0 | Bottleneck, then admixture (25%) | 37.05 |
| 0.1 | 0.8 | Admixture (2%) | 0 |
| 0.1 | 0.8 | Admixture (10%) | 2 |
| 0.1 | 0.8 | Admixture (25%) | 2 |
| 0.1 | 0.8 | Ancestral Structure (strong mig.) | 0 |
| 0.1 | 0.8 | Ancestral Structure (medium mig.) | 11 |
| 0.1 | 0.8 | Ancestral Structure (weak mig.) | 23.05 |
| 0.1 | 0.8 | Admixture (2%), then bottleneck | 0 |
| 0.1 | 0.8 | Admixture (10%), then bottleneck | 2 |
| 0.1 | 0.8 | Admixture (25%), then bottleneck | 2 |
| 0.1 | 0.8 | Bottleneck, then admixture (2%) | 1 |
| 0.1 | 0.8 | Bottleneck, then admixture (10%) | 2 |
| 0.1 | 0.8 | Bottleneck, then admixture (25%) | 2 |
| 0.1 | 0.5 | Admixture (2%) | 5 |
| 0.1 | 0.5 | Admixture (10%) | 6 |
| 0.1 | 0.5 | Admixture (25%) | 12 |
| 0.1 | 0.5 | Ancestral Structure (strong mig.) | 0 |
| 0.1 | 0.5 | Ancestral Structure (medium mig.) | 17 |
| 0.1 | 0.5 | Ancestral Structure (weak mig.) | 29 |
| 0.1 | 0.5 | Admixture (2%), then bottleneck | 6 |
| 0.1 | 0.5 | Admixture (10%), then bottleneck | 7 |
| 0.1 | 0.5 | Admixture (25%), then bottleneck | 12 |
| 0.1 | 0.5 | Bottleneck, then admixture (2%) | 6 |
| 0.1 | 0.5 | Bottleneck, then admixture (10%) | 6.05 |
| 0.1 | 0.5 | Bottleneck, then admixture (25%) | 12 |
| 0.1 | 0.2 | Admixture (2%) | 12 |
| 0.1 | 0.2 | Admixture (10%) | 18.05 |
| 0.1 | 0.2 | Admixture (25%) | 35 |
| 0.1 | 0.2 | Ancestral Structure (strong mig.) | 4 |
| 0.1 | 0.2 | Ancestral Structure (medium mig.) | 21 |
| 0.1 | 0.2 | Ancestral Structure (weak mig.) | 32.05 |
| 0.1 | 0.2 | Admixture (2%), then bottleneck | 14 |
| 0.1 | 0.2 | Admixture (10%), then bottleneck | 22 |
| 0.1 | 0.2 | Admixture (25%), then bottleneck | 37 |
| 0.1 | 0.2 | Bottleneck, then admixture (2%) | 14 |
| 0.1 | 0.2 | Bottleneck, then admixture (10%) | 20 |
| 0.1 | 0.2 | Bottleneck, then admixture (25%) | 37 |
| 0.1 | 0 | Admixture (2%) | 29 |
| 0.1 | 0 | Admixture (10%) | 44 |
| 0.1 | 0 | Admixture (25%) | 45 |
| 0.1 | 0 | Ancestral Structure (strong mig.) | 11 |
| 0.1 | 0 | Ancestral Structure (medium mig.) | 25 |
| 0.1 | 0 | Ancestral Structure (weak mig.) | 34 |
| 0.1 | 0 | Admixture (2%), then bottleneck | 28 |
| 0.1 | 0 | Admixture (10%), then bottleneck | 40 |
| 0.1 | 0 | Admixture (25%), then bottleneck | 44 |
| 0.1 | 0 | Bottleneck, then admixture (2%) | 35 |
| 0.1 | 0 | Bottleneck, then admixture (10%) | 41 |
| 0.1 | 0 | Bottleneck, then admixture (25%) | 45 |

**Table S2:**
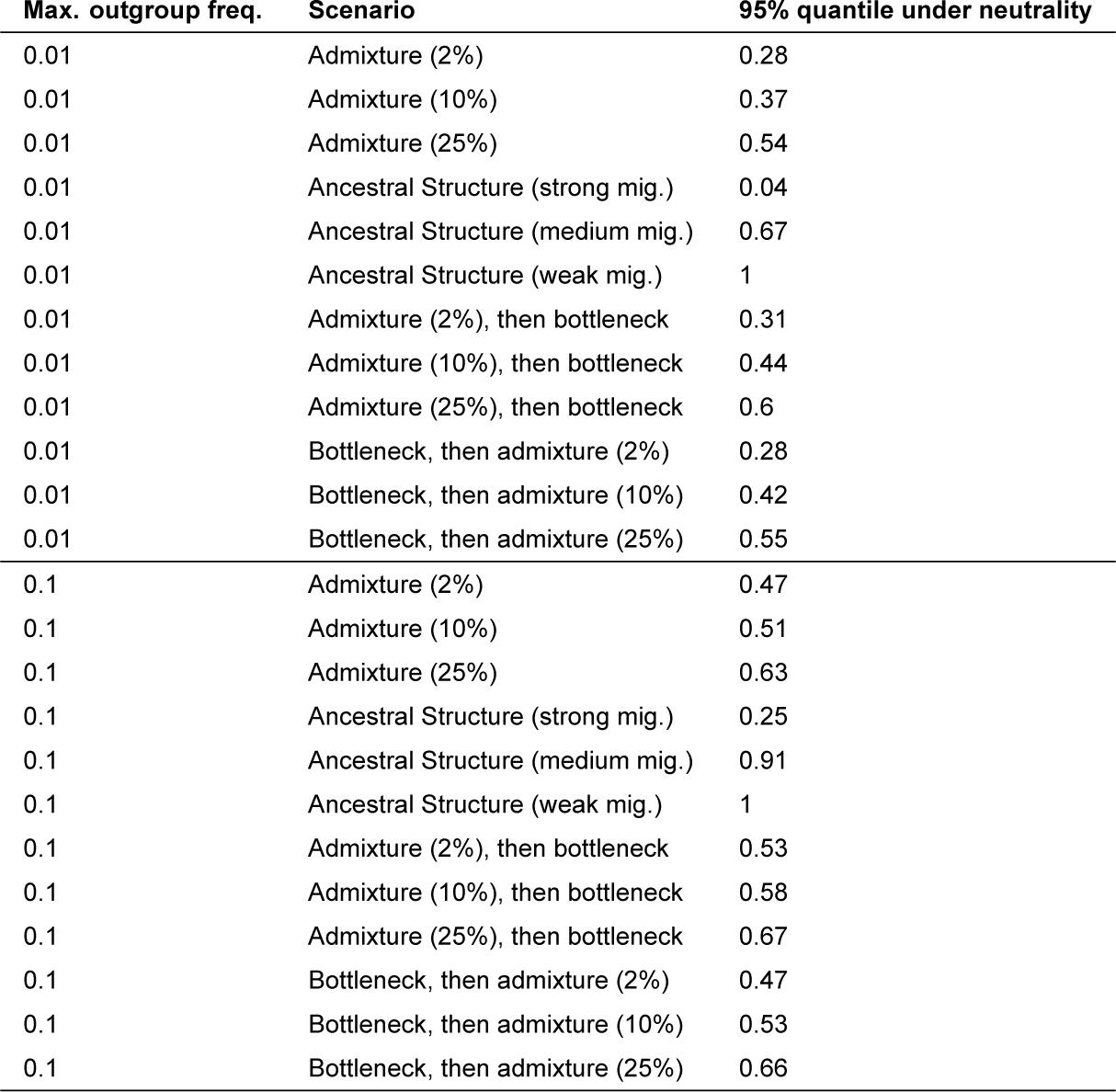
95% quantiles of the *Q*95_*A,B,C*_ statistic in a 40 kb window, under different demographic scenarios and archaic allele frequency cutoffs in the outgroup (A) population panel. The demographic scenarios correspond to scenarios A, B, C and G from Figure 2.

**Table S3:** 40 kb windows that lie in the highest 99.9% quantile of both *U_A,B,Nea,Den_* and *Q*95_*A,B,Nea,Den*_ for various outgroup panels A and target panels B, using an outgroup maximum frequency cutoff of 1%, and using different target allele frequency cutoffs (20%, 50%). For each region, we also show other statistics indicative of AI for reference. We partitioned the 1000 Genomes panels into outgroup panel A and target panel B in different ways (column “Mode”), depending on the signals we were looking for. These modes of partitioning are as follows. “Populations” = outgroup panel was the combination of all the populations that were not the target panel. “PopulationsB” = outgroup panel was the combination of all African panels (excluding admixed African-Americans), while target panel was one of the non-African panels. “Continents” = target panel was either the EUR continental panel (in which case the outgroup was AFR + EAS) or the EAS continental panel (in which case the outgroup was AFR + EUR). “ContinentsB” = target panel was the EUR continental panel (in which case the outgroup was AFR + EAS + SAS) or the EAS continental panel (in which case the outgroup was AFR + EUR + SAS) or the SAS continental panel (in which case the outgroup was AFR + EUR + EAS). “Eurasia” = target panel was EUR + EAS, while outgroup panel was AFR. https://www.dropbox.com/s/p9k94i2c50rincq/Extreme_gene_table.xlsx?dl=0

**Figure S1:**
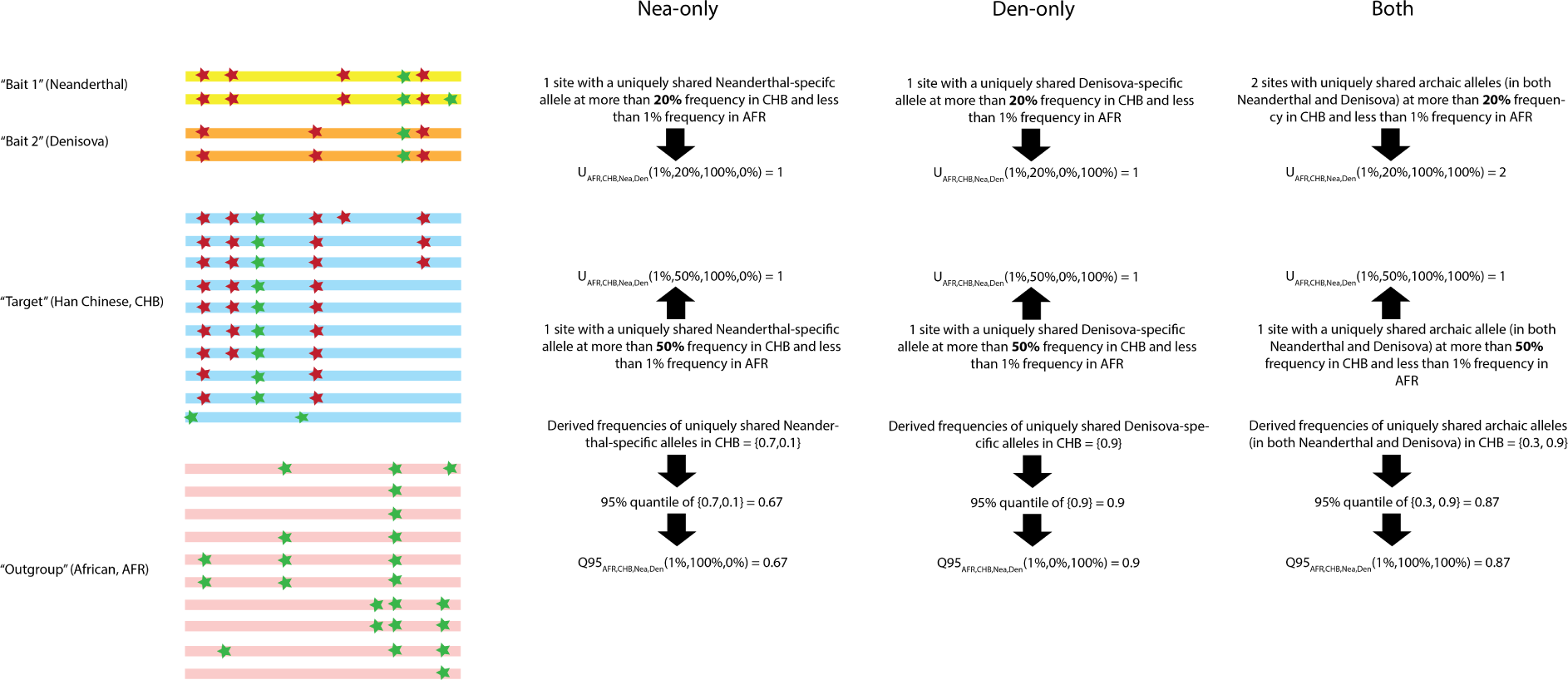
Schematic illustration of the way the *U_A,B,C,D_* and *Q*95_*A,B,C,D*_ statistics are calculated.

**Figure S2:**
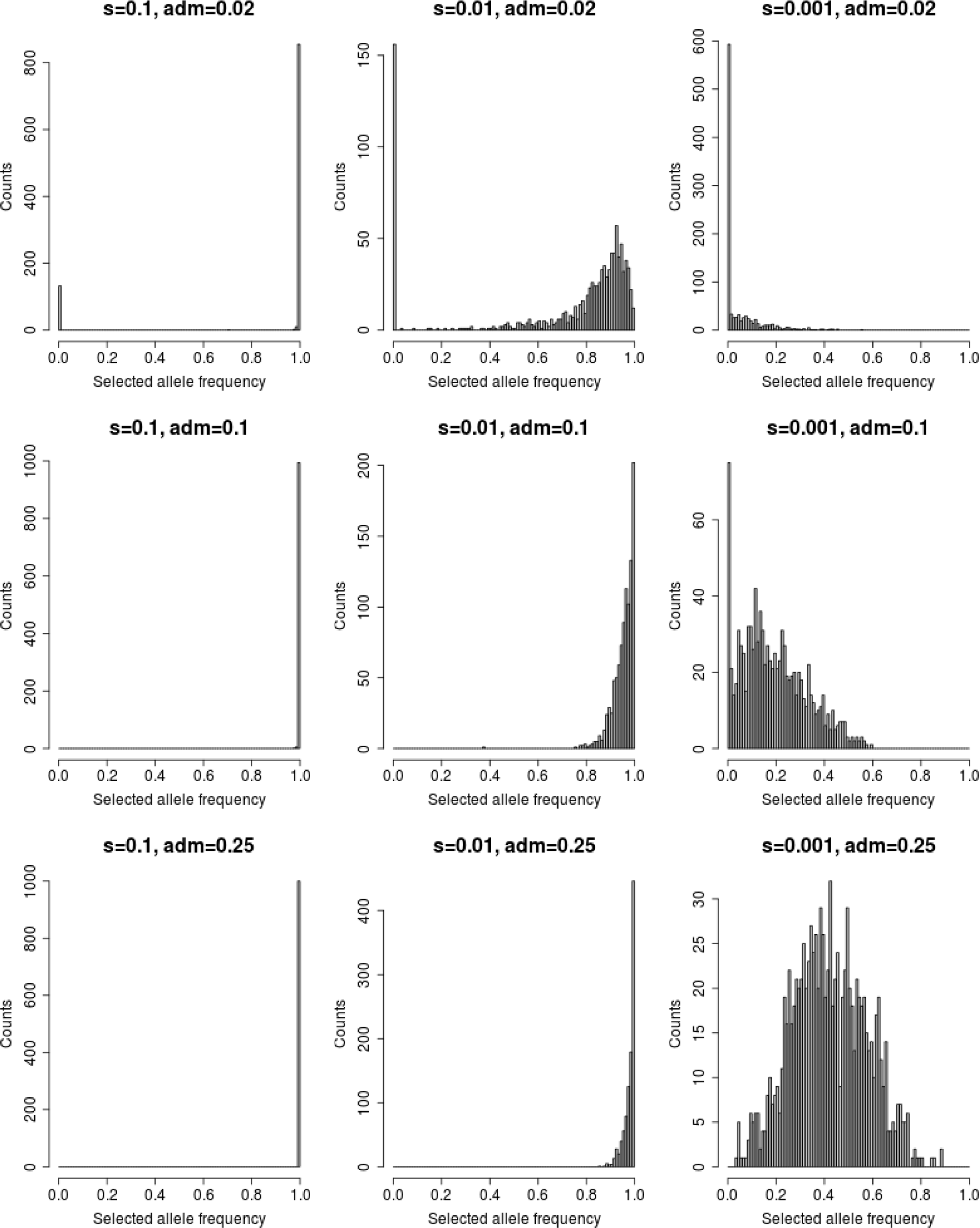
Histograms of the frequencies of the selected allele in the introgressed population in the present, for each AI scenario under constant population sizes.

**Figure S3:**
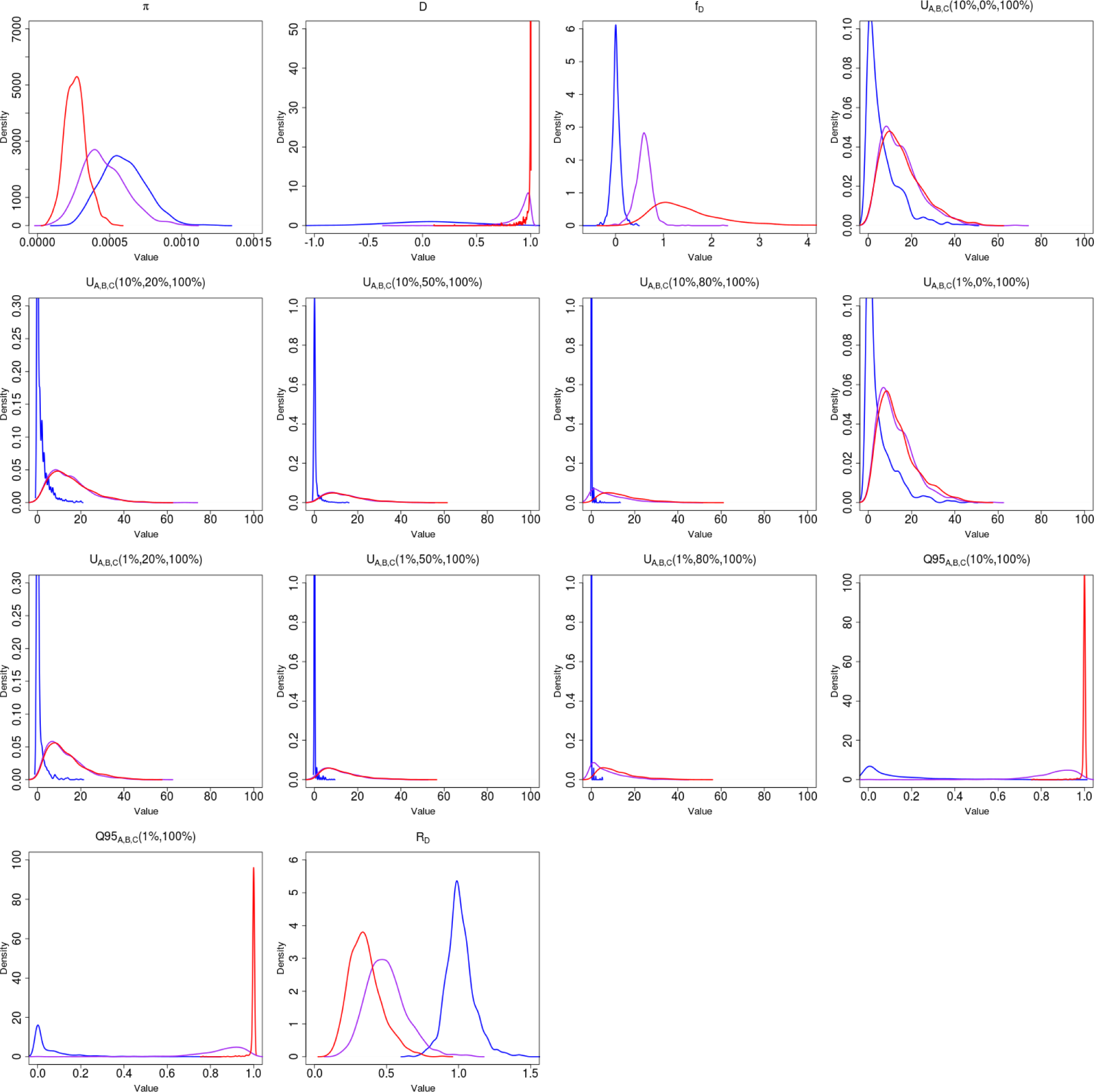
Density of various statistics meant to detect genetic patterns left by adaptive introgression, for three scenarios: neutrality (s=0) in blue, weak adaptive introgression (s=0.01) in purple and strong adaptive introgression (s=0.1) in red. The demography was the same as in Figure 3 and the admixture rate was set at 2%. See Table 1 for a definition of the statistics shown.

**Figure S4:**
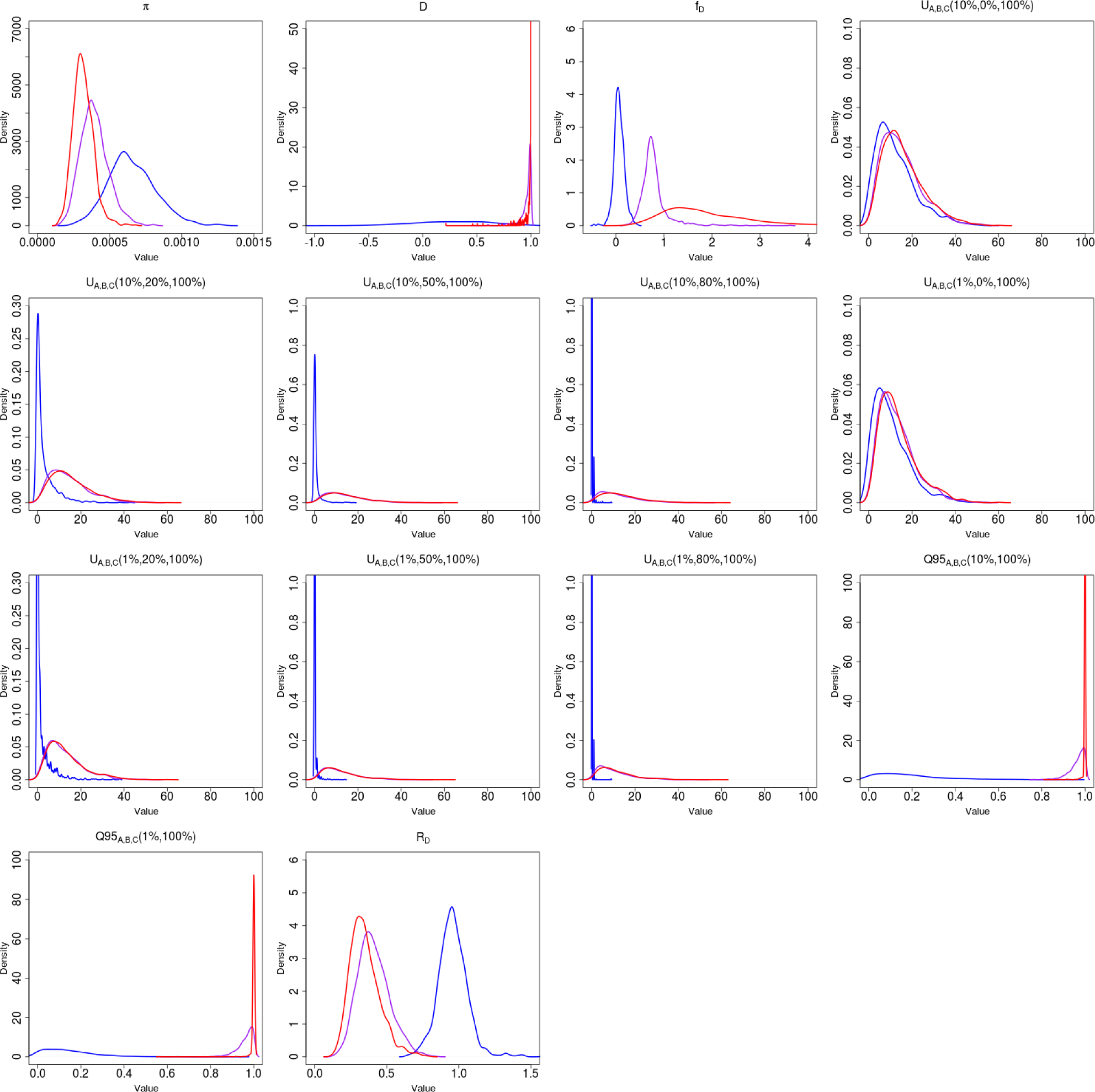
Density of various statistics meant to detect genetic patterns left by adaptive introgression, for three scenarios: neutrality (s=0) in blue, weak adaptive introgression (s=0.01) in purple and strong adaptive introgression (s=0.1) in red. The demography was the same as in Figure 3 and the admixture rate was set at 10%. See Table 1 for a definition of the statistics shown.

**Figure S5:**
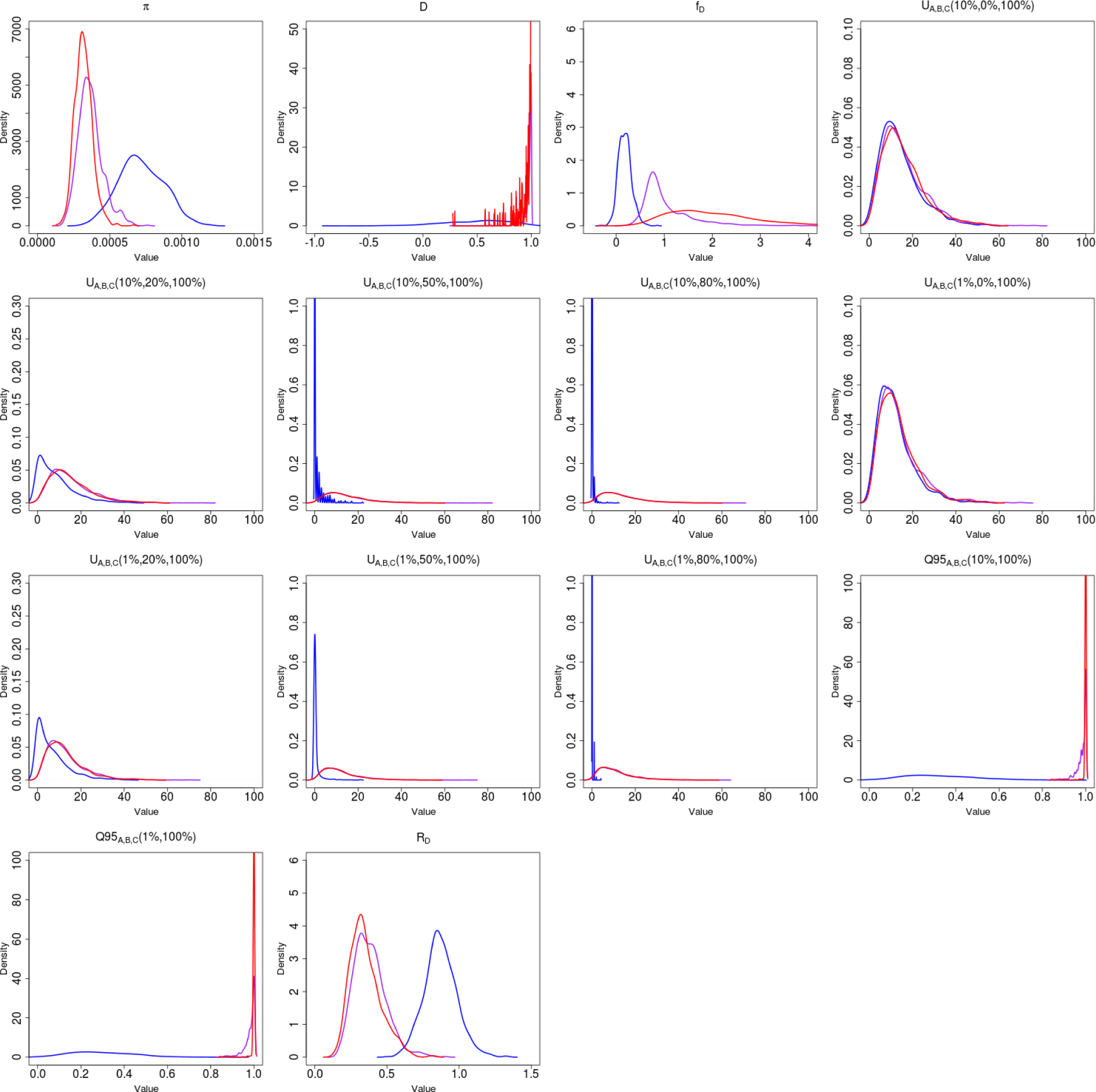
Density of various statistics meant to detect genetic patterns left by adaptive introgression, for three scenarios: neutrality (s=0) in blue, weak adaptive introgression (s=0.01) in purple and strong adaptive introgression (s=0.1) in red. The demography was the same as in Figure 3 and the admixture rate was set at 25%. See Table 1 for a definition of the statistics shown.

**Figure S6:**
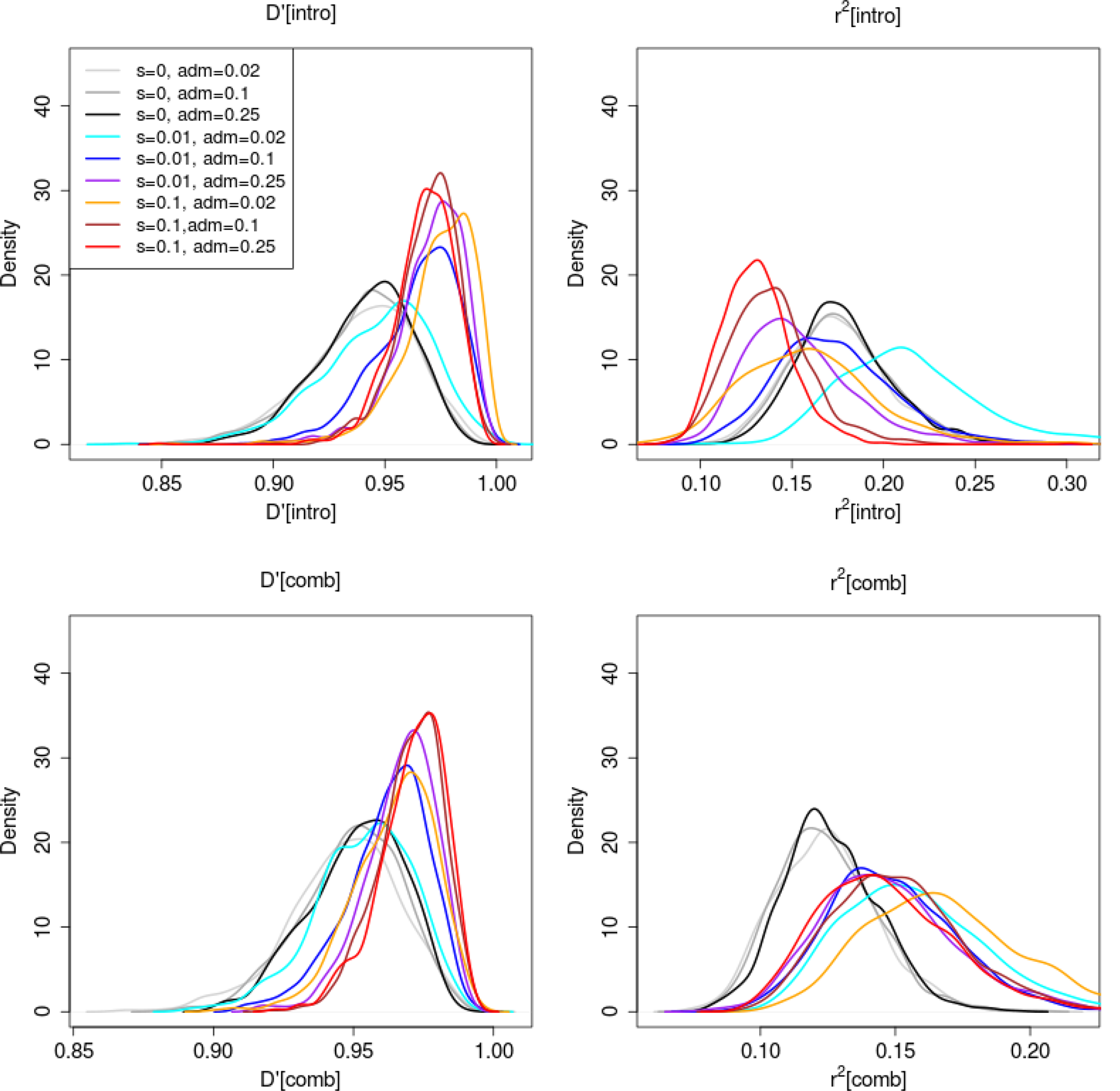
Density of statistics that detect patterns of linkage disequilibrium for various neutral and adaptive introgression scenarios. See Table 1 for a definition of the statistics shown.

**Figure S7:**
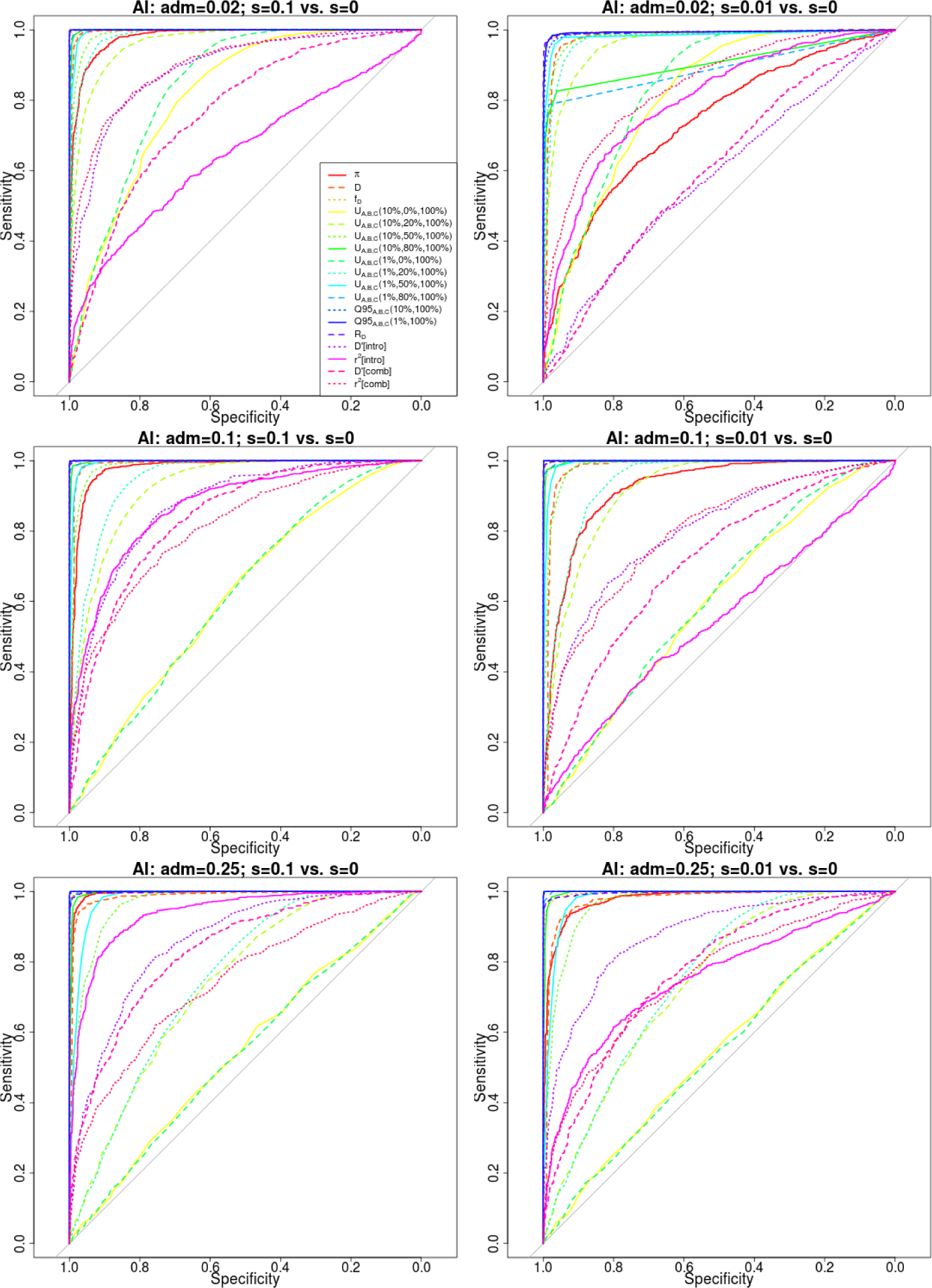
Receiver operating characteristic curves for adaptive introgression with constant population size, using 1,000 simulations of adaptive introgression, under various selection (s=0.1, s=0.01) and admixture rate (2%, 10%, 25%) regimes. Populations A and B split from each other 4,000 generations ago, and their ancestral population split from population C 16,000 generations ago. Population sizes were set at 2*N* = 20, 000. The admixture event occurred 1,600 generations ago from population C into population B,

**Figure S8:**
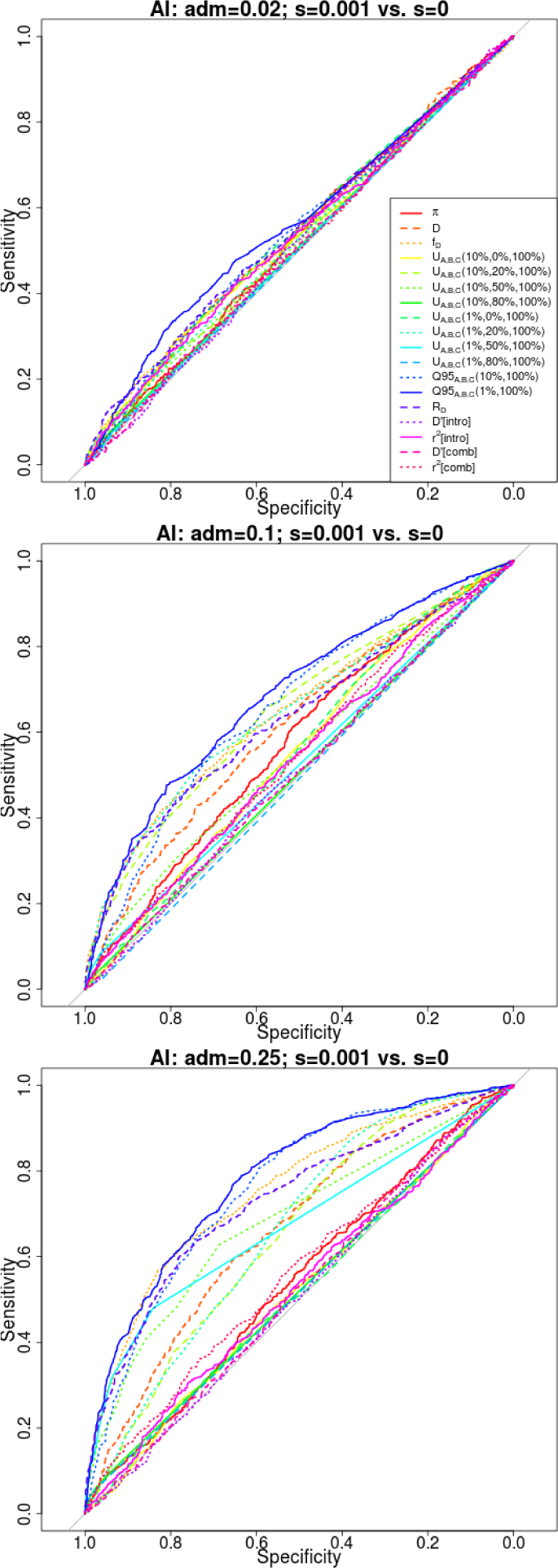
Receiver operating characteristic curves for adaptive introgression with constant population size, using 1,000 simulations of adaptive introgression, under weak selection (s=0.001) and different admixture rate (2%, 10%, 25%) regimes. Populations A and B split from each other 4,000 generations ago, and their ancestral population split from population C 16,000 generations ago. Population sizes were set at 2*N* = 20, 000. The admixture event occurred 1,600 generations ago from population C into population B,

**Figure S9:**
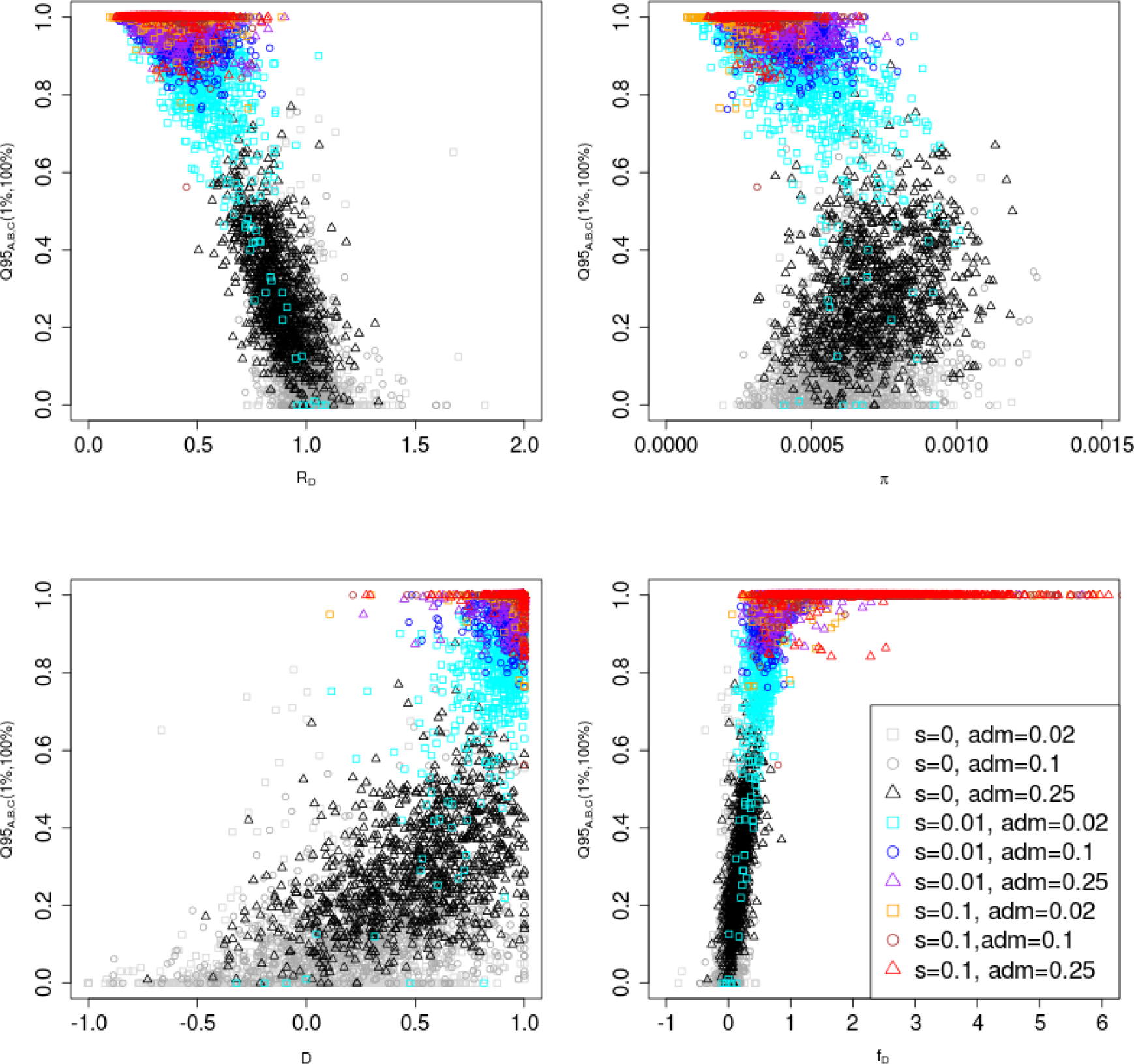
Joint distribution of *Q*95_*A,B,C*_ (1%,100%) and other statistics (*R_D_*, *π*, *D* and *f_D_*). 100 individuals were sampled from panel A, 100 from panel B and 2 from panel C. The demographic parameters were the same as in Figure 3.

**Figure S10:**
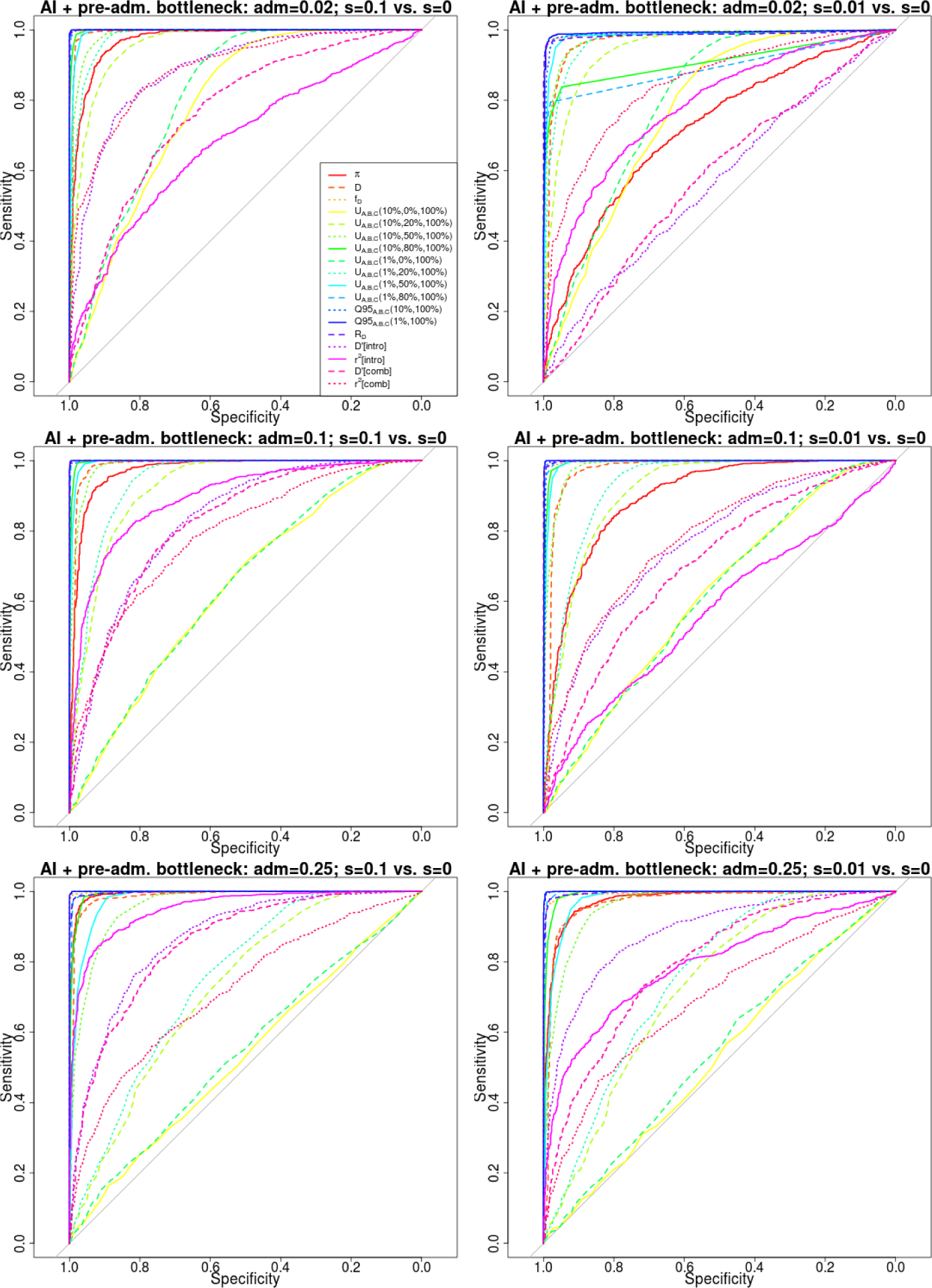
Receiver operating characteristic curves for adaptive introgression with a pre-admixture bottleneck, using 1,000 simulations under adaptive introgression. We simulated the same demography as in Figure 3, but also included a 5X bottleneck in population *B* before the introgression event, starting 3,000 generations ago and finishing 2,800 generations ago.

**Figure S11:**
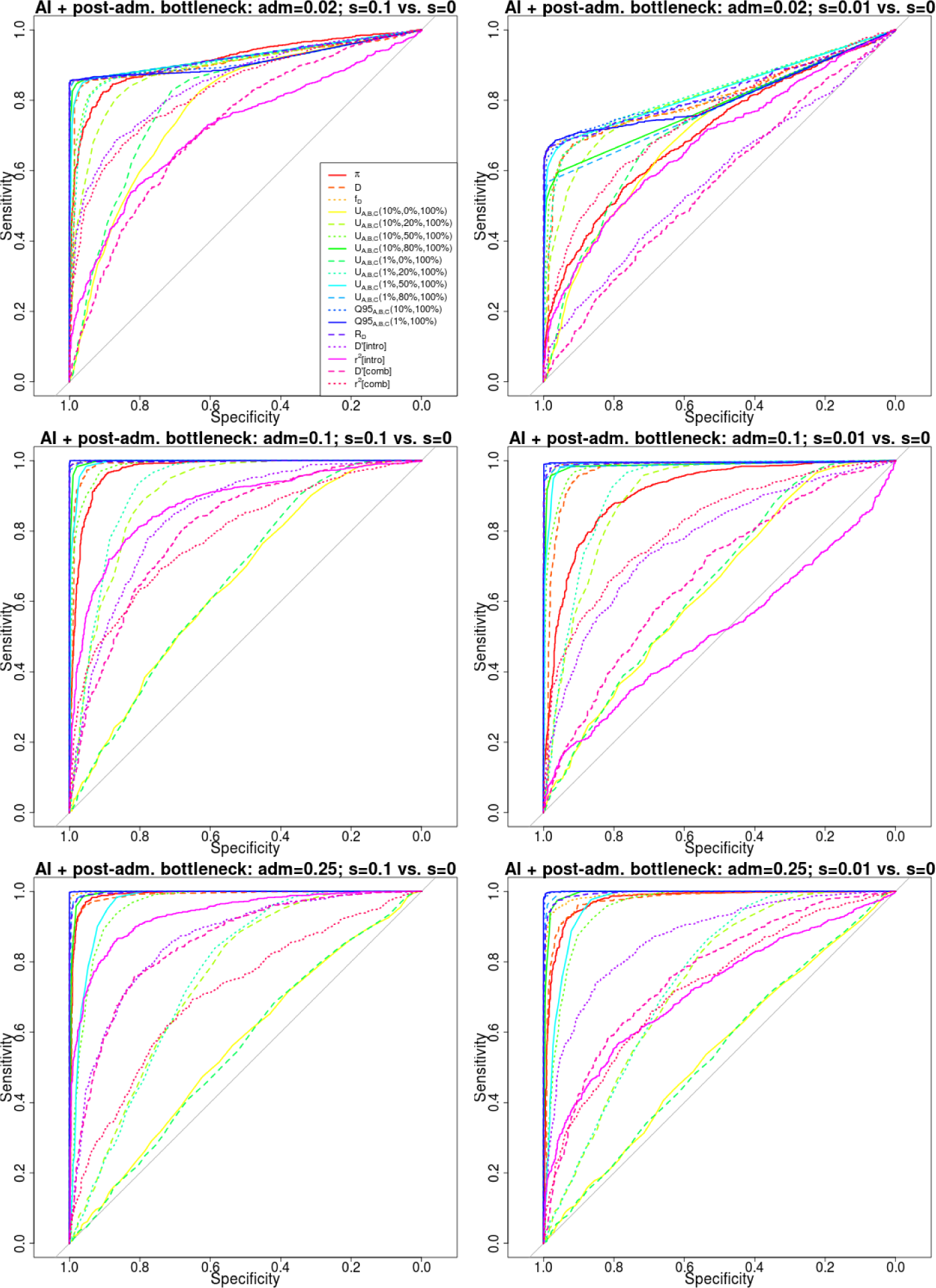
Receiver operating characteristic curves for adaptive introgression with a post-admixture bottleneck, using 1,000 simulations under adaptive introgression. We simulated the same demography as in Figure 3, but also included a 5X bottleneck in population *B* after the introgression event, starting 1,400 generations ago and finishing 1,200 generations ago.

**Figure S12:**
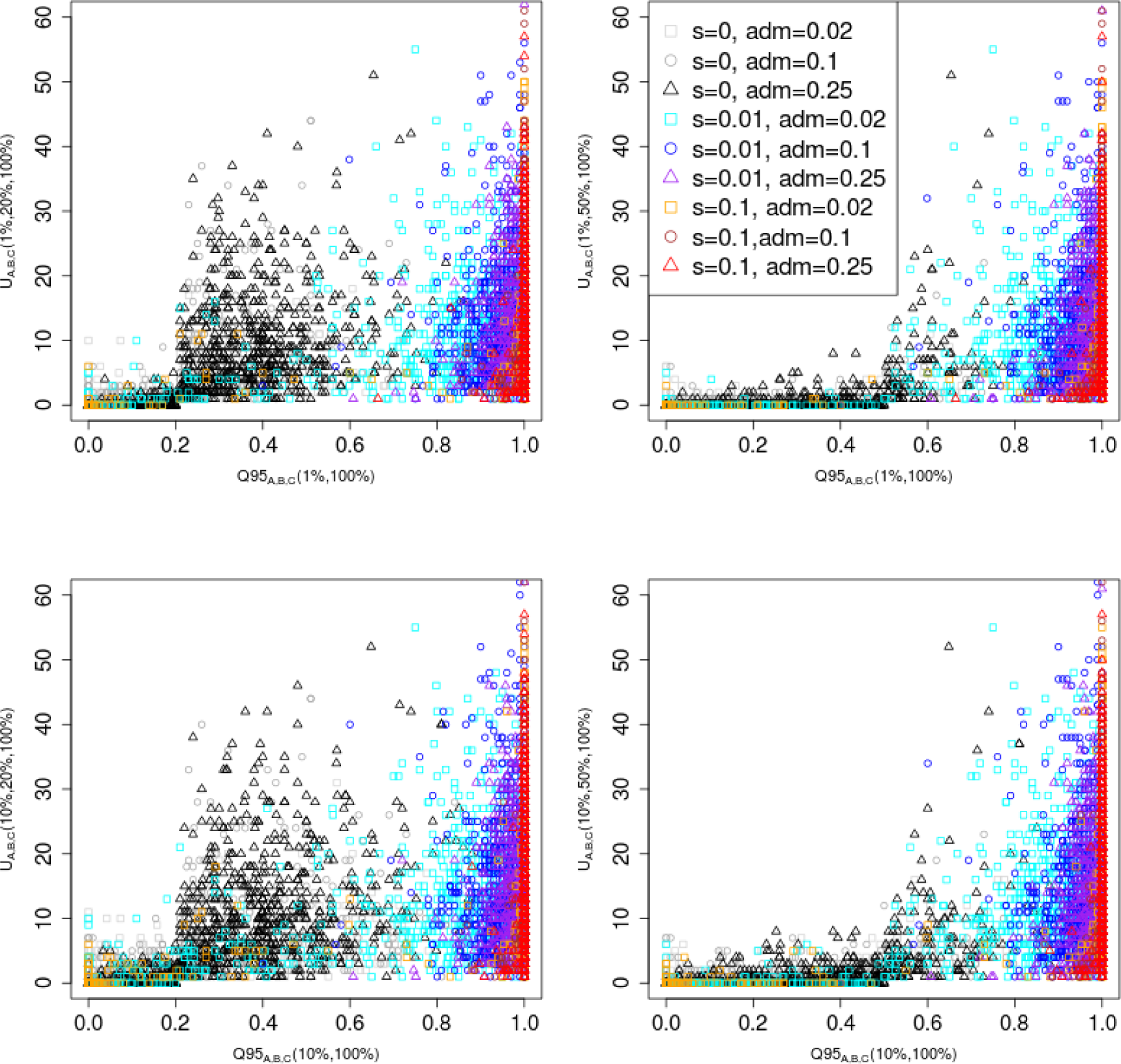
Joint distribution of *Q*95_*A,B,C*_ (*w, y*) and *U_A,B,C_*(*w, x, y*) for different choices of w (1%, 10%) and x (20%, 50%). We set y to 100% in all cases. 100 individuals were sampled from panel A, 100 from panel B and 2 from panel C. In this case, we included a 5X bottleneck in population *B* after the introgression event, starting 1,400 generations ago and finishing 1,200 generations ago.

**Figure S13:**
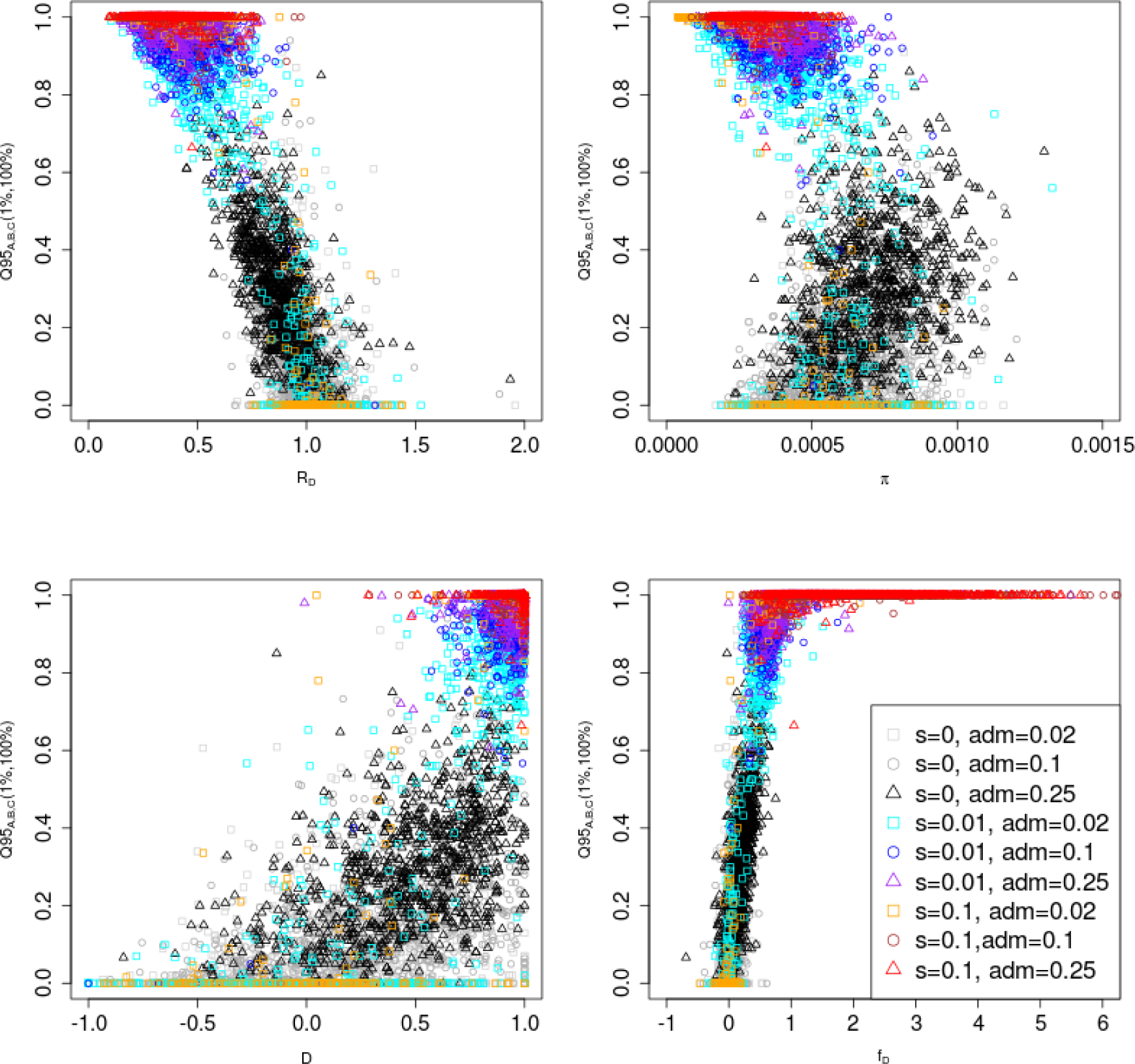
Joint distribution of *Q*95_*A,B,C*_ (1%,100%) and other statistics (*R_D_*, *π*, *D* and *f_D_*). 100 individuals were sampled from panel A, 100 from panel B and 2 from panel C. In this case, we included a 5X bottleneck in population *B* after the introgression event, starting 1,400 generations ago and finishing 1,200 generations ago.

**Figure S14:**
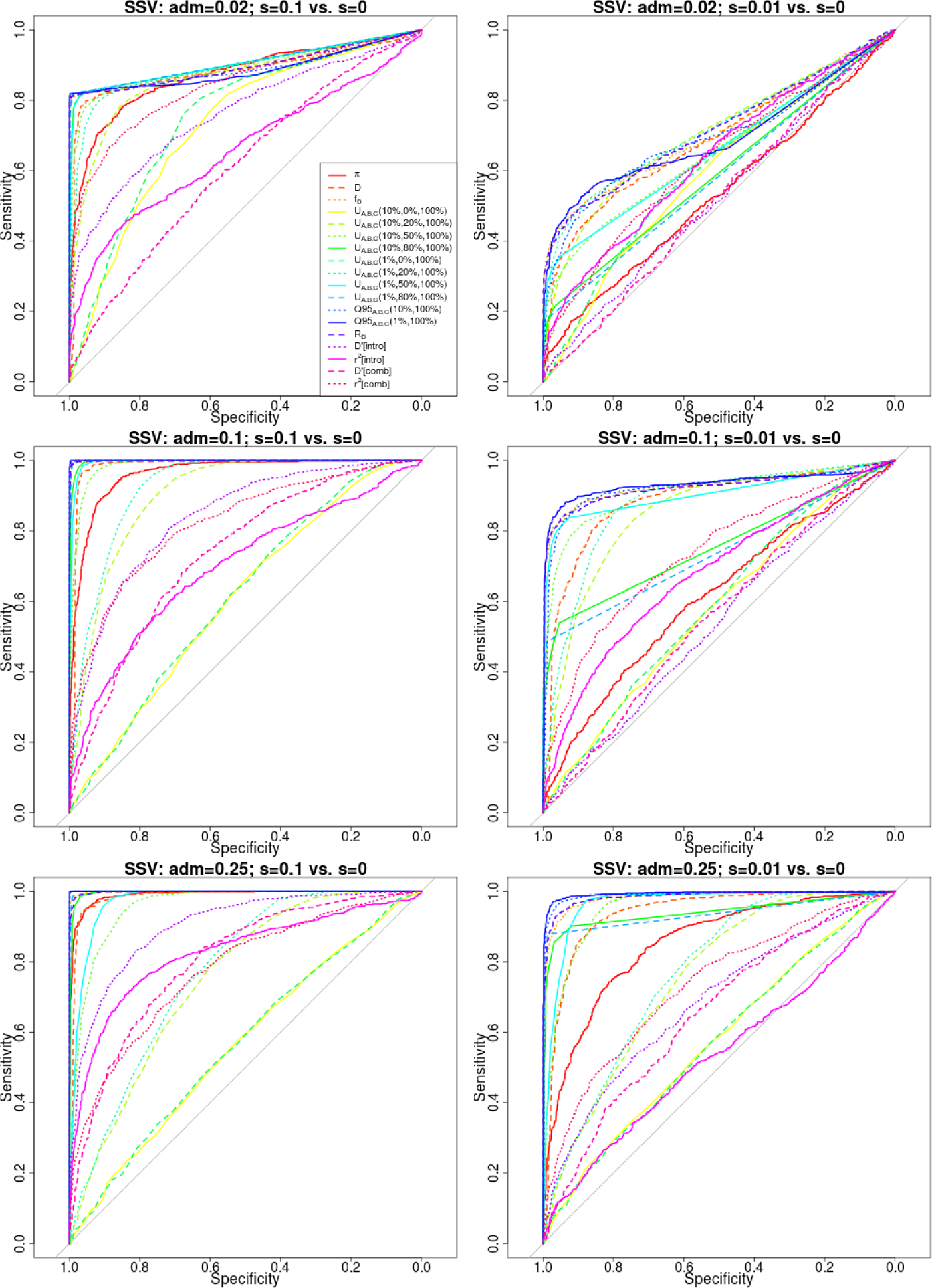
Receiver operating characteristic curves for adaptive introgression with an intermediate neutrality period. We simulated the same demography as in Figure 3, but changed the selection coefficient of the beneficial variant to be 0 right after the introgression event (1,600 generations ago). If still present in population *B*, the variant regained its original coefficient 800 generations ago.

**Figure S15:**
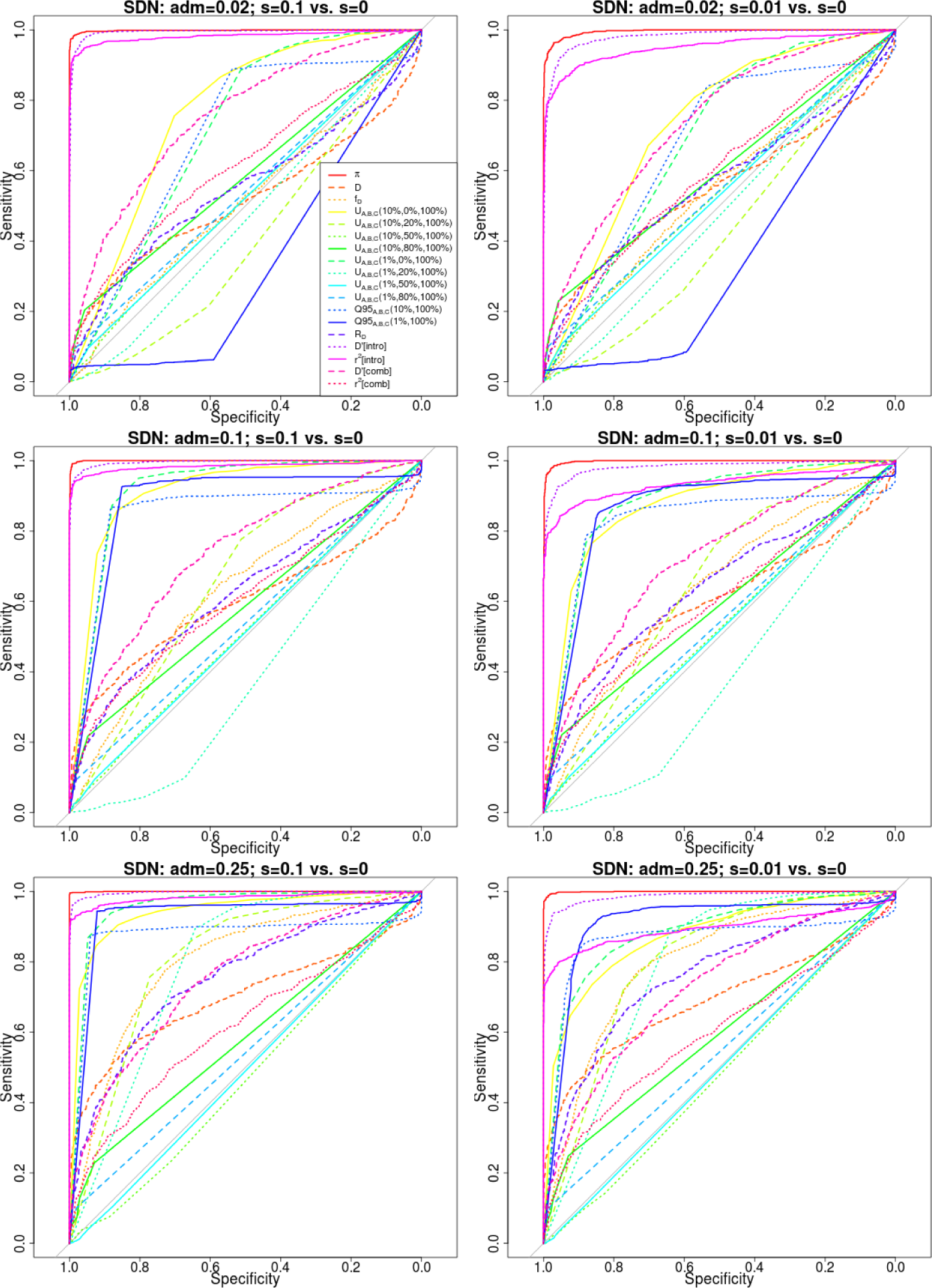
Receiver operating characteristic curves for a selective sweep from de novo mutation. We simulated the same demography as in Figure 3, but rather than introducing the beneficial variant in the introgressed population via admixture from an archaic population, we introduced it by mutation in the introgressed population (*B*) 3,900 generations ago.

**Figure S16:**
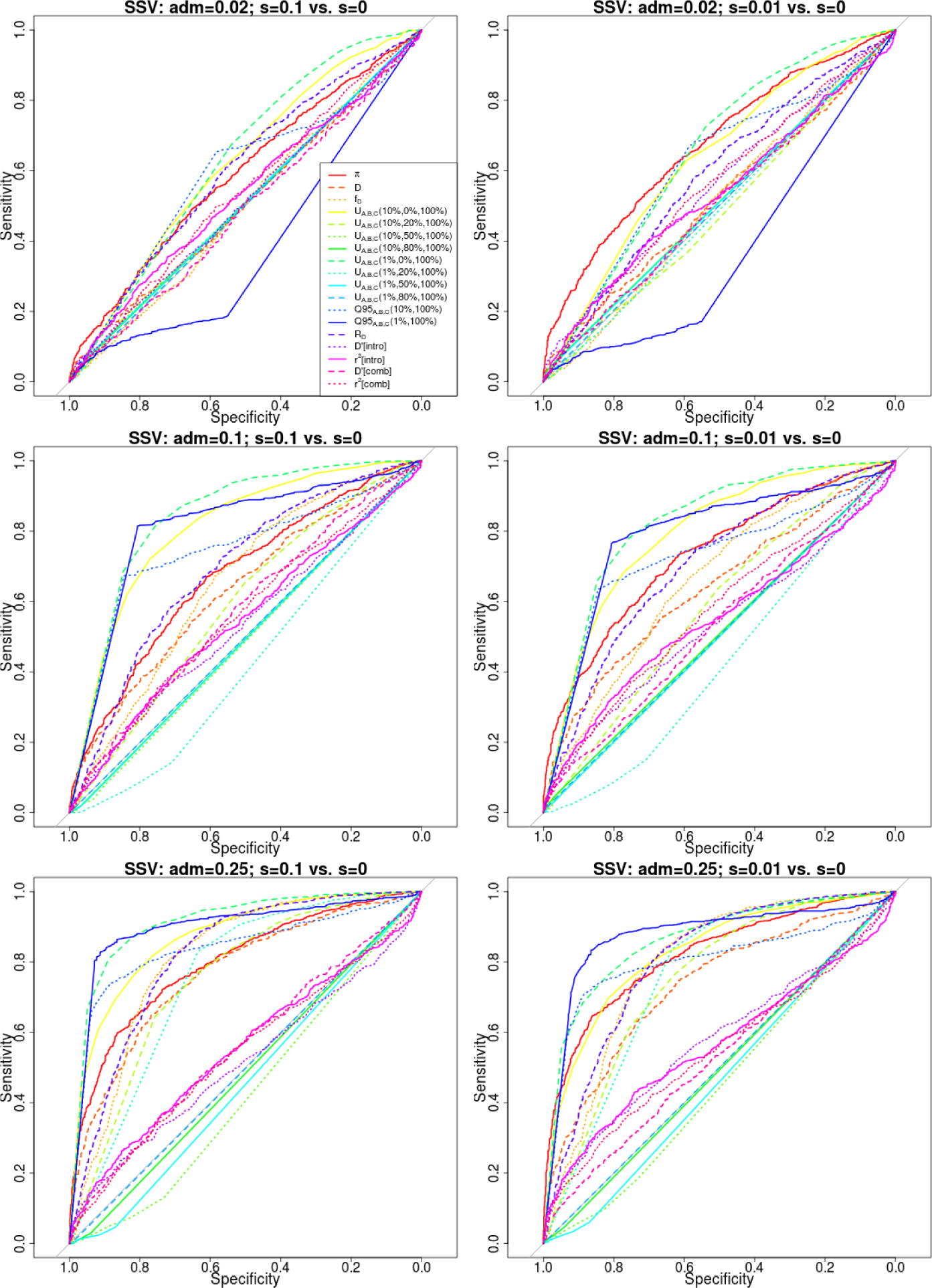
Receiver operating characteristic curves for selection from standing variation. We simulated the same demography as in Figure 3, but rather than introducing the beneficial variant in the introgressed population via admixture from an archaic population, we introduced it with a starting frequency of 20% in the introgressed population (*B*) 3,900 generations ago.

**Figure S17:**
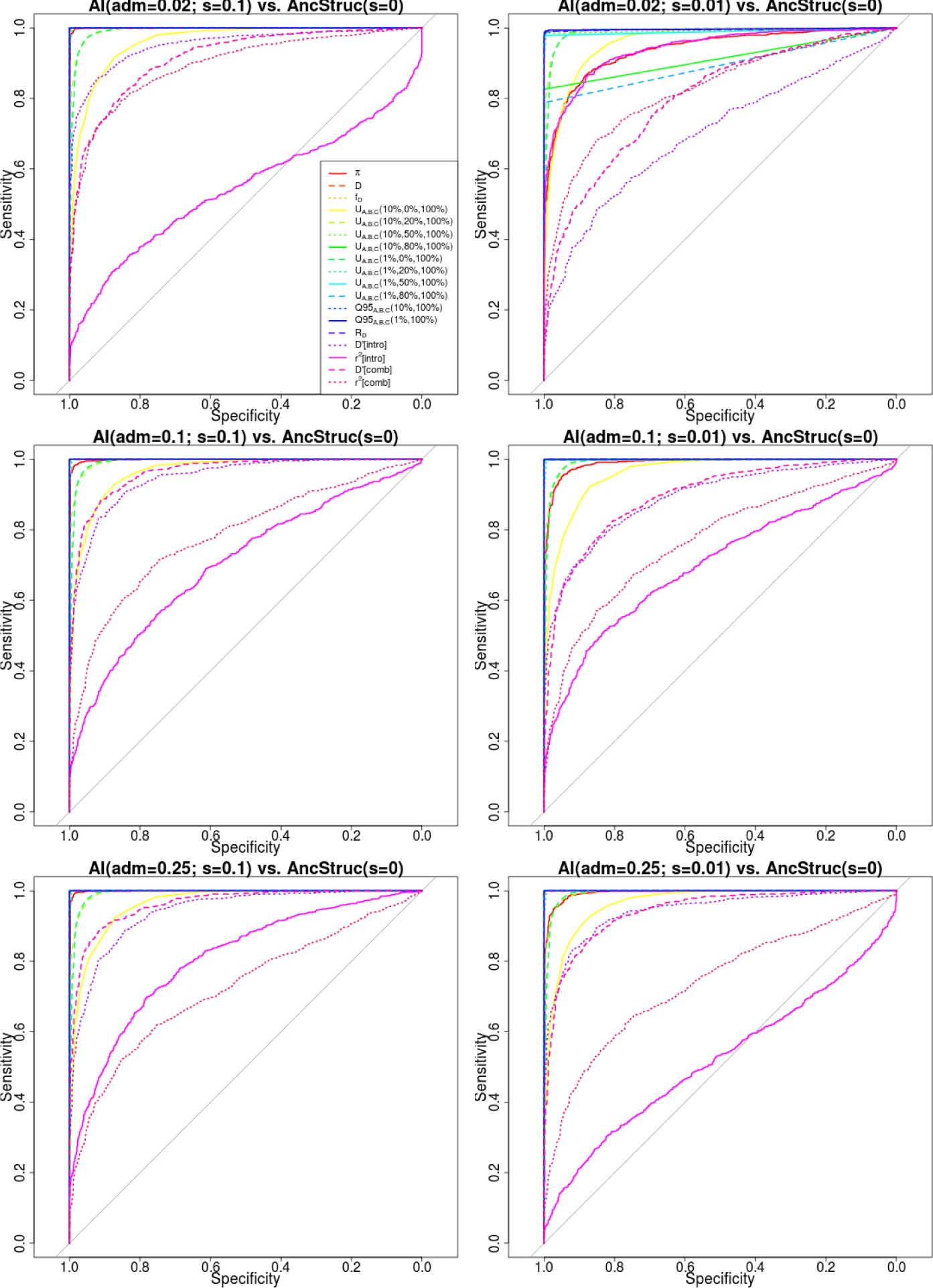
Receiving operating characteristic curves for adaptive introgression against a neutral ancestral structure model with strong migration rates. The demographic scenario for adaptive introgression was the same as in Figure 3. For a description of the ancestral structure model, see main text.

**Figure S18:**
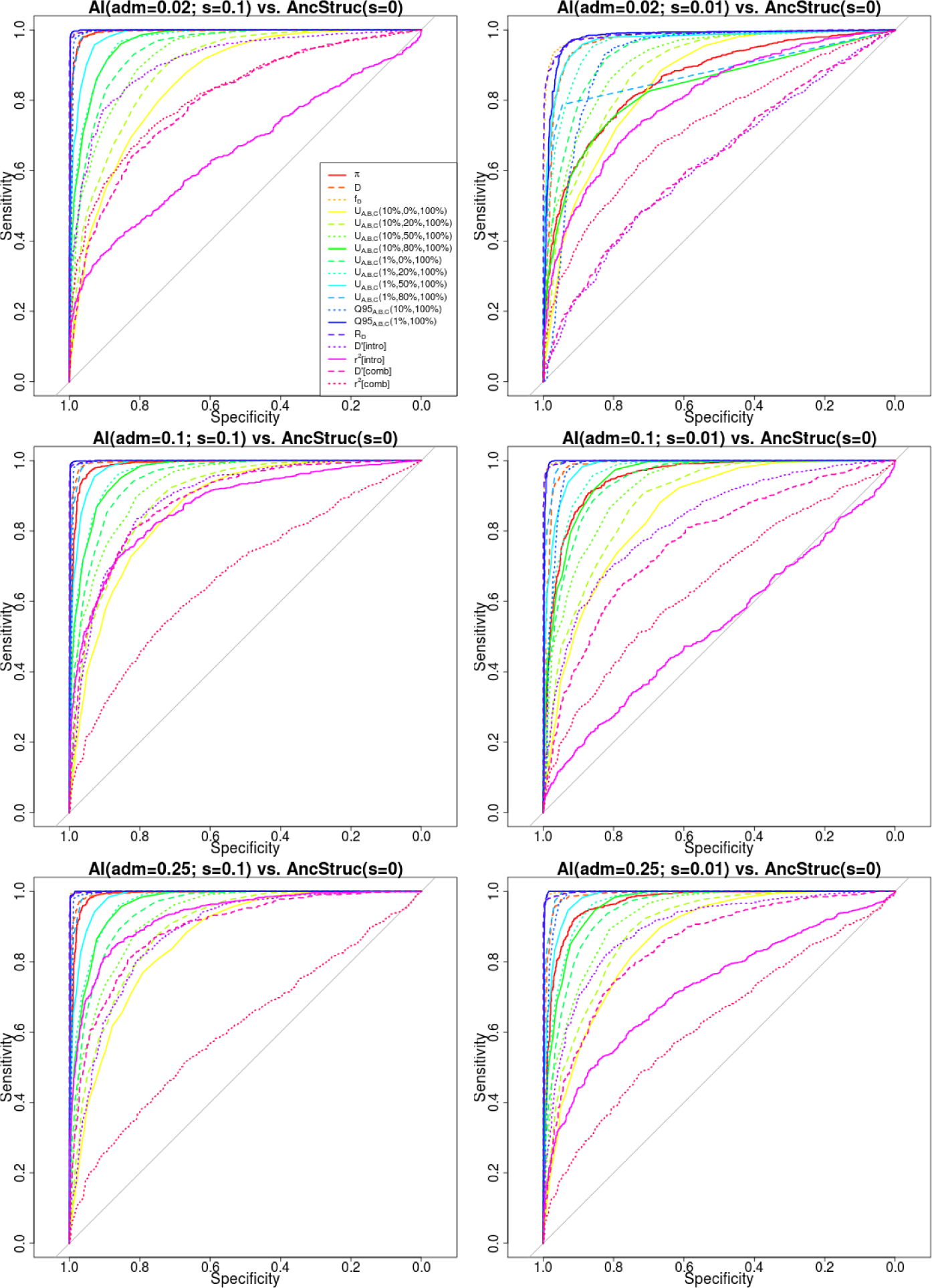
Receiving operating characteristic curves for adaptive introgression against a neutral ancestral structure model with intermediate migration rates. The demographic scenario for adaptive introgression was the same as in Figure 3. For a description of the ancestral structure model, see main text.

**Figure S19:**
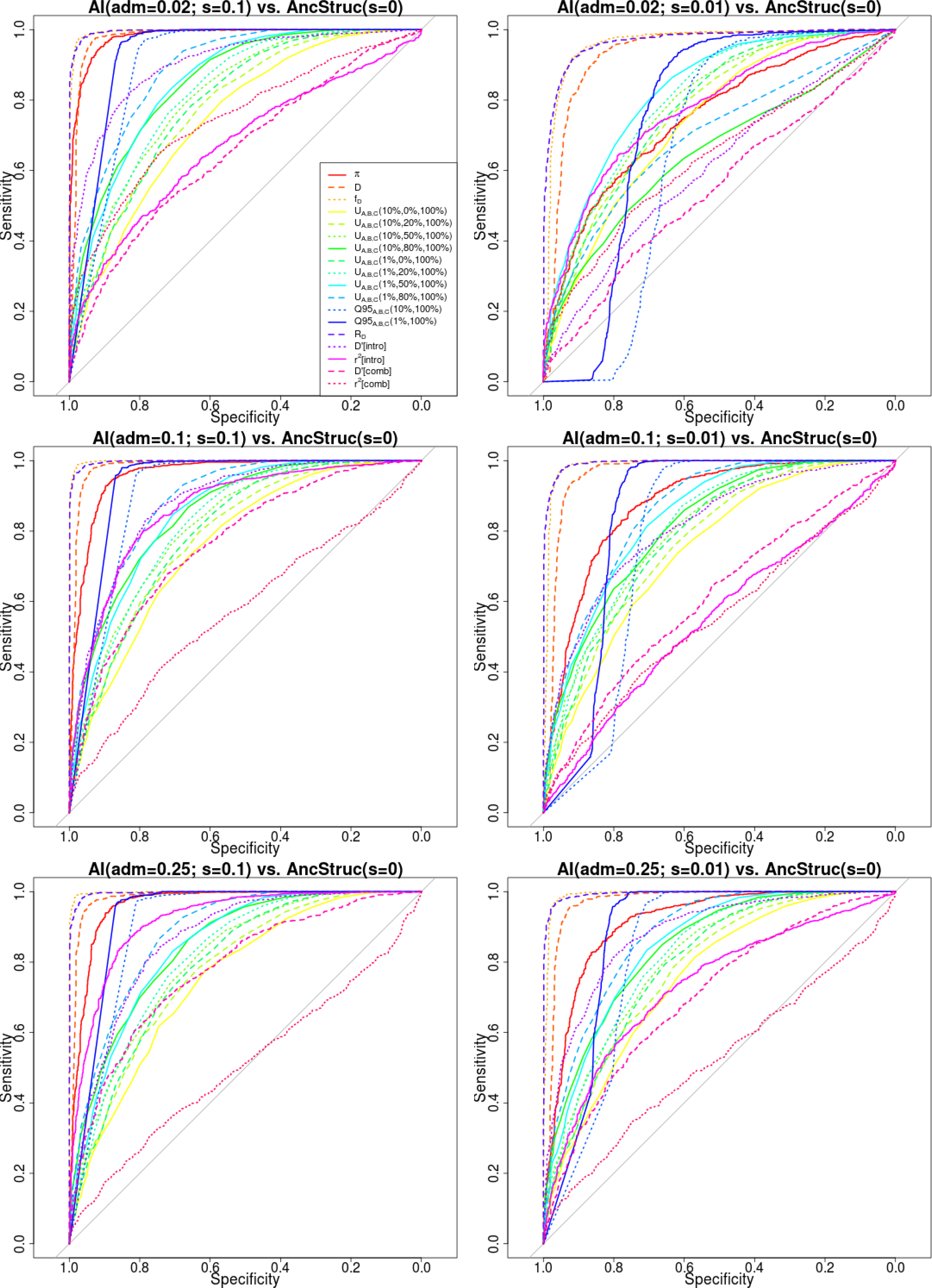
Receiving operating characteristic curves for adaptive introgression against a neutral ancestral structure model with weak migration rates. The demographic scenario for adaptive introgression was the same as in Figure 3. For a description of the ancestral structure model, see main text.

**Figure S20:**
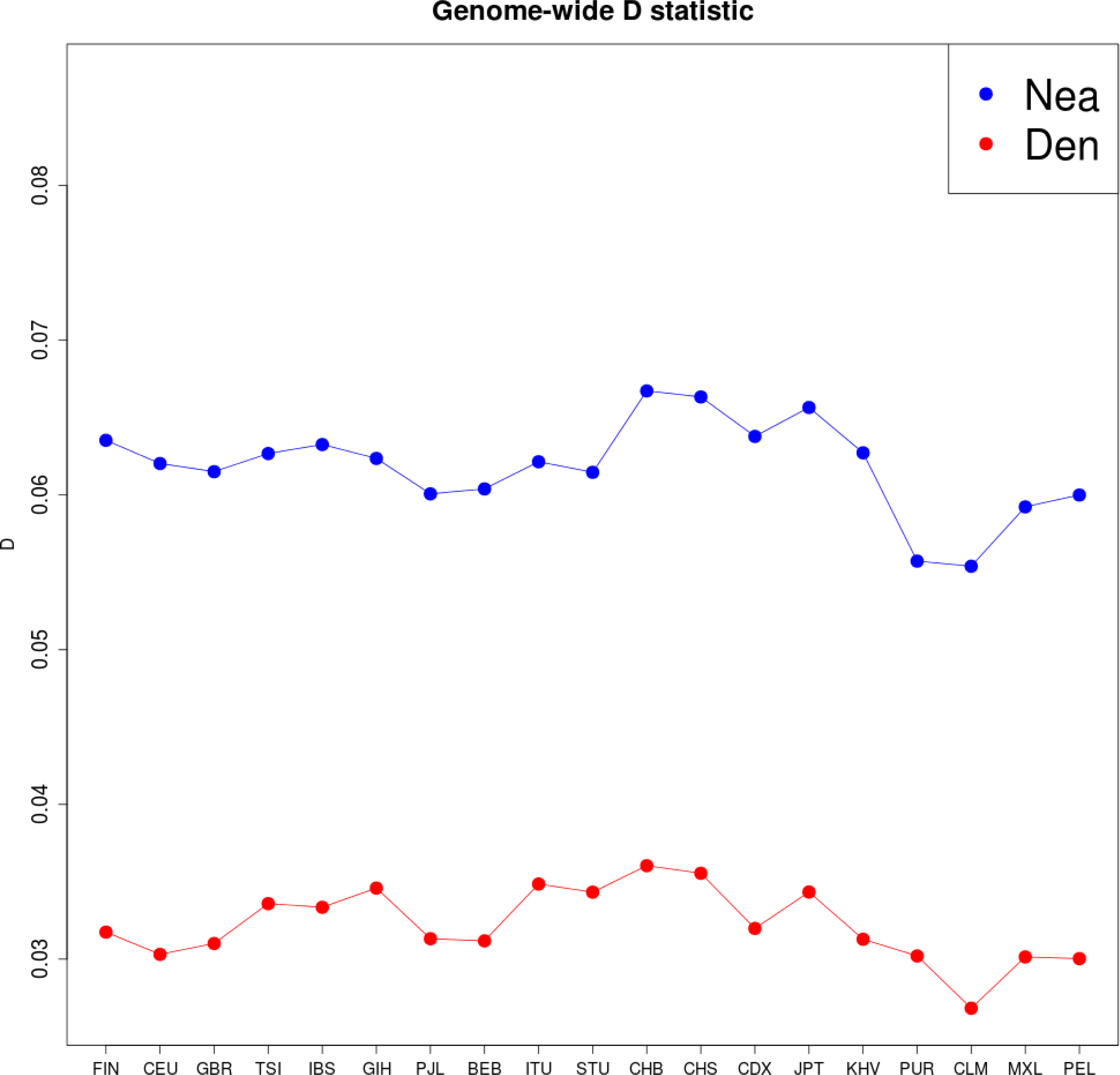
We computed D(X,YRI,Y,Chimpanzee) for different choices of present-day human panels X (x-axis) from phase 3 of the 1000 Genomes Project, and for two high-coverage archaic human genomes Y: Altai Neanderthal (blue) and Denisova (red). The low value of the right-most panel is due to that panel being composed of African-Americans, which have a higher proportion of African ancestry than the other panels.

**Figure S21:**
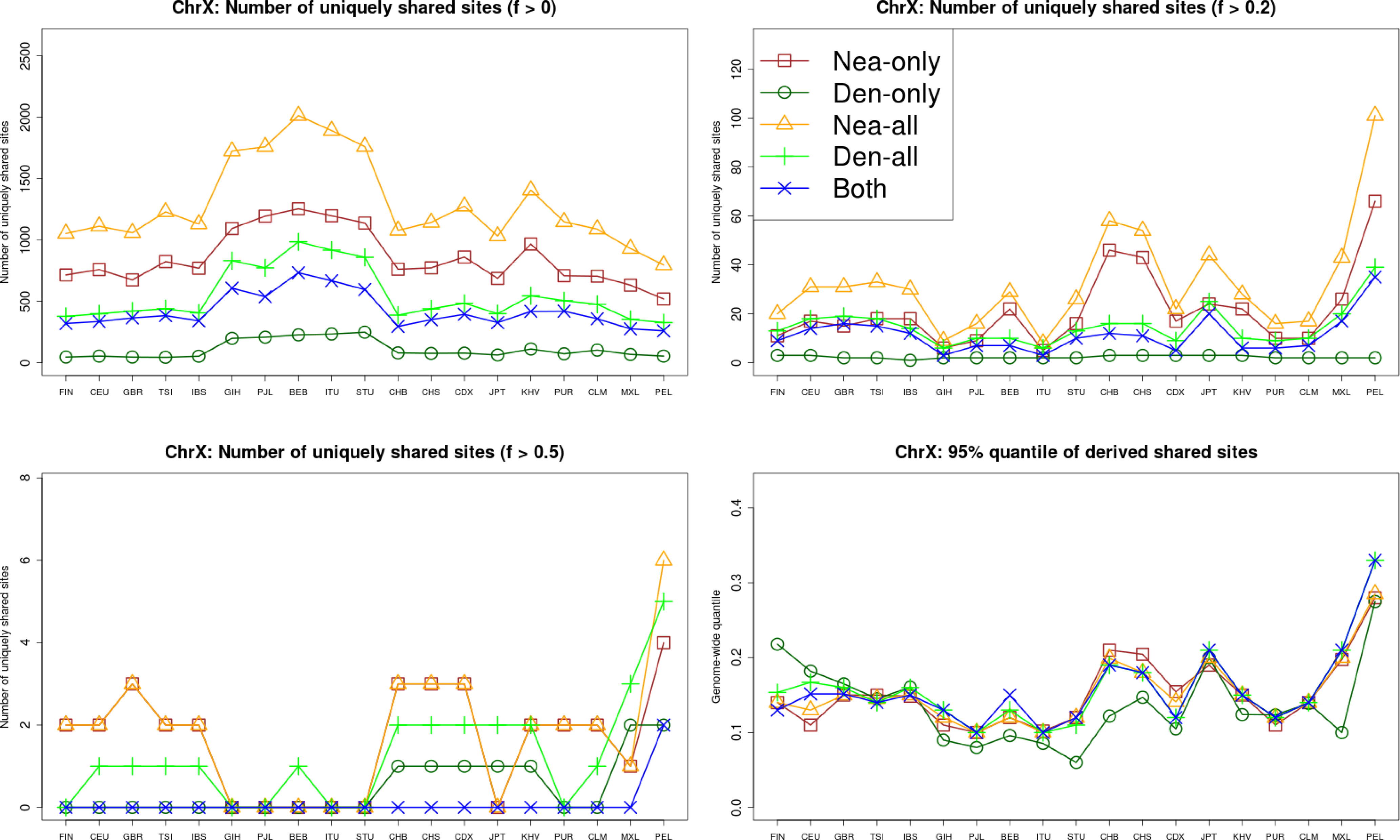
We computed the number of uniquely shared sites in the X chromosome between particular archaic humans genomes and different choices of present-day non-African human panels X (x-axis) from phase 3 of the 1000 Genomes Project, using a shared frequency cutoff of 0% (top-left panel), 20% (top-right panel) and 50% (bottom-left panel). Nea-only = *U_Af r,X,Nea,Den_* (1%, 20%, 100%, 0%). Den-only = *U_Af r,X,Nea,Den_* (1%, 20%, 0%, 100%). Nea-all = *U_Af r,X,Nea_* (1%, 20%, 100%). Den-all = *U_Af r,X,Den_* (1%, 20%, 100%). Both = *U_Af r,X,Nea,Den_* (1%, 20%, 100%, 100%). We also computed the quantile statistics Q95 for different choices of present-day non-African human panels (x-axis) from phase 3 of the 1000 Genomes Project (bottom-right panel). Nea-only = *Q*95_*Af r,X,Nea,Den*_ (1%, 100%, 0%). Den-only = *Q*95_*Af r,X,Nea,Den*_ (1%, 0%, 100%). Nea-all = *Q*95_*Af r,X,Nea*_ (1%, 100%). Den-all = *Q*95_*Af r,X,Den*_ (1%, 50%, 100%). Both = *Q*95_*Af r,X,Nea,Den*_ (1%, 100%, 100%).

**Figure S22:**
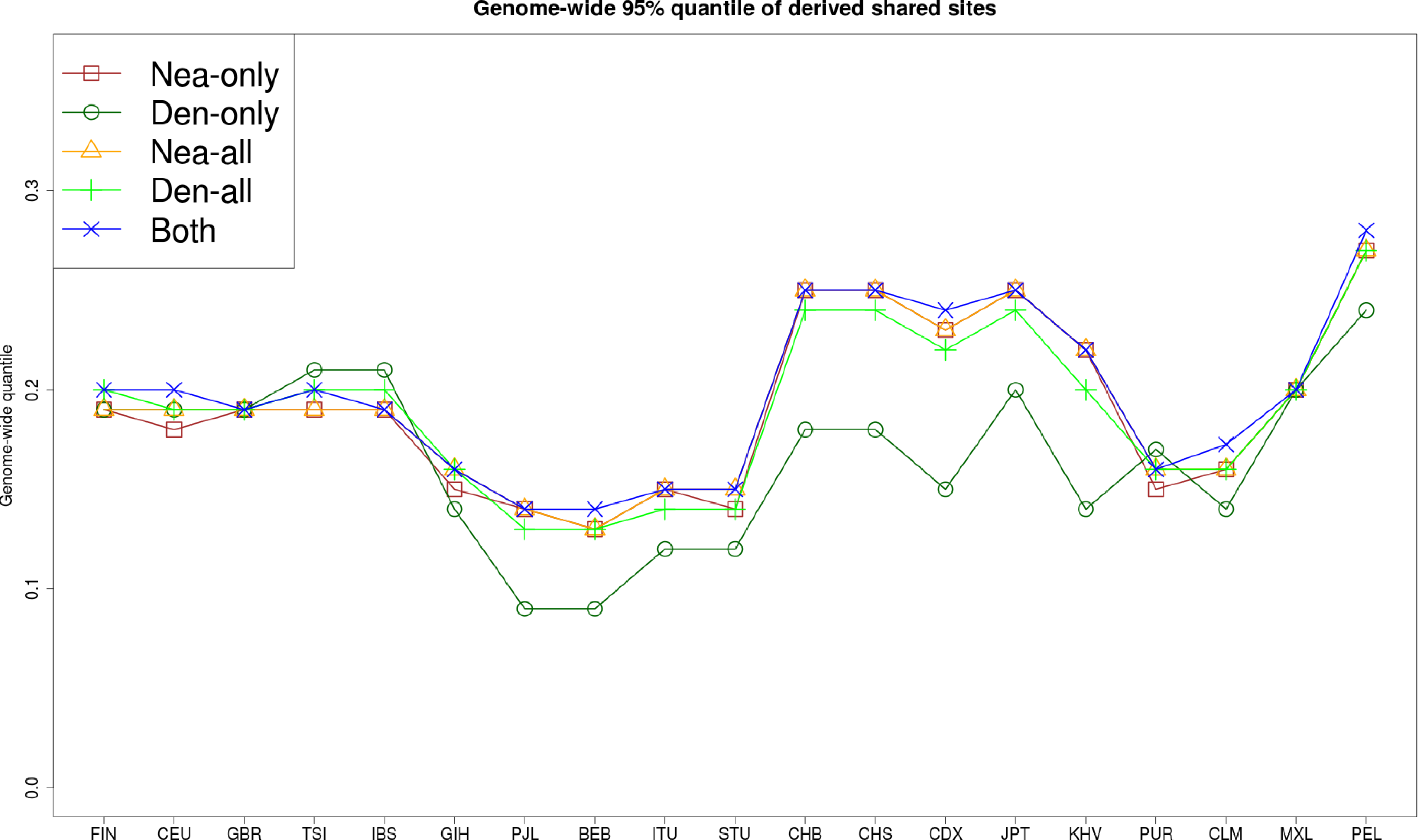
We computed the quantile statistics *Q*95 for different choices of present-day non-African human panels (x-axis) from phase 3 of the 1000 Genomes Project (D). Nea-only = *Q*95_*Af r,X,Nea,Den*_(1%, 100%, 0%). Den-only = *Q*95_*Af r,X,Nea,Den*_(1%, 0%, 100%). Nea-all = *Q*95_*Af r,X,Nea*_(1%, 100%). Den-all = *Q*95_*Af r,X,Den*_ (1%, 100%). Both = *Q*95_*Af r,X,Nea,Den*_ (1%, 100%, 100%).

**Figure S23:**
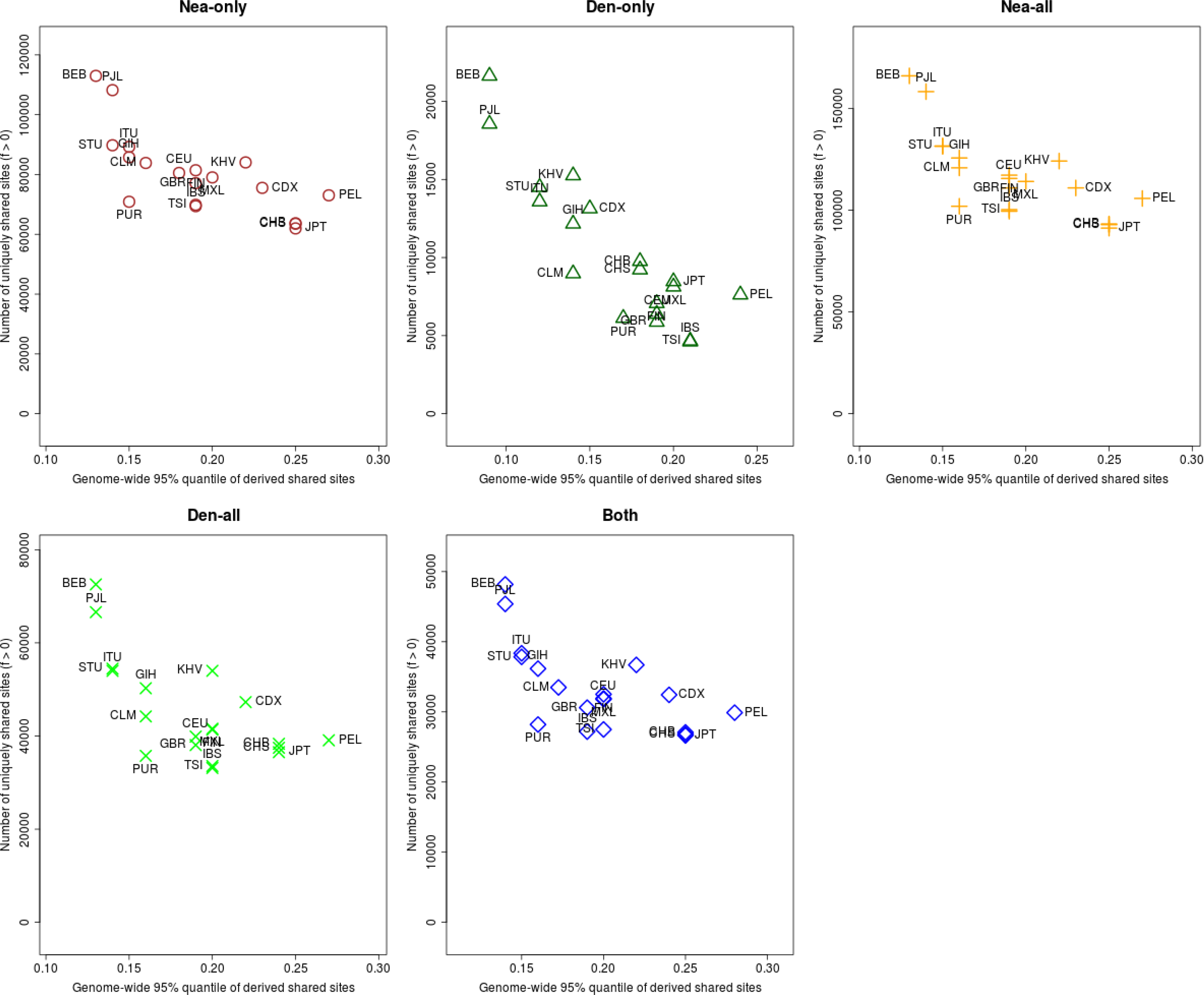
For each population panel from the 1000 Genomes Project, we jointly plotted the *U* and *Q*95 statistics with an archaic frequency cutoff of > 0% within each population. Nea-only = *U_Af r,X,Nea,Den_*(1%, 0%, 100%, 0%) and *Q*95_*Af r,X,Nea,Den*_(1%, 100%, 0%). Den-only = *U_Af r,X,Nea,Den_*(1%, 0%, 0%, 100%) and *Q*95_*Af r,X,Nea,Den*_(1%, 0%, 100%). Nea-all = *U_Af r,X,Nea_* (1%, 0%, 100%) and *Q*95_*Af r,X,Nea*_(1%, 100%). Den-all = *U_Af r,X,Den_* (1%, 0%, 100%) and *Q*95_*Af r,X,Den*_ (1%, 100%). Both = *U_Af r,X,Nea,Den_*(1%, 0%, 100%, 100%) and *Q*95_*Af r,X,Nea,Den*_(1%, 100%, 100%).

**Figure S24:**
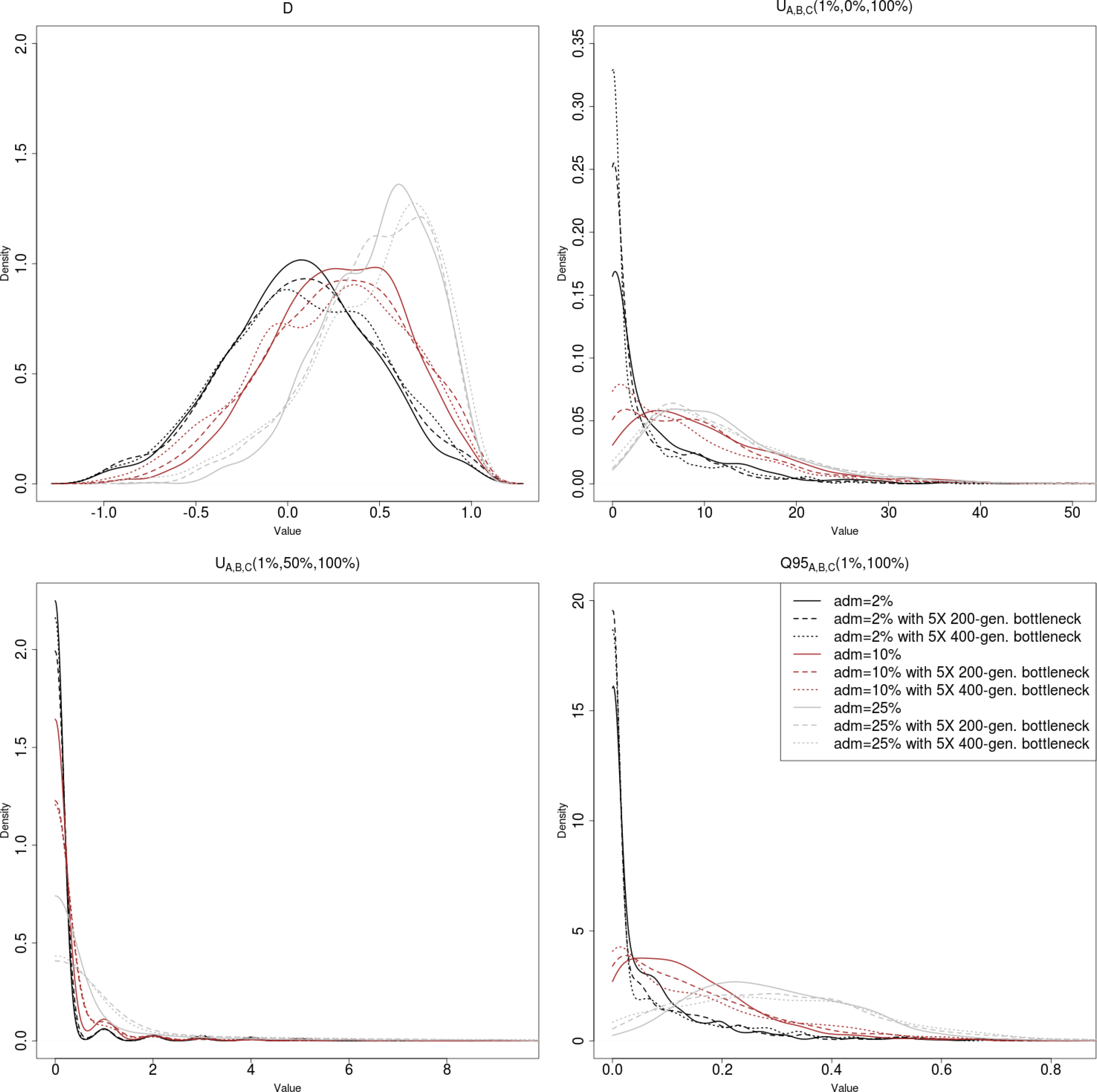
Effect of bottlenecks after the admixture event on the distribution of various statistics under introgression and neutrality.

**Figure S25:**
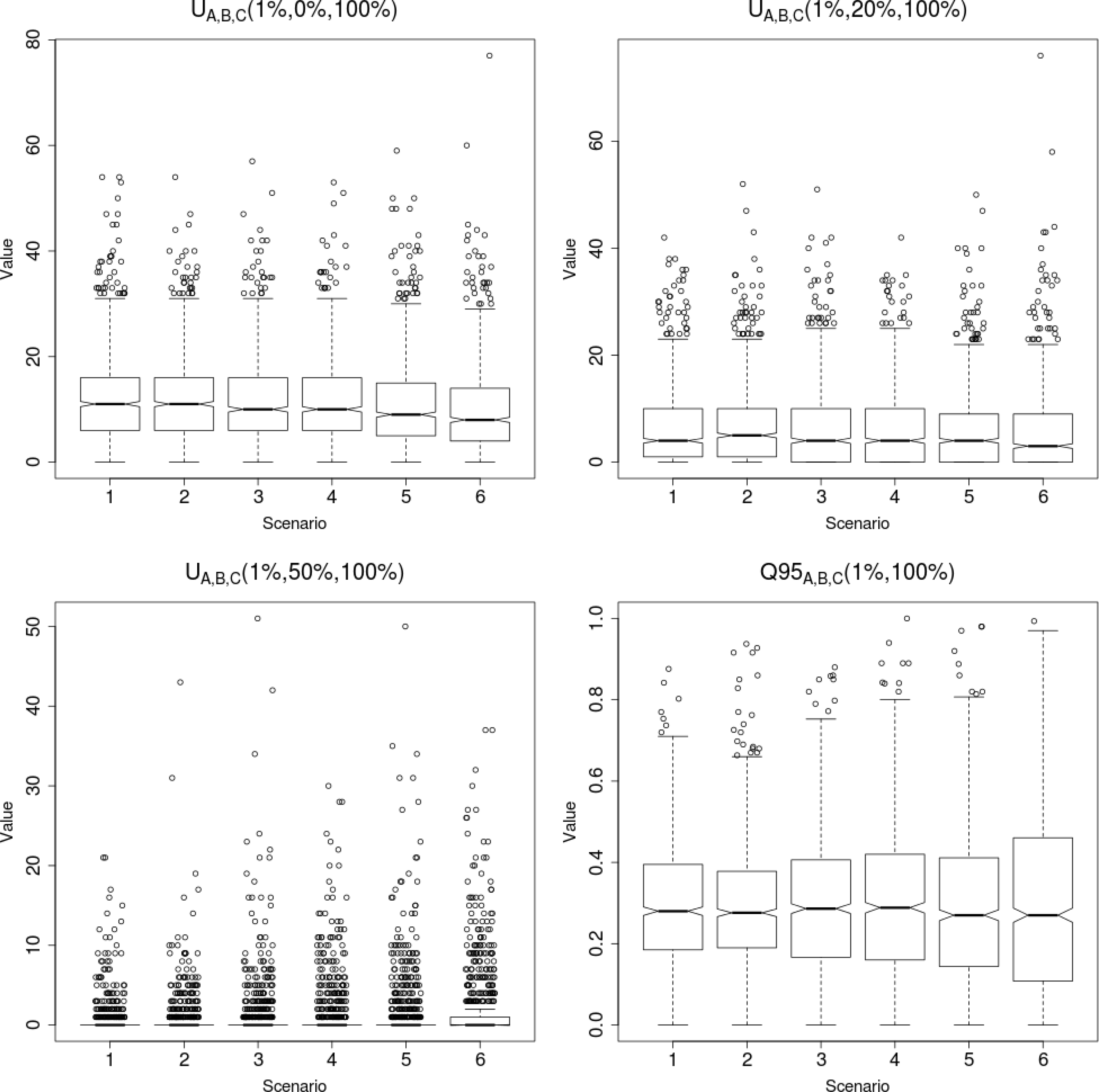
Boxplots showing the effect of different types of bottlenecks on the distribution of *U* and *Q*95 under neutrality. We performed 1,000 simulations for each of 6 different 3-population scenarios with 25% admixture from population C into population B, following the models described in Figure 2. Scenario 1: Constant population size (Figure 2.A). Scenario 2: Pre-admixture 5X bottleneck for 200 generations (Figure 2.B). Scenario 3: Post-admixture 5X bottleneck for 200 generations (Figure 2.C). Scenario 4: Post-admixture 5X bottleneck for 400 generations. Scenario 5: Post-admixture 10X bottleneck for 200 generations. Scenario 6: Post-admixture 10X bottleneck for 400 generations.

**Figure S26:**
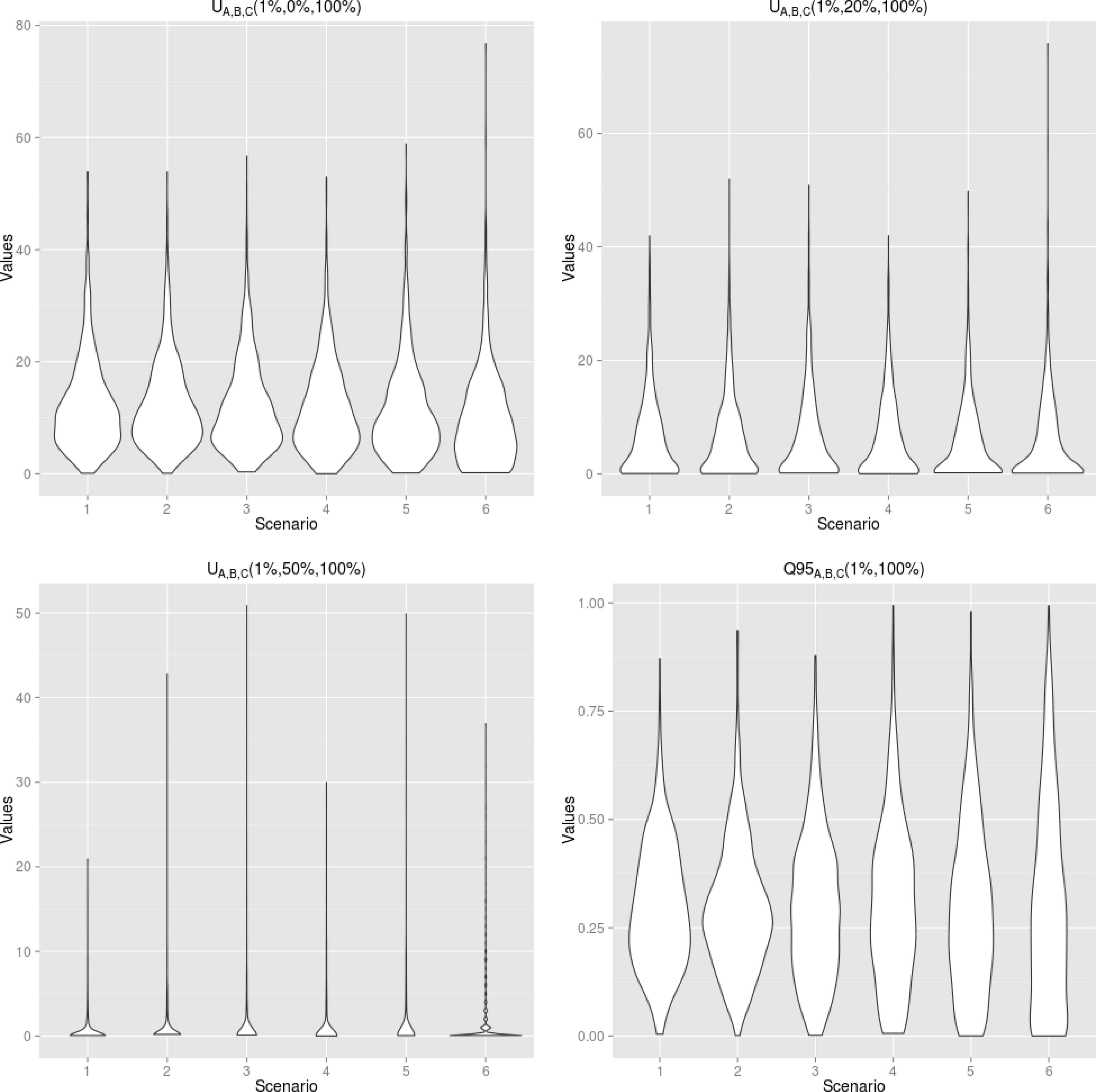
Violin plots showing the effect of different types of bottlenecks on the distribution of *U* and *Q*95 under neutrality (this is the same data as Figure S25). We performed 1,000 simulations for each of 6 different 3-population scenarios with 25% admixture from population C into population B, following the models described in Figure 2. Scenario 1: Constant population size (Figure 2.A). Scenario 2: Pre-admixture 5X bottleneck for 200 generations (Figure 2.B). Scenario 3: Post-admixture 5X bottleneck for 200 generations (Figure 2.C). Scenario 4: Post-admixture 5X bottleneck for 400 generations. Scenario 5: Post-admixture 10X bottleneck for 200 generations. Scenario 6: Post-admixture 10X bottleneck for 400 generations.

**Figure S27:**
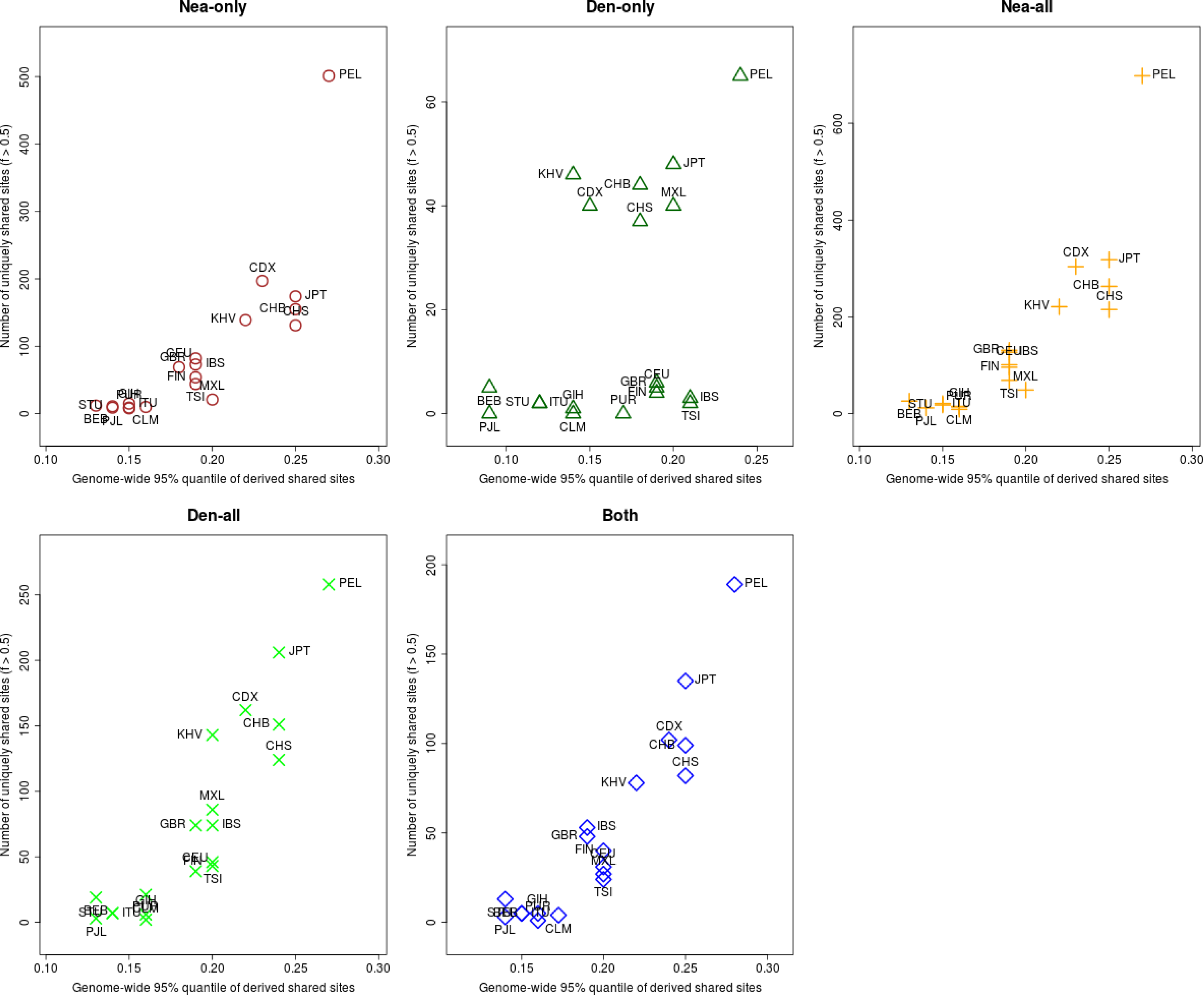
For each population panel from the 1000 Genomes Project, we jointly plotted the *U* and *Q*95 statistics with an archaic frequency cutoff of > 50% within each population. Nea-only = *U_Af r,X,Nea,Den_*(1%, 50%, 100%, 0%) and *Q*95_*Af r,X,Nea,Den*_(1%, 100%, 0%). Den-only = *U_Af r,X,Nea,Den_*(1%, 50%, 0%, 100%) and *Q*95_*Af r,X,Nea,Den*_(1%, 0%, 100%). Nea-all = *U_Af r,X,Nea_*(1%, 50%, 100%) and *Q*95_*Af r,X,Nea*_(1%, 100%). Den-all = *U_Af r,X,Den_*(1%, 50%, 100%) and *Q*95_*Af r,X,Den*_ (1%, 100%). Both = *U_Af r,X,Nea,Den_* (1%, 50%, 100%, 100%) and *Q*95_*Af r,X,Nea,Den*_(1%, 100%, 100%).

**Figure S28:**
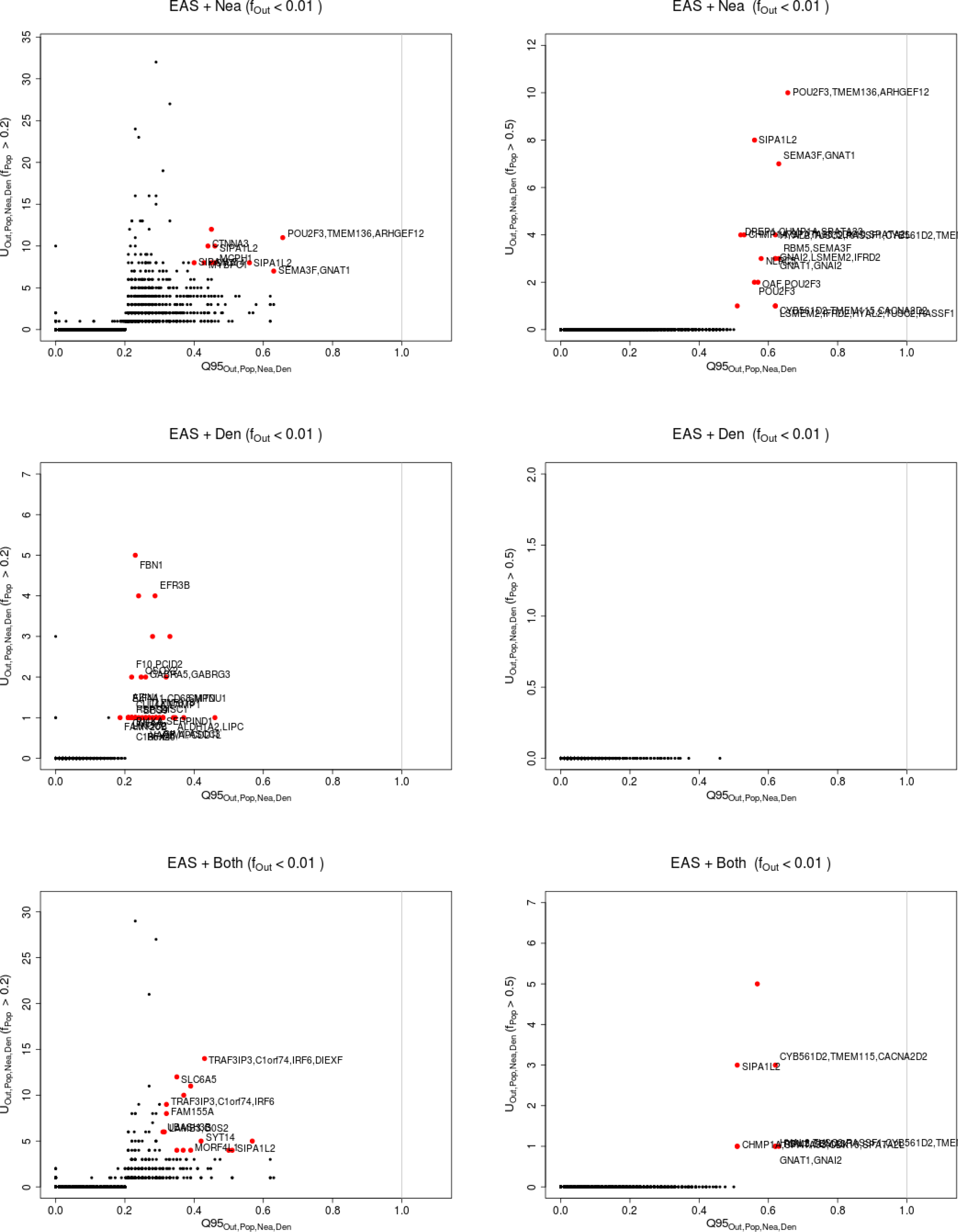
Uniquely shared archaic alleles in an East Asian (EAS) panel. Joint distribution of *Q*95_*EUR* + *AF R,EAS,Nea,Den*_ (1%, *y, z*) and *U_EUR + AF R,EAS,Nea,Den_* (1%,x,y,z), for 40kb non-overlapping regions along the genome, using two choices of x (20% in left column panels,50% in right column panels). Red dots refer to regions that are in the 99.9% quantiles for both statistics. Neanderthal-specific shared alleles are displayed in the top panels, Denisovan-specific shared alleles are displayed in the middle-row panels, and alleles shared with both archaic human genome are displayed in the bottom panels.

**Figure S29:**
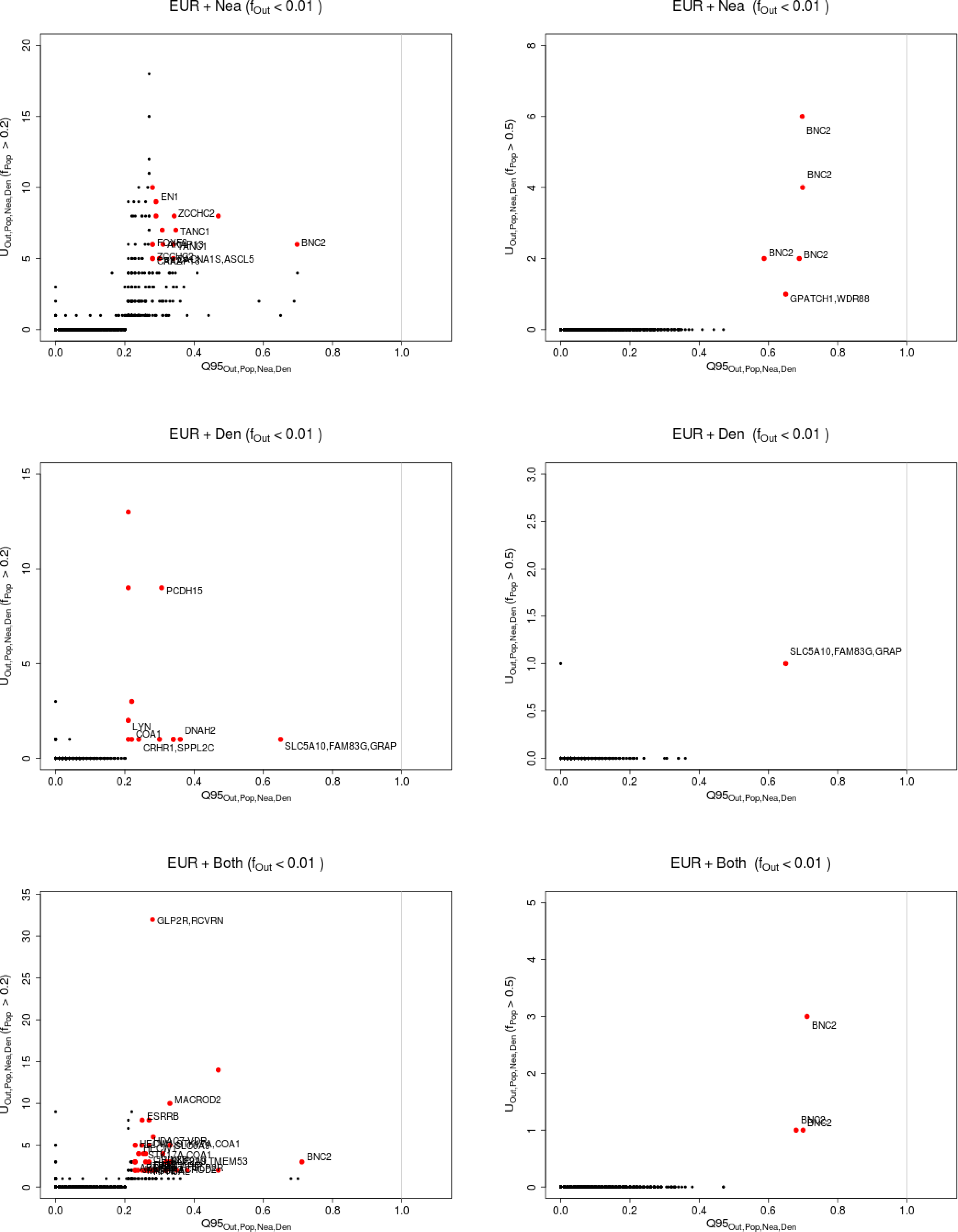
Uniquely shared archaic alleles in an European (EUR) panel. Joint distribution of *Q*95_*EAS* + *AFR,EUR,Nea,Den*_ (1%, *y, z*) and *U_EAS + AFR,EUR,Nea,Den_* (1%,x,y,z), for 40kb non-overlapping regions along the genome, using two choices of x (20% in left column panels,50% in right column panels). Red dots refer to regions that are in the 99.9% quantiles for both statistics. Neanderthal-specific shared alleles are displayed in the top panels, Denisovan-specific shared alleles are displayed in the middle-row panels, and alleles shared with both archaic human genome are displayed in the bottom panels.

**Figure S30:**
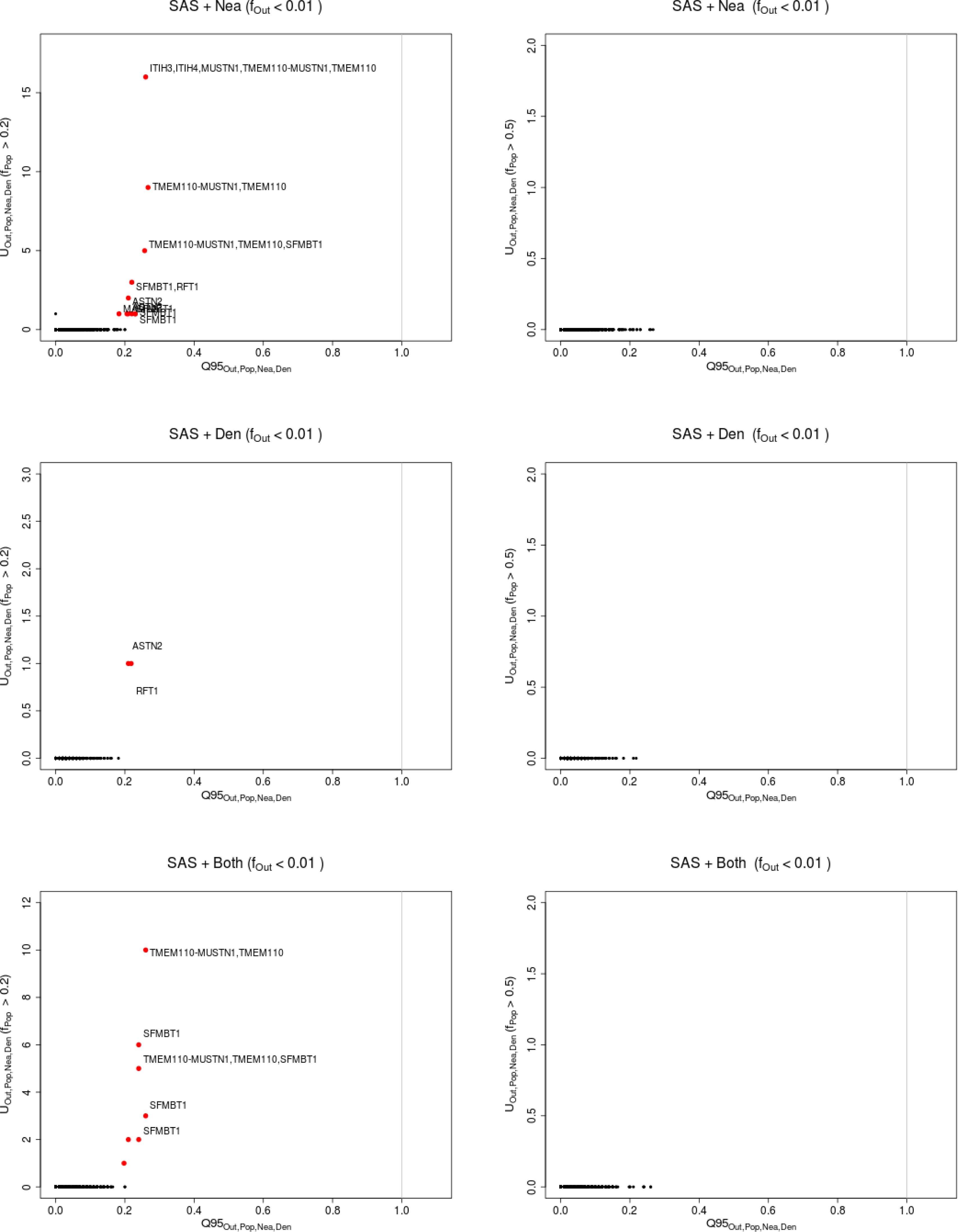
Uniquely shared archaic alleles in a South Asian (SAS) panel. Joint distribution of *Q*95_*EAS + EUR + AFR,SAS,Nea,Den*_ (1%, *y, z*) and *U_EAS + EUR + AF R,SAS,Nea,Den_*(1%,x,y,z), for 40kb non-overlapping regions along the genome, using two choices of x (20% in left column panels,50% in right column panels). Red dots refer to regions that are in the 99.9% quantiles for both statistics. Neanderthal-specific shared alleles are displayed in the top panels, Denisovan-specific shared alleles are displayed in the middle-row panels, and alleles shared with both archaic human genome are displayed in the bottom panels.

**Figure S31:**
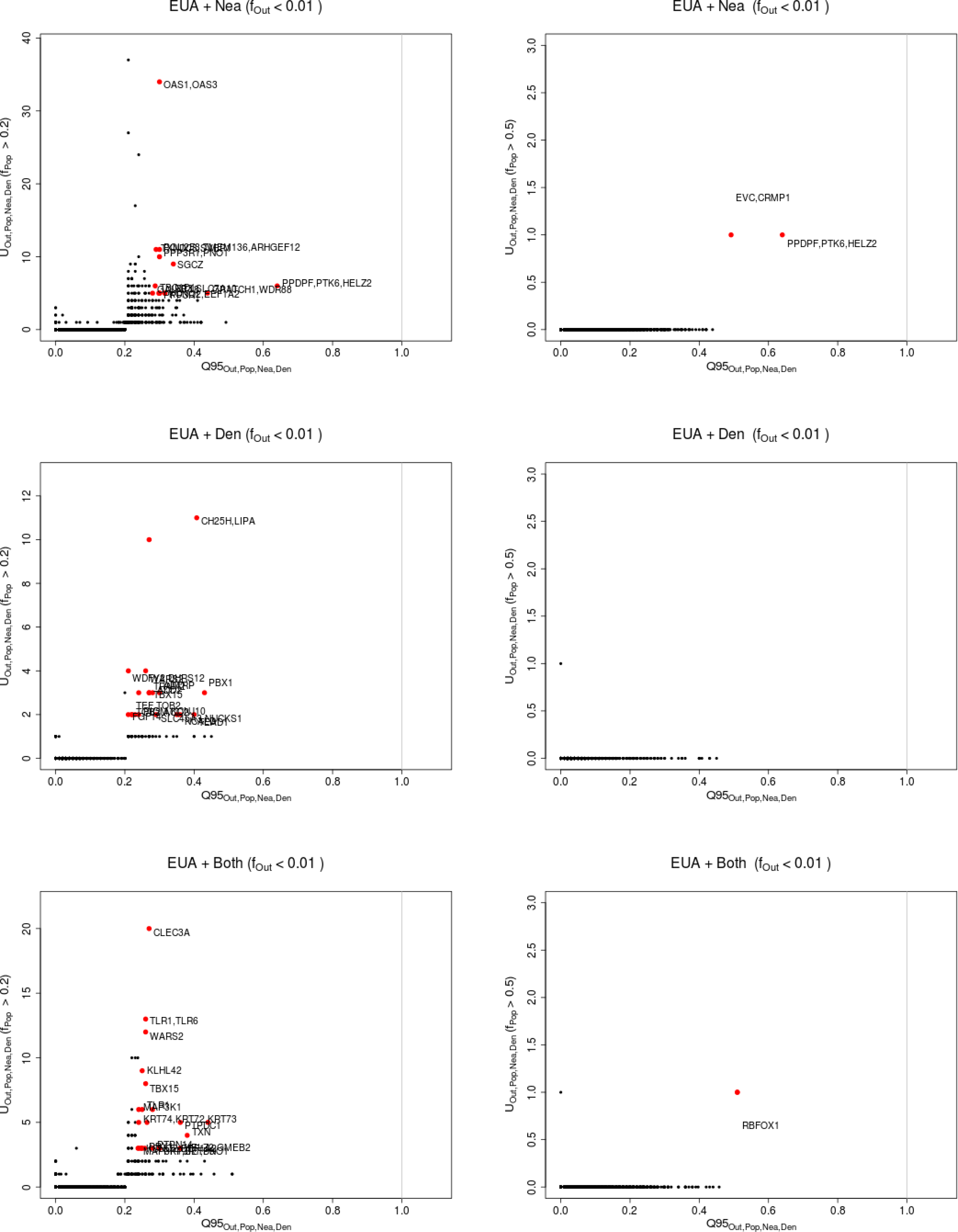
Uniquely shared archaic alleles in a Eurasian (EUA=EUR + SAS + EAS) panel. Joint distribution of *Q*95_*AFR,EUR* + *SAS* + *EAS,Nea,Den*_ (1%, *y, z*) and *U_AFR,EUR + SAS + EAS,Nea,Den_*(1%,x,y,z), for 40kb non-overlapping regions along the genome, using two choices of x (20% in left column panels,50% in right column panels). Red dots refer to regions that are in the 99.9% quantiles for both statistics. Neanderthal-specific shared alleles are displayed in the top panels, Denisovan-specific shared alleles are displayed in the middle-row panels, and alleles shared with both archaic human genome are displayed in the bottom panels.

**Figure S32:**
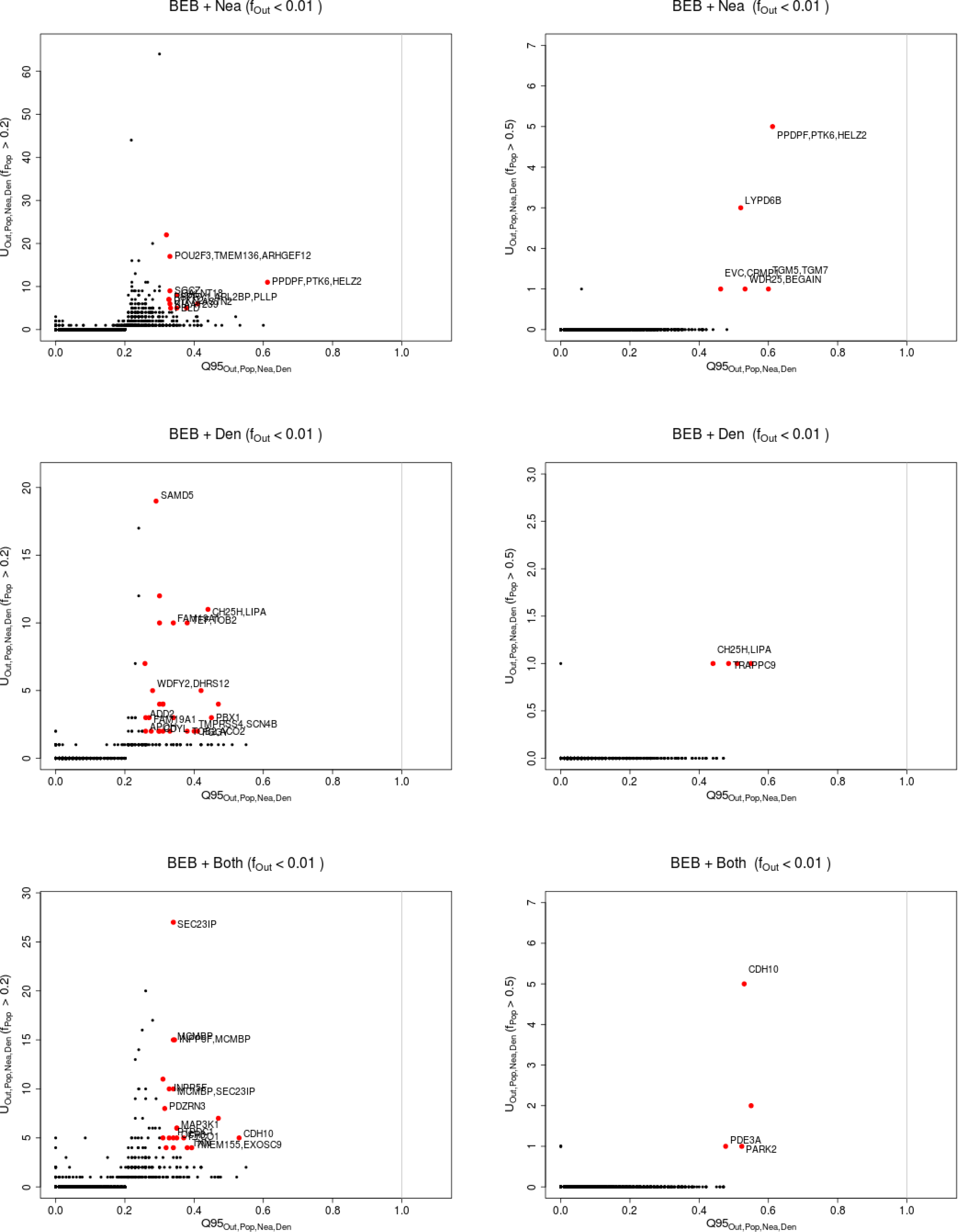
Uniquely shared archaic alleles in a Bengali (BEB) panel. Joint distribution of *Q*95_*AF R,BEB,Nea,Den*_ (1%, *y, z*) and *U_AF R,BEB,Nea,Den_*(1%,x,y,z), for 40kb non-overlapping regions along the genome, using two choices of x (20% in left column panels,50% in right column panels). Red dots refer to regions that are in the 99.9% quantiles for both statistics. Neanderthal-specific shared alleles are displayed in the top panels, Denisovan-specific shared alleles are displayed in the middle-row panels, and alleles shared with both archaic human genome are displayed in the bottom panels.

**Figure S33:**
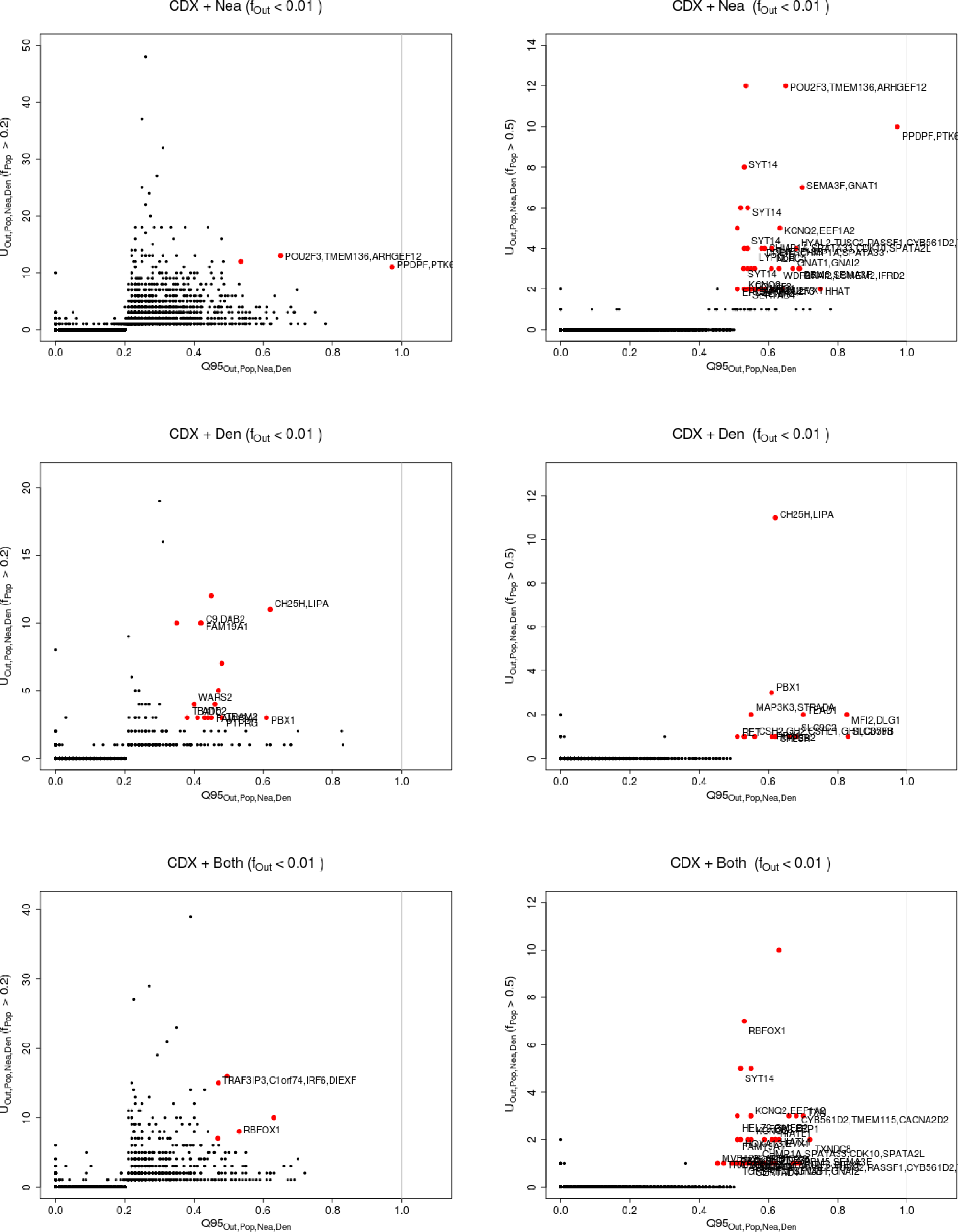
Uniquely shared archaic alleles in a Chinese Dai (CDX) panel. Joint distribution of *Q*95_*AFR,CDX,Nea,Den*_ (1%, *y, z*) and *U_AFR,CDX,Nea,Den_*(1%,x,y,z), for 40kb non-overlapping regions along the genome, using two choices of x (20% in left column panels,50% in right column panels). Red dots refer to regions that are in the 99.9% quantiles for both statistics. Neanderthal-specific shared alleles are displayed in the top panels, Denisovan-specific shared alleles are displayed in the middle-row panels, and alleles shared with both archaic human genome are displayed in the bottom panels.

**Figure S34:**
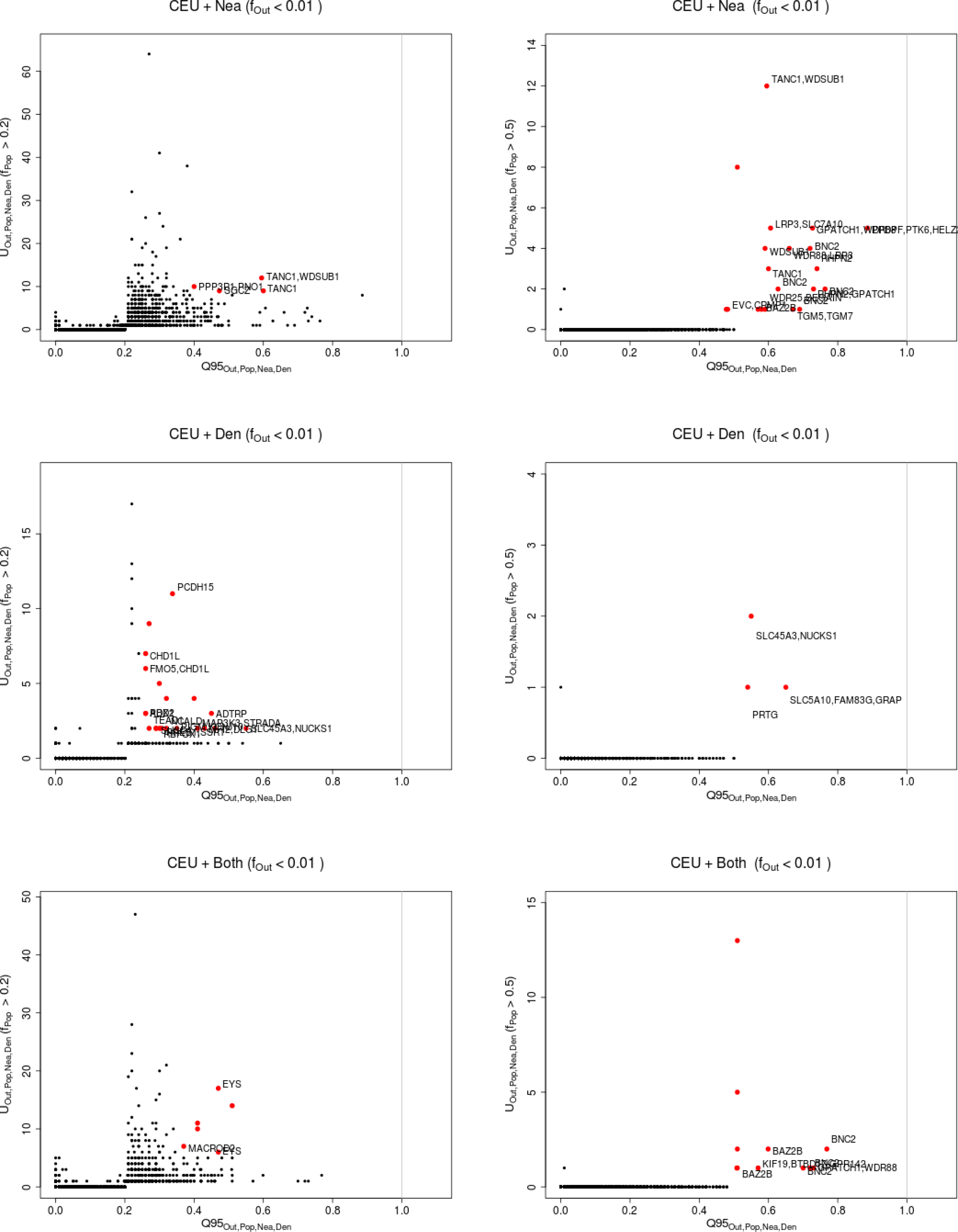
Uniquely shared archaic alleles in a Central European (CEU) panel. Joint distribution of *Q*95_*AFR,CEU,Nea,Den*_(1%, *y, z*) and *U_AFR,CEU,Nea,Den_*(1%,x,y,z), for 40kb non-overlapping regions along the genome, using two choices of x (20% in left column panels,50% in right column panels). Red dots refer to regions that are in the 99.9% quantiles for both statistics. Neanderthal-specific shared alleles are displayed in the top panels, Denisovan-specific shared alleles are displayed in the middle-row panels, and alleles shared with both archaic human genome are displayed in the bottom panels.

**Figure S35:**
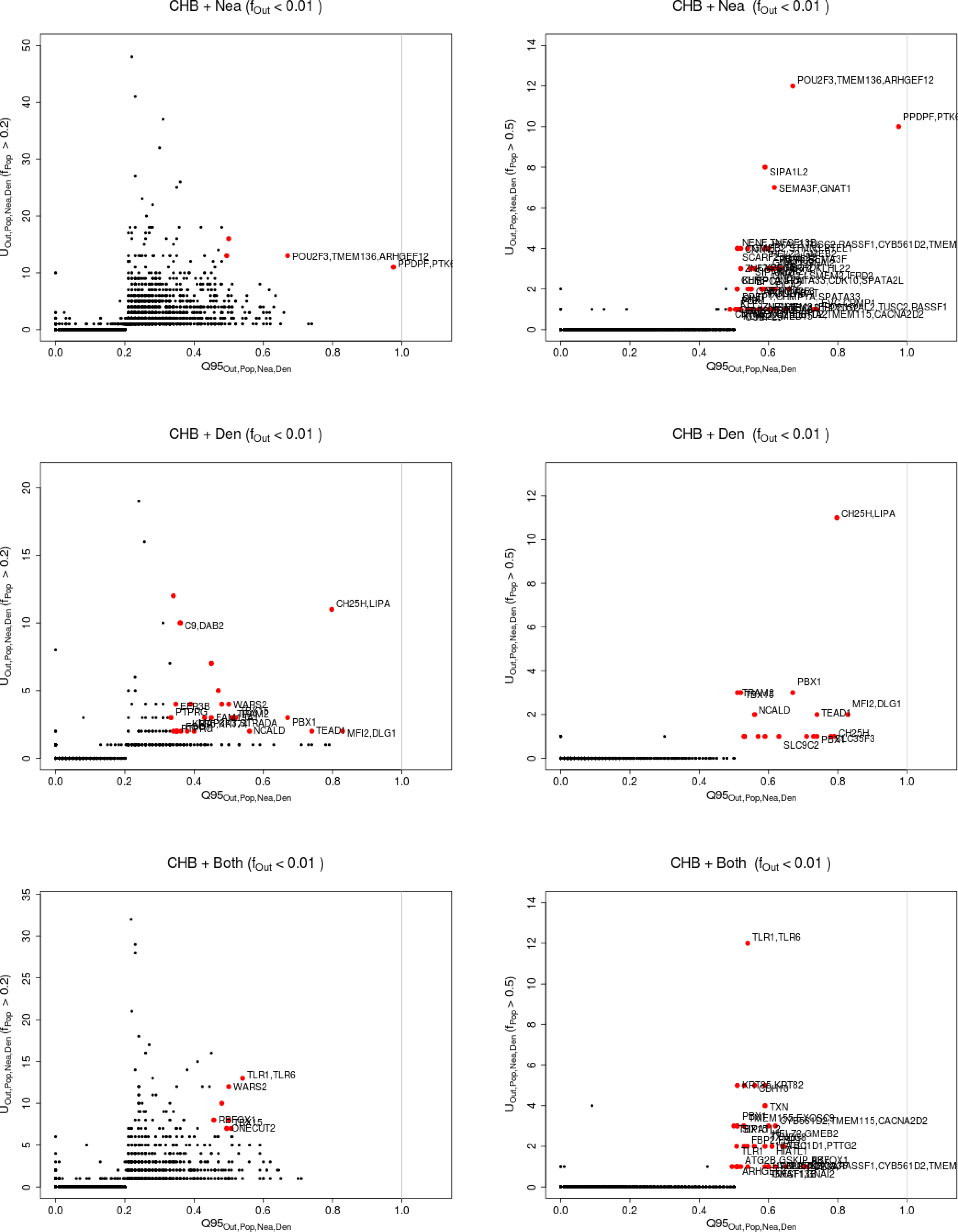
Uniquely shared archaic alleles in a Han Chinese (CHB) panel. Joint distribution of *Q*95_*AFR,CHB,Nea,Den*_(1%, *y, z*) and *U_AFR,CHB,Nea,Den_*(1%,x,y,z), for 40kb non-overlapping regions along the genome, using two choices of x (20% in left column panels,50% in right column panels). Red dots refer to regions that are in the 99.9% quantiles for both statistics. Neanderthal-specific shared alleles are displayed in the top panels, Denisovan-specific shared alleles are displayed in the middle-row panels, and alleles shared with both archaic human genome are displayed in the bottom panels.

**Figure S36:**
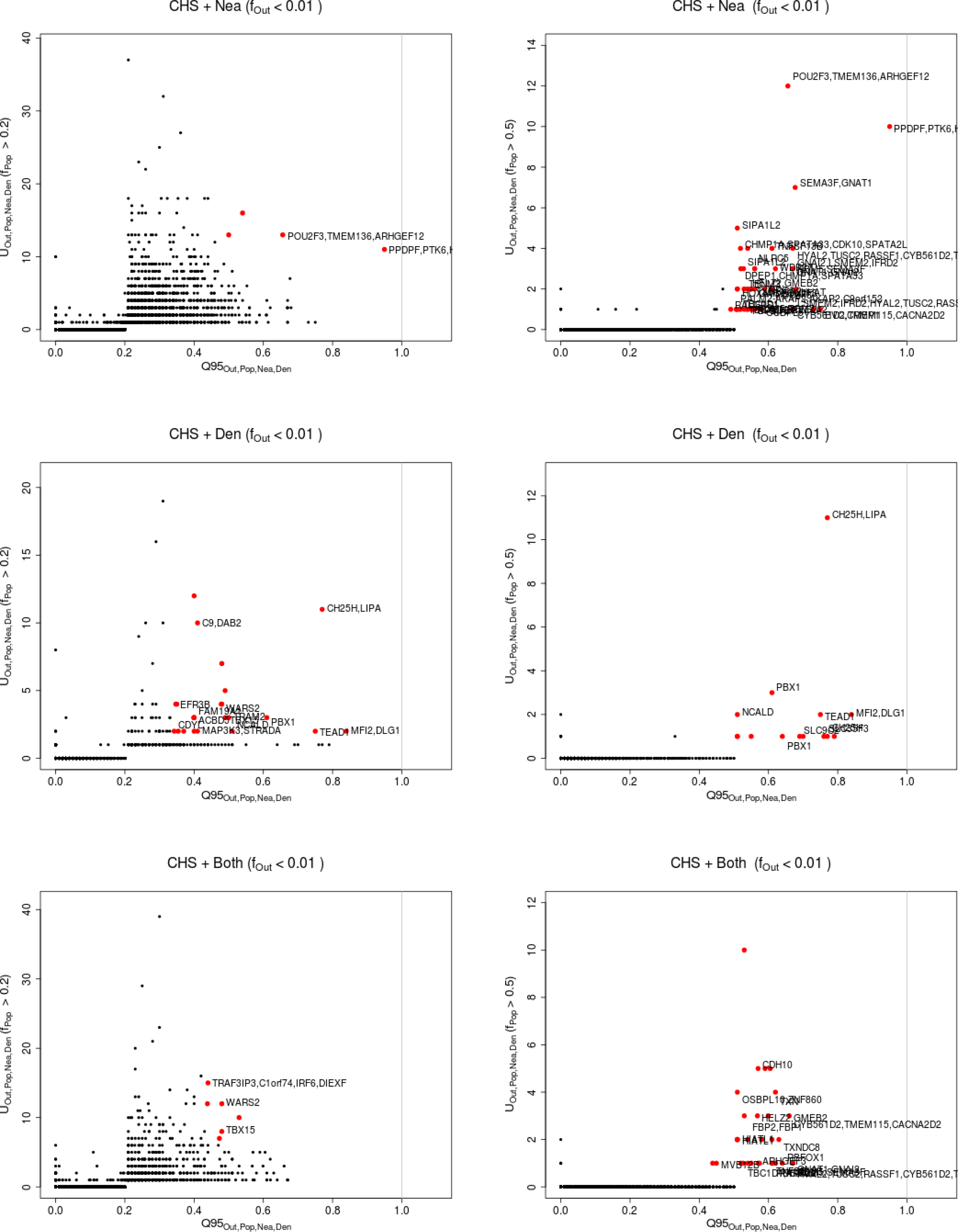
Uniquely shared archaic alleles in a Southern Han Chinese (CHS) panel. Joint distribution of *Q*95_*AF R,BEB,Nea,Den*_(1%, *y, z*) and *U_AFR,CHS,Nea,Den_*(1%,x,y,z), for 40kb non-overlapping regions along the genome, using two choices of x (20% in left column panels,50% in right column panels). Red dots refer to regions that are in the 99.9% quantiles for both statistics. Neanderthal-specific shared alleles are displayed in the top panels, Denisovan-specific shared alleles are displayed in the middle-row panels, and alleles shared with both archaic human genome are displayed in the bottom panels.

**Figure S37:**
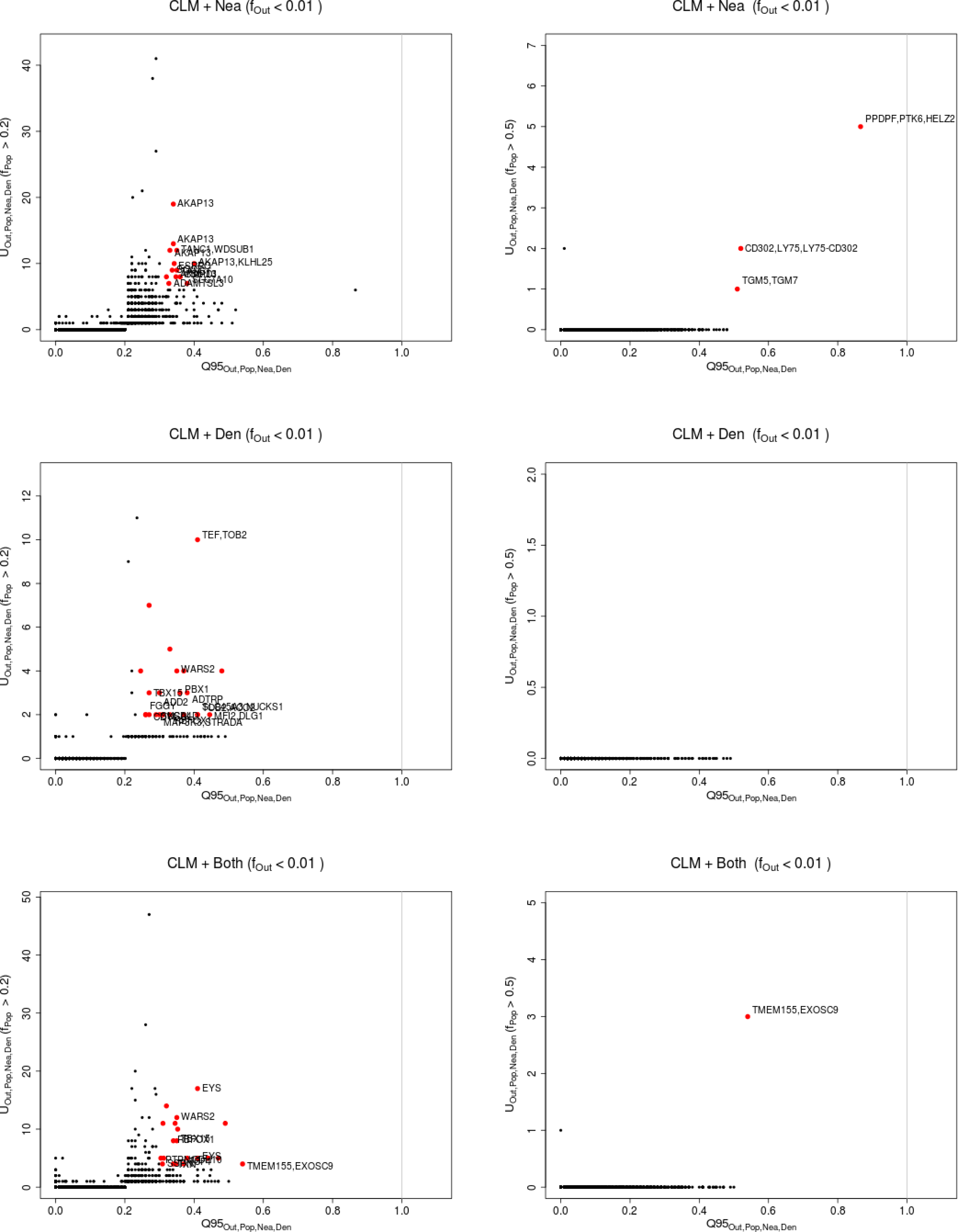
Uniquely shared archaic alleles in a Colombian (CLM) panel. Joint distribution of *Q*95_*AFR,CLM,Nea,Den*_(1%, *y, z*) and *U_AFR,CLM,Nea,Den_*(1%,x,y,z), for 40kb non-overlapping regions along the genome, using two choices of x (20% in left column panels,50% in right column panels). Red dots refer to regions that are in the 99.9% quantiles for both statistics. Neanderthal-specific shared alleles are displayed in the top panels, Denisovan-specific shared alleles are displayed in the middle-row panels, and alleles shared with both archaic human genome are displayed in the bottom panels.

**Figure S38:**
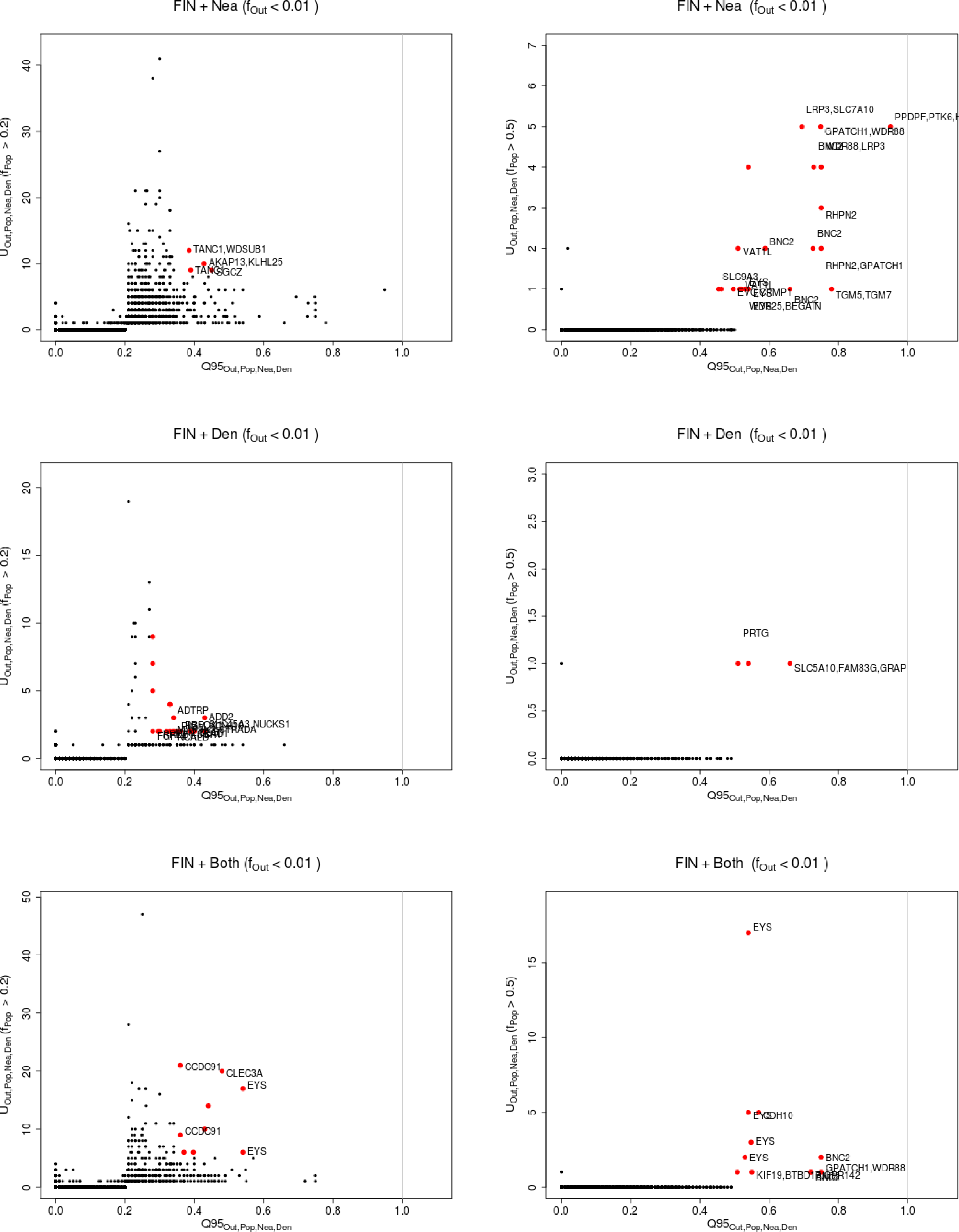
Uniquely shared archaic alleles in a Finnish (FIN) panel. Joint distribution of *Q*95_*AFR,FIN,Nea,Den*_(1%, *y, z*) and *U_AFR,FIN,Nea,Den_*(1%,x,y,z), for 40kb non-overlapping regions along the genome, using two choices of x (20% in left column panels,50% in right column panels). Red dots refer to regions that are in the 99.9% quantiles for both statistics. Neanderthal-specific shared alleles are displayed in the top panels, Denisovan-specific shared alleles are displayed in the middle-row panels, and alleles shared with both archaic human genome are displayed in the bottom panels.

**Figure S39:**
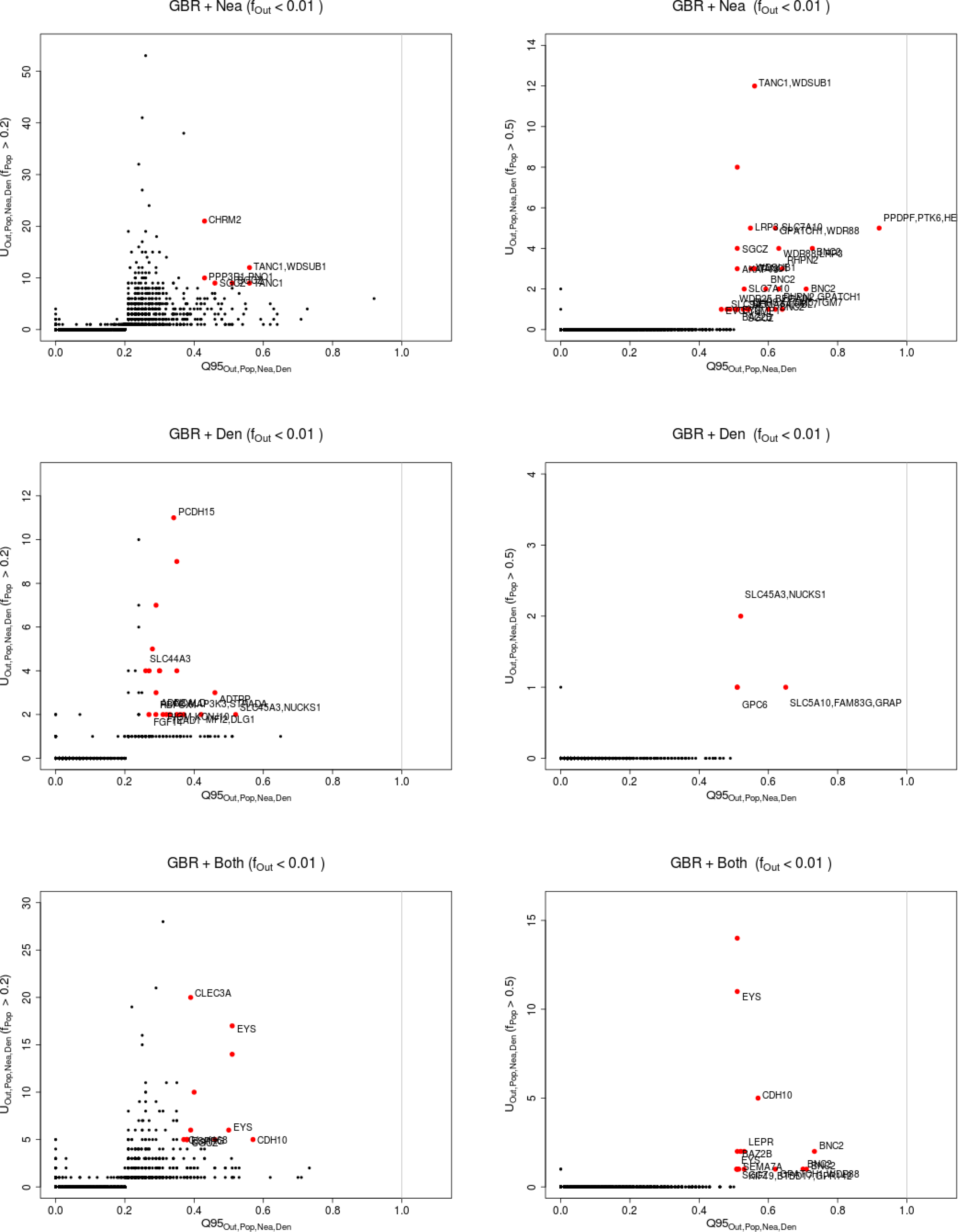
Uniquely shared archaic alleles in a British (GBR) panel. Joint distribution of *Q*95_*AF R,GBR,Nea,Den*_ (1%, *y, z*) and *U_AF R,GBR,Nea,Den_*(1%,x,y,z), for 40kb non-overlapping regions along the genome, using two choices of x (20% in left column panels,50% in right column panels). Red dots refer to regions that are in the 99.9% quantiles for both statistics. Neanderthal-specific shared alleles are displayed in the top panels, Denisovan-specific shared alleles are displayed in the middle-row panels, and alleles shared with both archaic human genome are displayed in the bottom panels.

**Figure S40:**
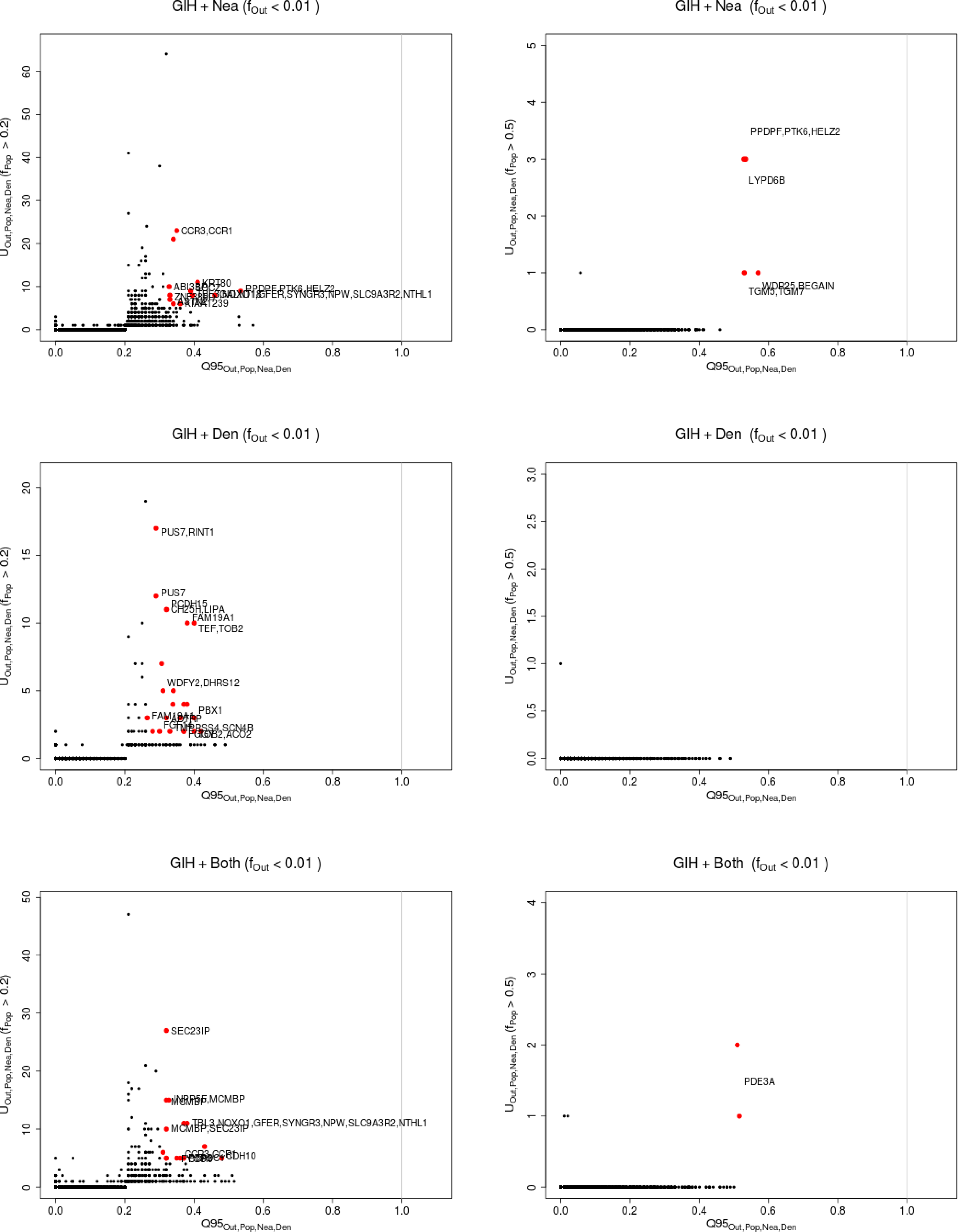
Uniquely shared archaic alleles in a Gujarati Indian (GIH) panel. Joint distribution of *Q*95_*AFR,GIH,Nea,Den*_ (1%, *y, z*) and *U_AFR,GIH,Nea,Den_*(1%,x,y,z), for 40kb non-overlapping regions along the genome, using two choices of x (20% in left column panels,50% in right column panels). Red dots refer to regions that are in the 99.9% quantiles for both statistics. Neanderthal-specific shared alleles are displayed in the top panels, Denisovan-specific shared alleles are displayed in the middle-row panels, and alleles shared with both archaic human genome are displayed in the bottom panels.

**Figure S41:**
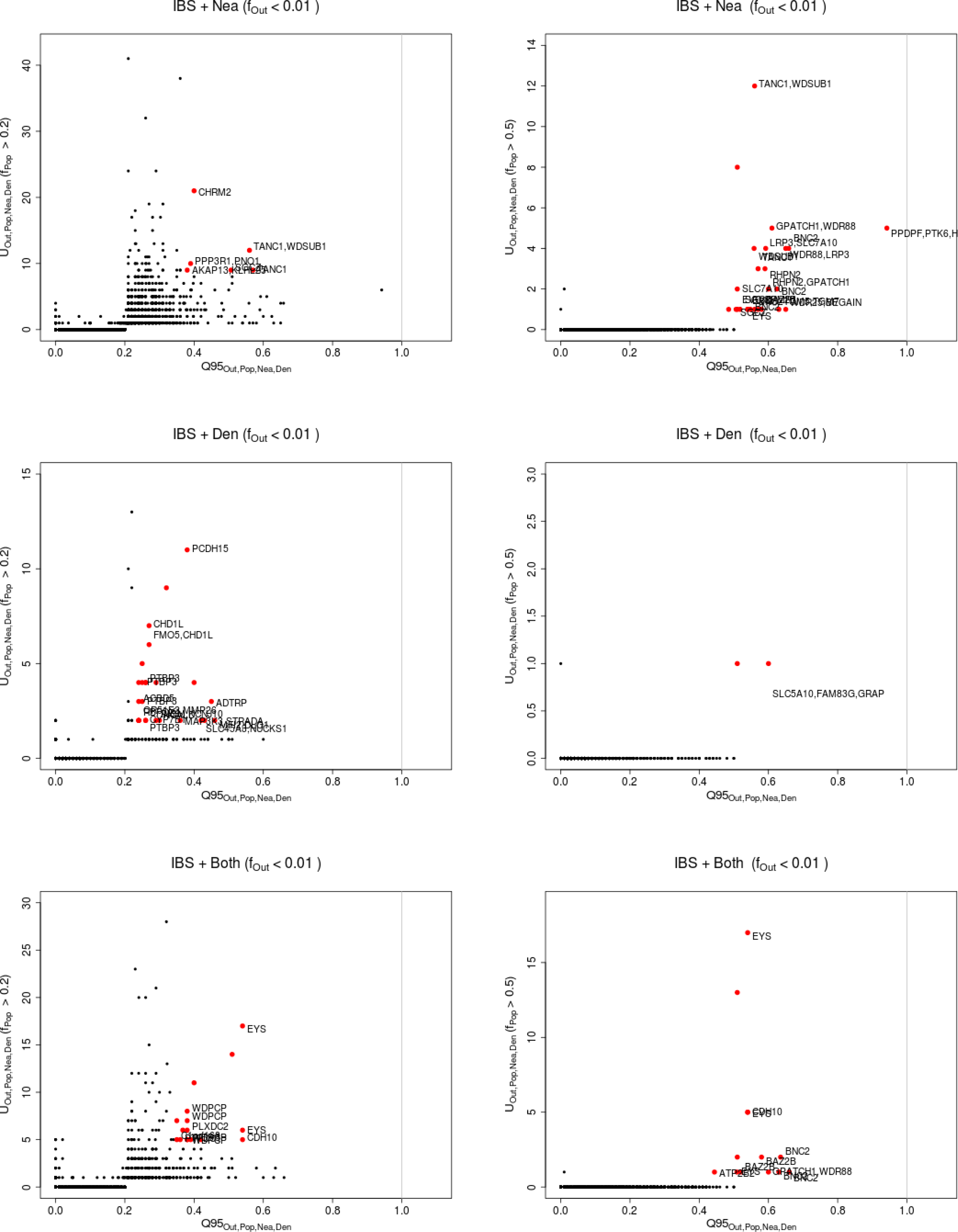
Uniquely shared archaic alleles in an Iberian (IBS) panel. Joint distribution of *Q*95_*AFR,IBS,Nea,Den*_(1%, *y, z*) and *U_AFR,IBS,Nea,Den_*(1%,x,y,z), for 40kb non-overlapping regions along the genome, using two choices of x (20% in left column panels,50% in right column panels). Red dots refer to regions that are in the 99.9% quantiles for both statistics. Neanderthal-specific shared alleles are displayed in the top panels, Denisovan-specific shared alleles are displayed in the middle-row panels, and alleles shared with both archaic human genome are displayed in the bottom panels.

**Figure S42:**
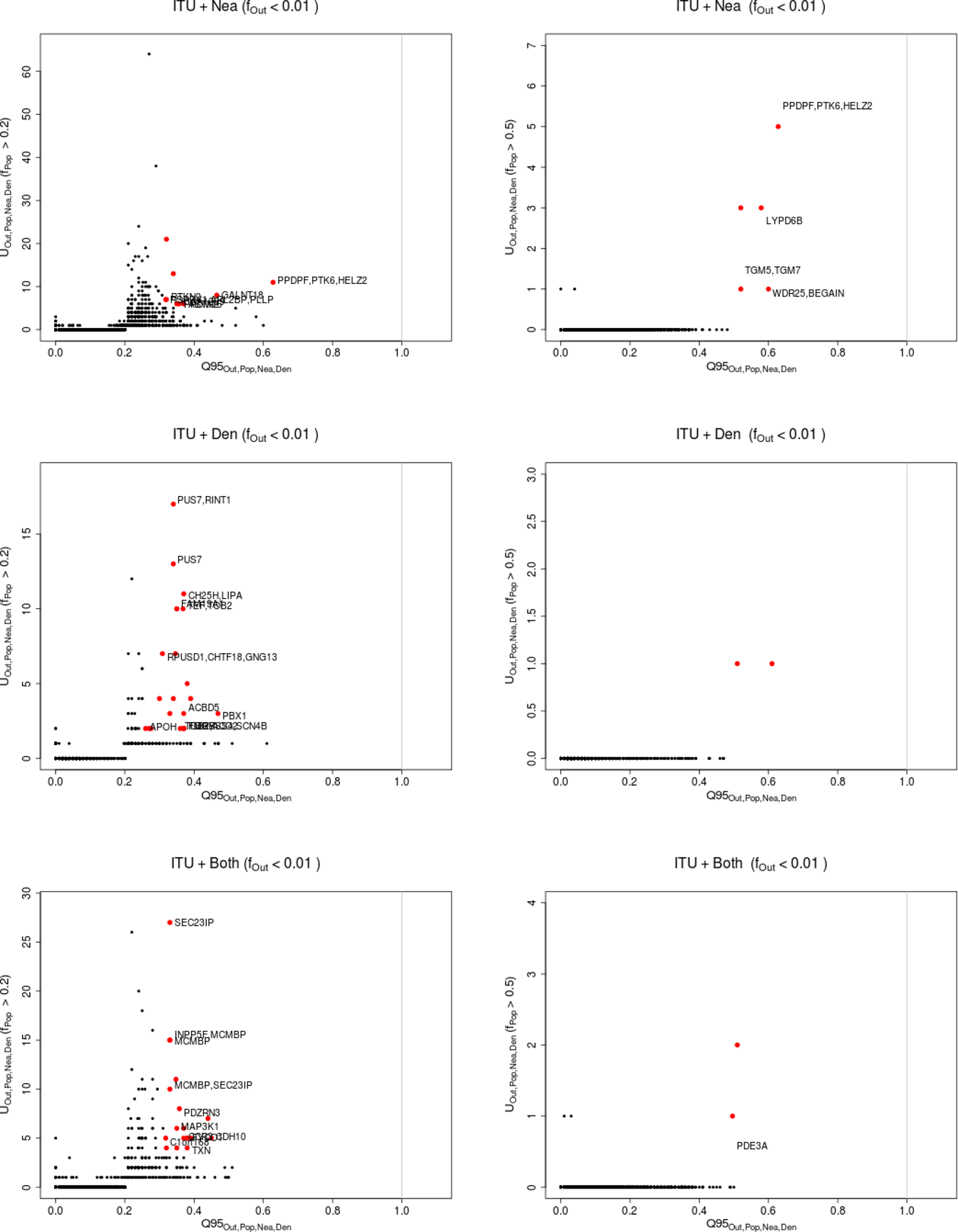
Uniquely shared archaic alleles in an Indian Telugu (ITU) panel. Joint distribution of *Q*95_*AFR,ITU,Nea,Den*_ (1%, *y, z*) and *U_AFR,ITU,Nea,Den_*(1%,x,y,z), for 40kb non-overlapping regions along the genome, using two choices of x (20% in left column panels,50% in right column panels). Red dots refer to regions that are in the 99.9% quantiles for both statistics. Neanderthal-specific shared alleles are displayed in the top panels, Denisovan-specific shared alleles are displayed in the middle-row panels, and alleles shared with both archaic human genome are displayed in the bottom panels.

**Figure S43:**
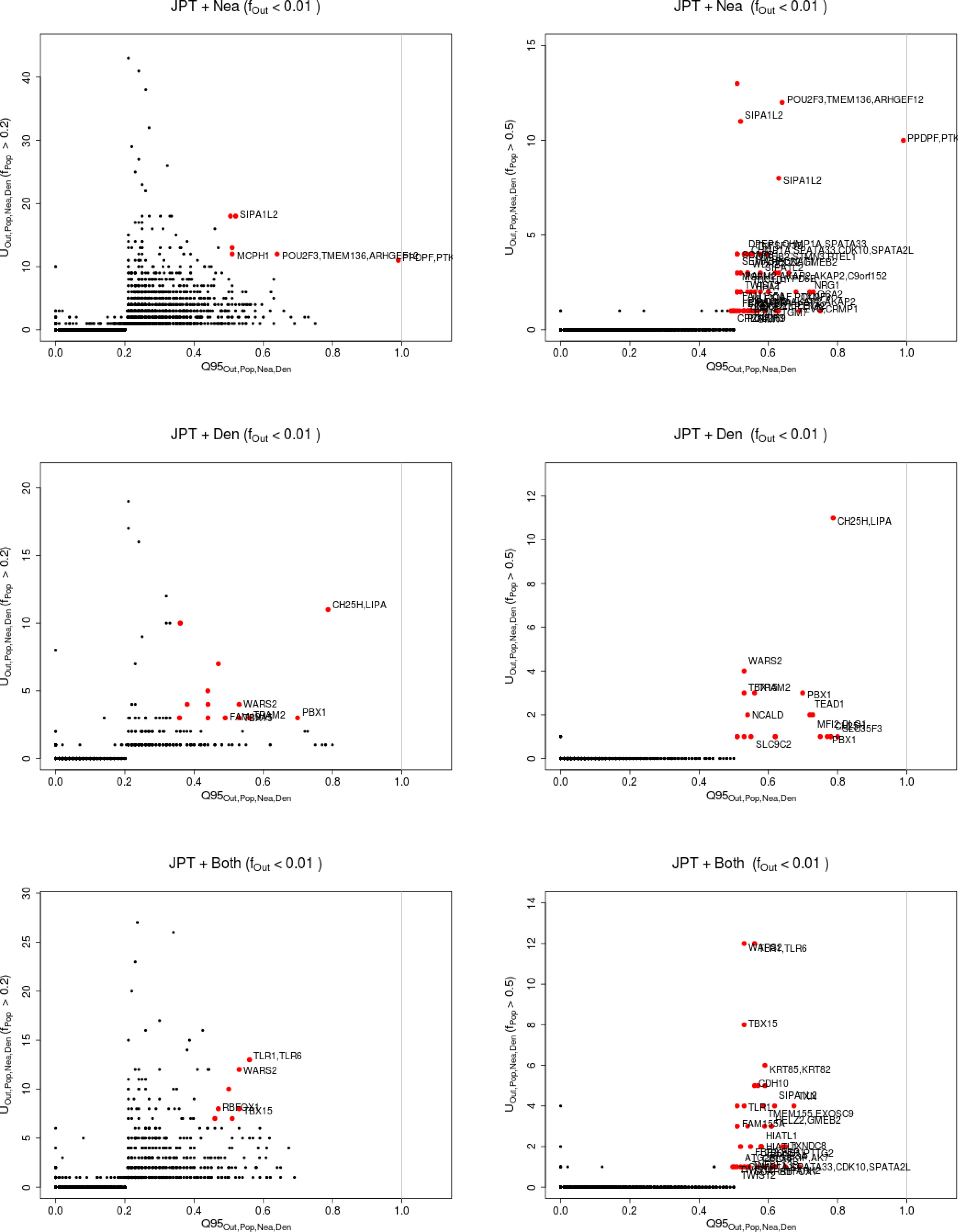
Uniquely shared archaic alleles in a Japanese (JPT) panel. Joint distribution of *Q*95_*AFR,JPT,Nea,Den*_(1%, *y, z*) and *U_AFR,JPT,Nea,Den_*(1%,x,y,z), for 40kb non-overlapping regions along the genome, using two choices of x (20% in left column panels,50% in right column panels). Red dots refer to regions that are in the 99.9% quantiles for both statistics. Neanderthal-specific shared alleles are displayed in the top panels, Denisovan-specific shared alleles are displayed in the middle-row panels, and alleles shared with both archaic human genome are displayed in the bottom panels.

**Figure S44:**
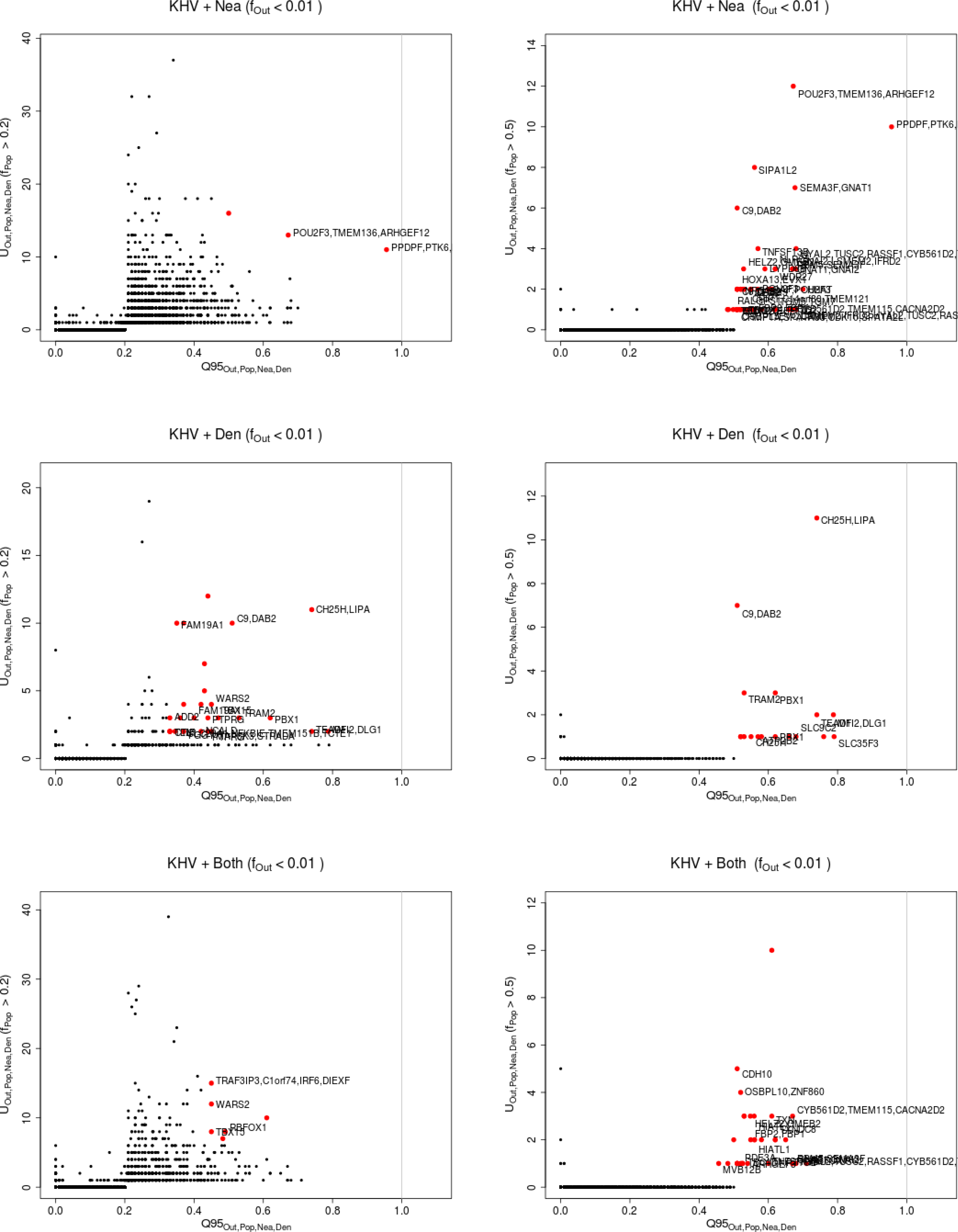
Uniquely shared archaic alleles in a Kinh (KHV) panel. Joint distribution of *Q*95_*AFR,KHV,Nea,Den*_(1%, *y, z*) and *U_AFR,KHV,Nea,Den_*(1%,x,y,z), for 40kb non-overlapping regions along the genome, using two choices of x (20% in left column panels,50% in right column panels). Red dots refer to regions that are in the 99.9% quantiles for both statistics. Neanderthal-specific shared alleles are displayed in the top panels, Denisovan-specific shared alleles are displayed in the middle-row panels, and alleles shared with both archaic human genome are displayed in the bottom panels.

**Figure S45:**
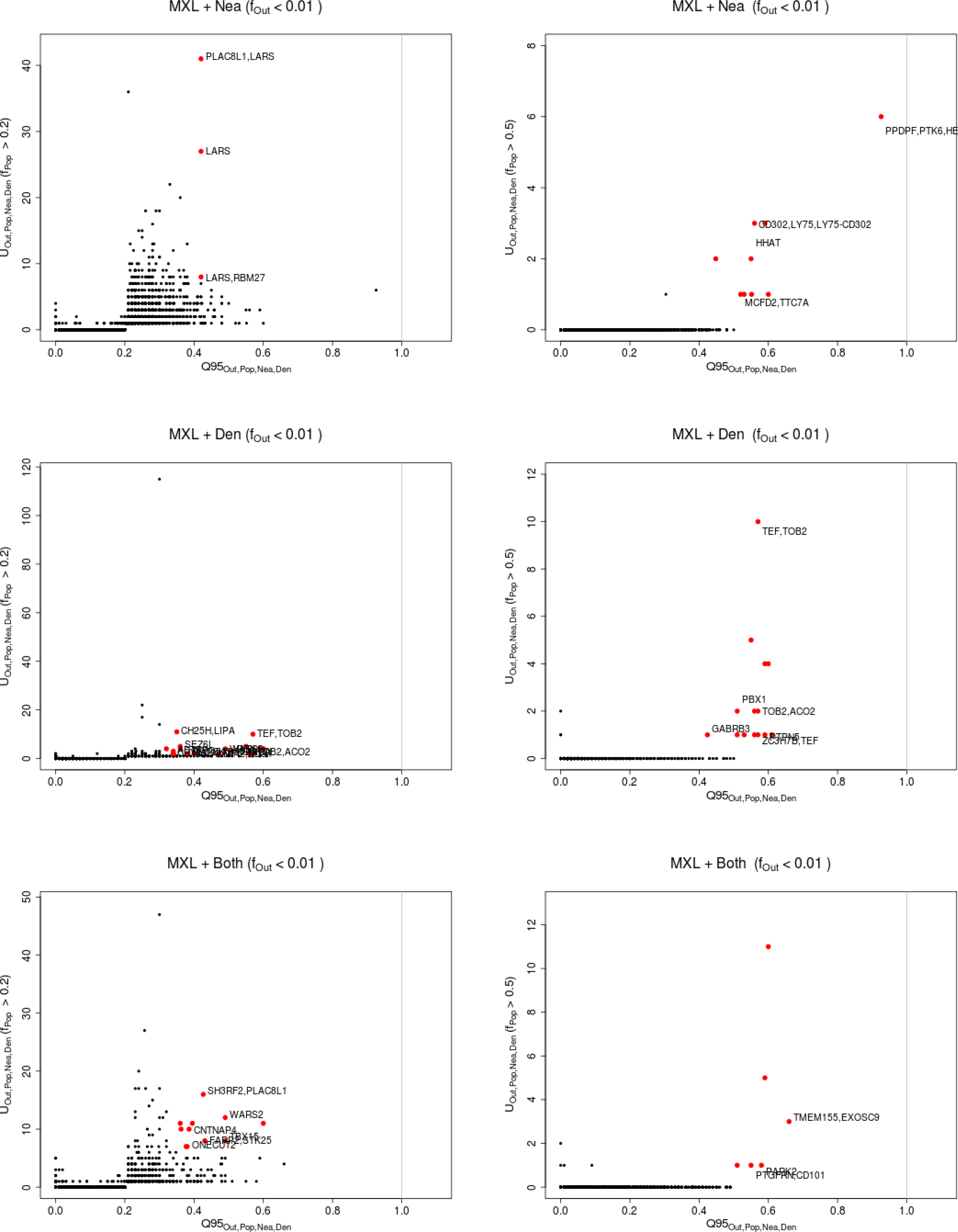
Uniquely shared archaic alleles in a Mexican (MXL) panel. Joint distribution of *Q*95_*AFR,MXL,Nea,Den*_(1%, *y, z*) and *U_AFR,MXL,Nea,Den_*(1%,x,y,z), for 40kb non-overlapping regions along the genome, using two choices of x (20% in left column panels,50% in right column panels). Red dots refer to regions that are in the 99.9% quantiles for both statistics. Neanderthal-specific shared alleles are displayed in the top panels, Denisovan-specific shared alleles are displayed in the middle-row panels, and alleles shared with both archaic human genome are displayed in the bottom panels.

**Figure S46:**
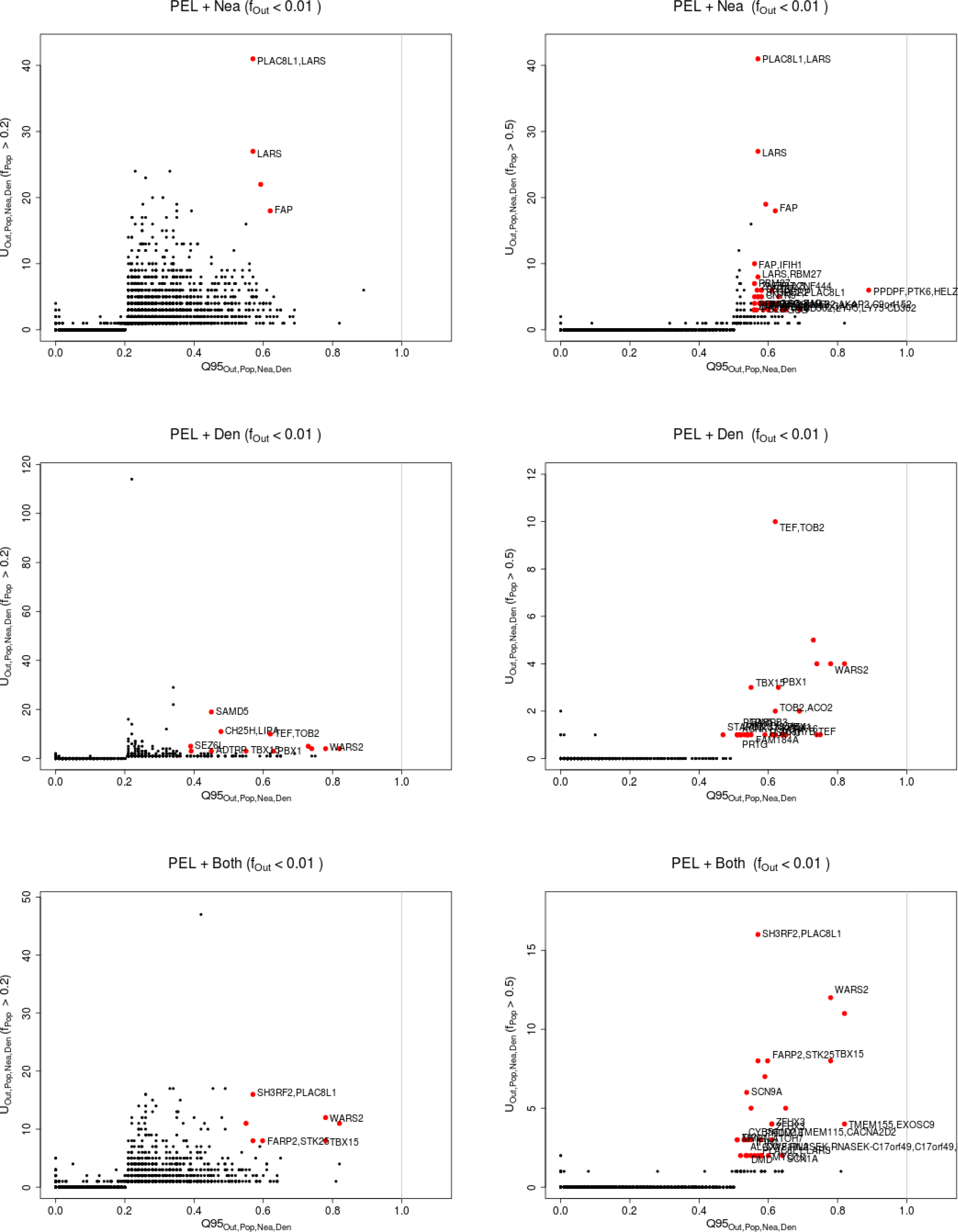
Uniquely shared archaic alleles in a Peruvian (PEL) panel. Joint distribution of *Q*95_*AFR,PEL,Nea,Den*_(1%, *y, z*) and *U_AFR,PEL,Nea,Den_*(1%,x,y,z), for 40kb non-overlapping regions along the genome, using two choices of x (20% in left column panels,50% in right column panels). Red dots refer to regions that are in the 99.9% quantiles for both statistics. Neanderthal-specific shared alleles are displayed in the top panels, Denisovan-specific shared alleles are displayed in the middle-row panels, and alleles shared with both archaic human genome are displayed in the bottom panels.

**Figure S47:**
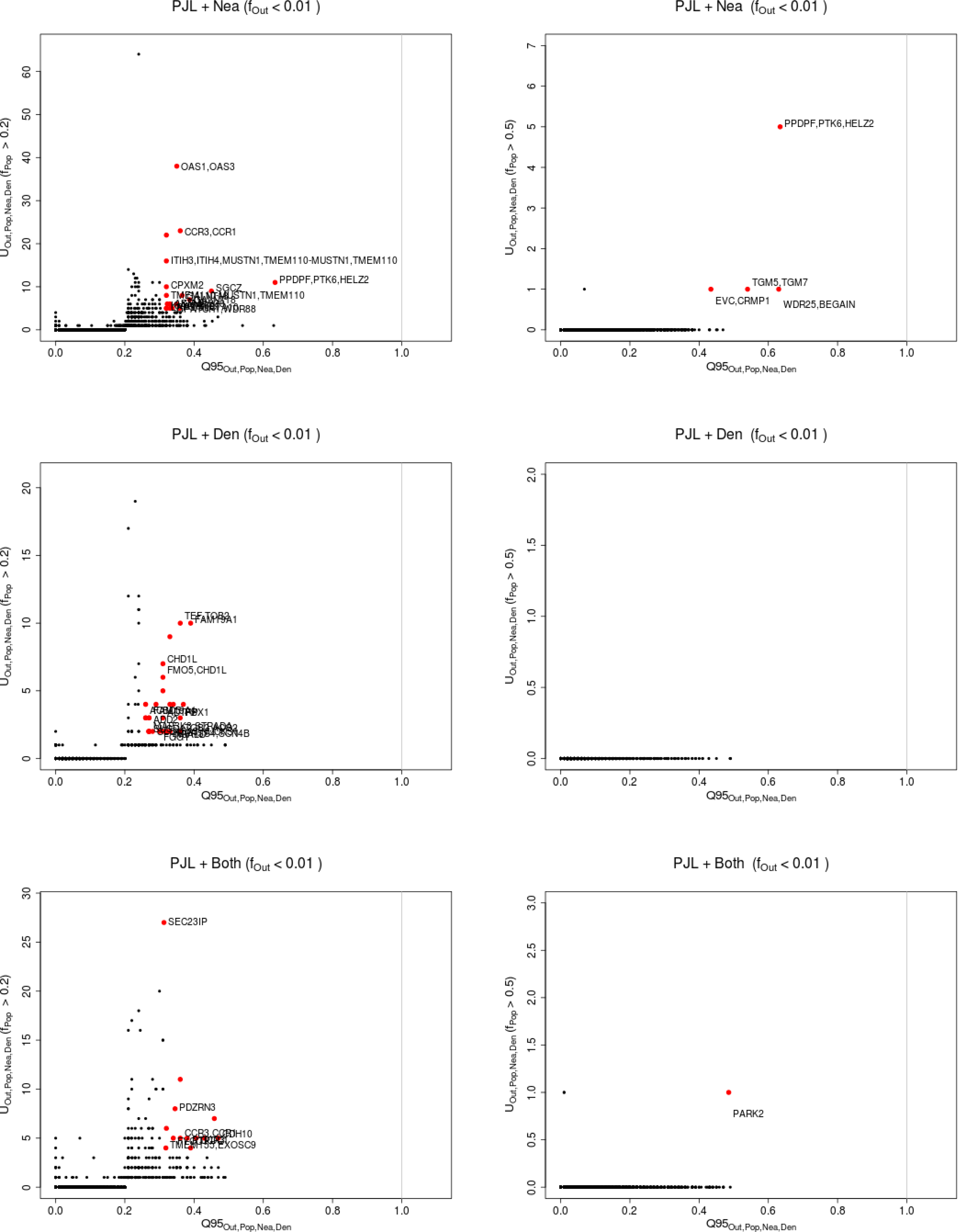
Uniquely shared archaic alleles in a Punjabi (PJL) panel. Joint distribution of *Q*95_*AFR,PJL,Nea,Den*_ (1%, *y, z*) and *U_AFR,PJL,Nea,Den_* (1%,x,y,z), for 40kb non-overlapping regions along the genome, using two choices of x (20% in left column panels,50% in right column panels). Red dots refer to regions that are in the 99.9% quantiles for both statistics. Neanderthal-specific shared alleles are displayed in the top panels, Denisovan-specific shared alleles are displayed in the middle-row panels, and alleles shared with both archaic human genome are displayed in the bottom panels.

**Figure S48:**
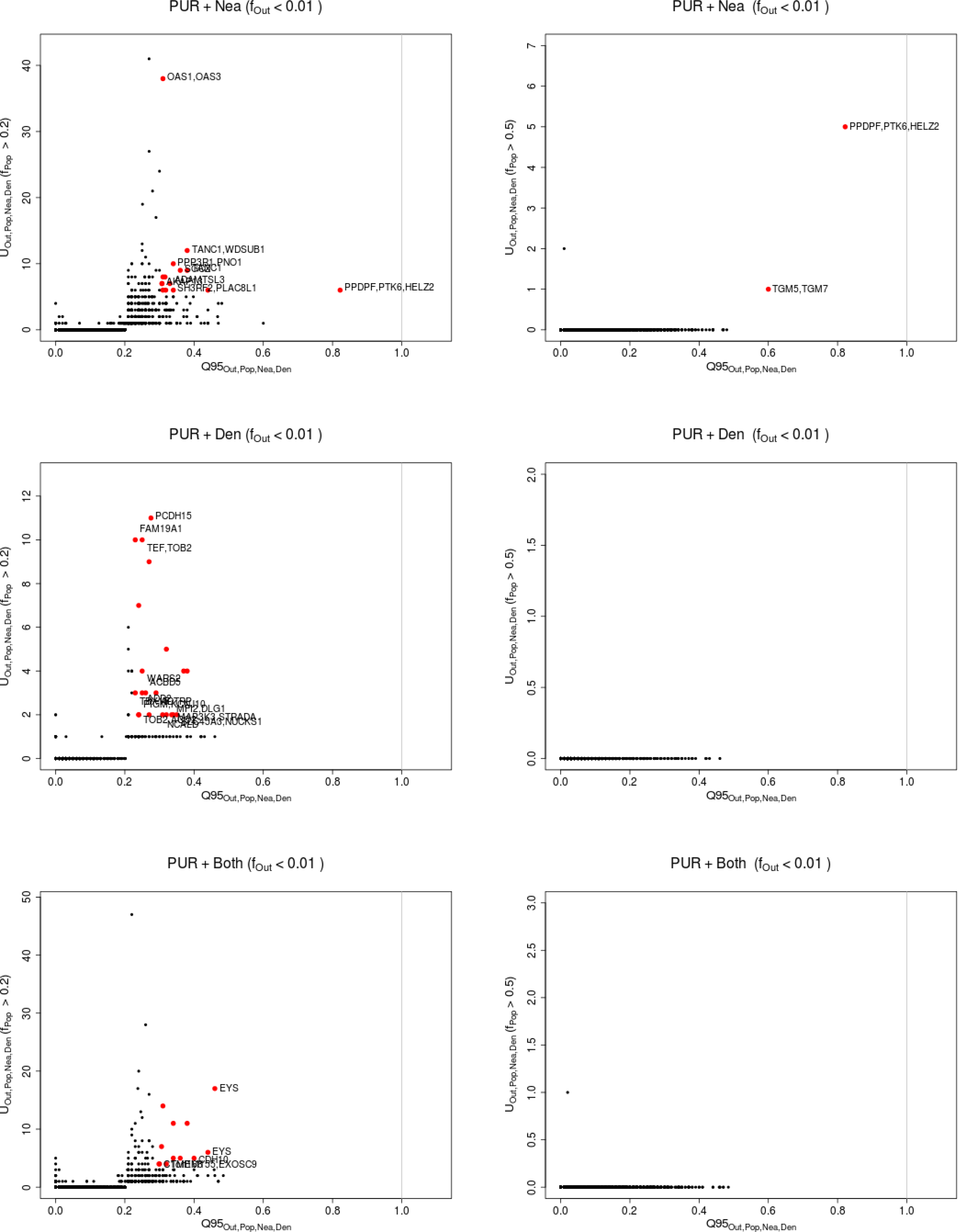
Uniquely shared archaic alleles in a Puerto Rican (PUR) panel. Joint distribution of *Q*95_*AFR,PUR,Nea,Den*_(1%, *y, z*) and *U_AFR,PUR,Nea,Den_*(1%,x,y,z), for 40kb non-overlapping regions along the genome, using two choices of x (20% in left column panels,50% in right column panels). Red dots refer to regions that are in the 99.9% quantiles for both statistics. Neanderthal-specific shared alleles are displayed in the top panels, Denisovan-specific shared alleles are displayed in the middle-row panels, and alleles shared with both archaic human genome are displayed in the bottom panels.

**Figure S49:**
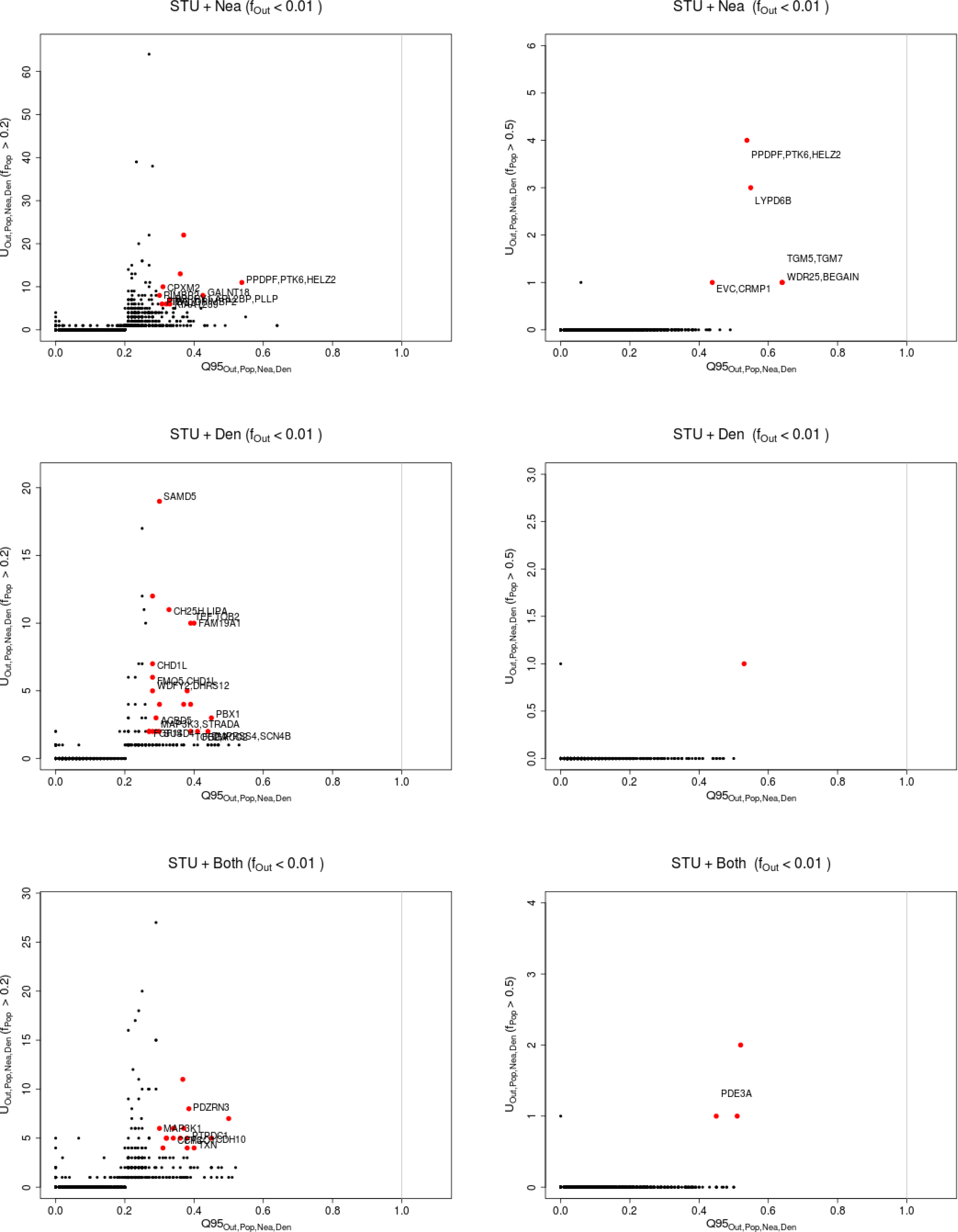
Uniquely shared archaic alleles in a Sri Lankan Tamil (STU) panel. Joint distribution of *Q*95_*AFR,STU,Nea,Den*_(1%, *y, z*) and *U_AFR,STU,Nea,Den_*(1%,x,y,z), for 40kb non-overlapping regions along the genome, using two choices of x (20% in left column panels,50% in right column panels). Red dots refer to regions that are in the 99.9% quantiles for both statistics. Neanderthal-specific shared alleles are displayed in the top panels, Denisovan-specific shared alleles are displayed in the middle-row panels, and alleles shared with both archaic human genome are displayed in the bottom panels.

**Figure S50:**
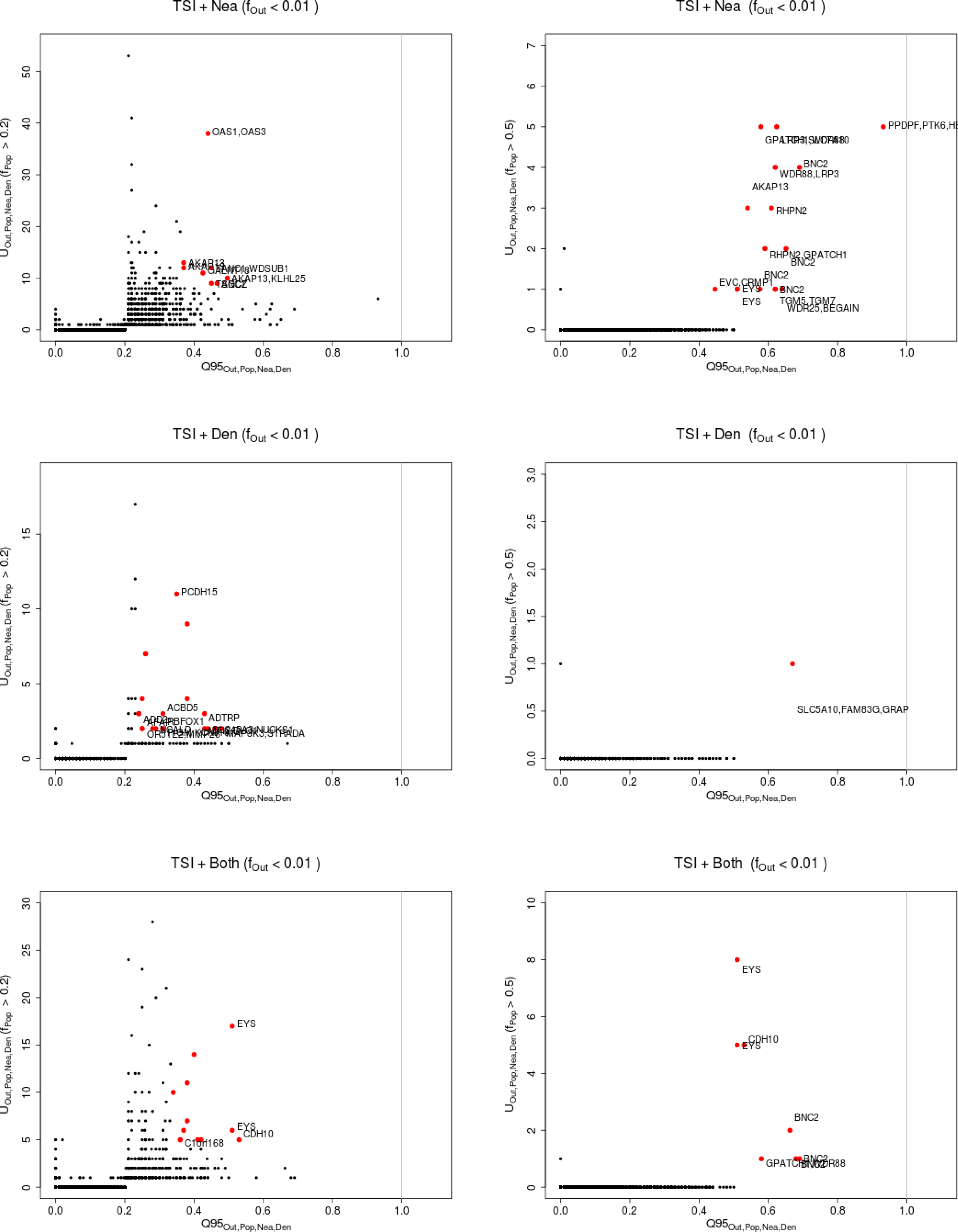
Uniquely shared archaic alleles in a Toscani (TSI) panel. Joint distribution of *Q*95_*AFR,TSI,Nea,Den*_ (1%, *y, z*) and *U_AFR,TSI,Nea,Den_*(1%,x,y,z), for 40kb non-overlapping regions along the genome, using two choices of x (20% in left column panels,50% in right column panels). Red dots refer to regions that are in the 99.9% quantiles for both statistics. Neanderthal-specific shared alleles are displayed in the top panels, Denisovan-specific shared alleles are displayed in the middle-row panels, and alleles shared with both archaic human genome are displayed in the bottom panels.

**Figure S51:**
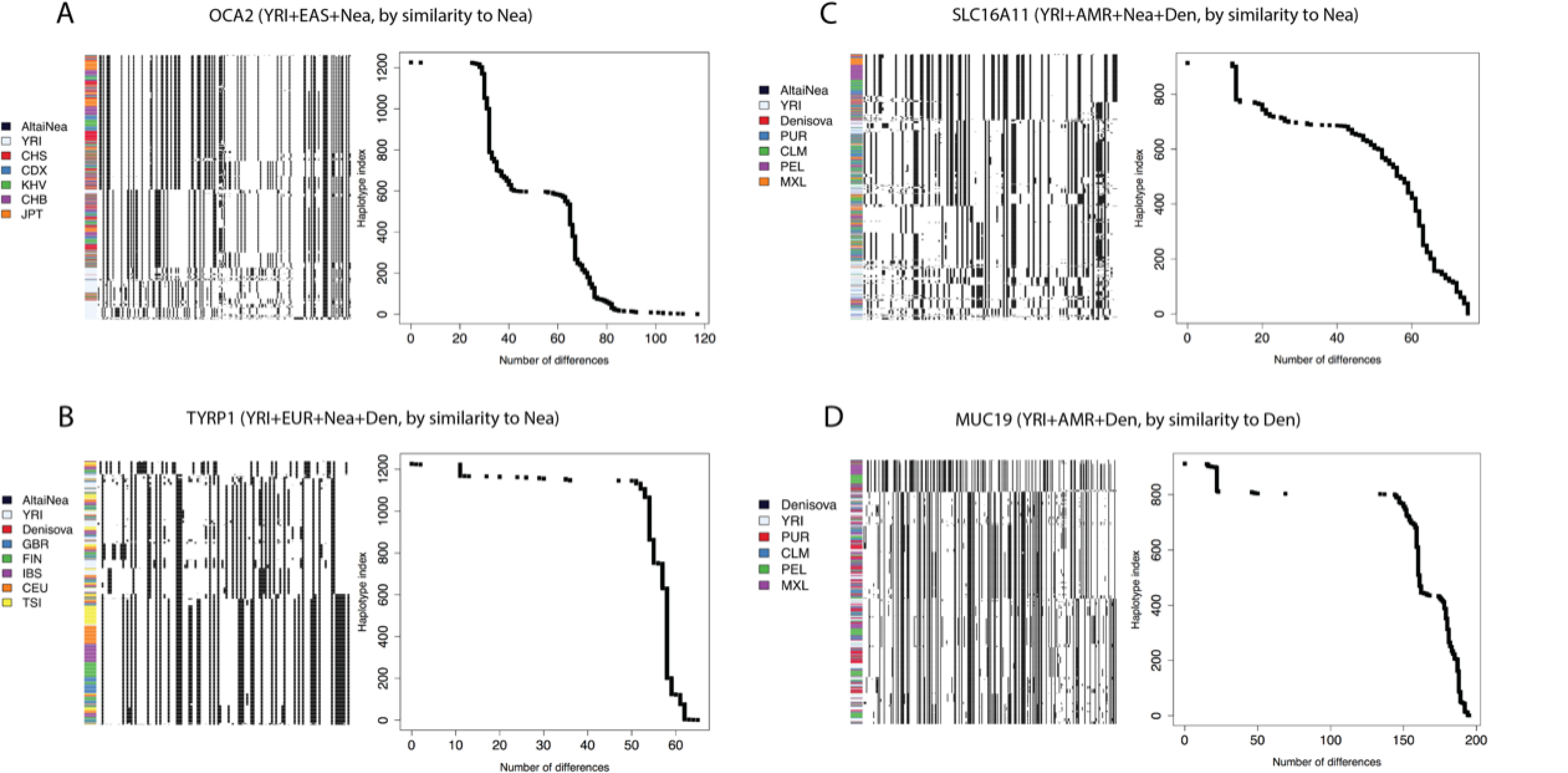
We explored the haplotype structure of *OCA2*, *TYRP1*, *SLC16A11* and *MUC19*. We applied a clustering algorithm to the haplotypes of particular human populations and then ordered the clusters by decreasing similarity to the archaic human genome with the larger number of uniquely shared sites. We also plotted the number of differences to the closest archaic haplotype for each human haplotype and sorted them simply by decreasing similarity. Note that, in the latter case, no clustering was performed, so the rows in the cumulative difference plots do not necessarily correspond to the rows in the adjacent haplotype structure plots. *OCA2*: chr15:28160001-28200000. *TYRP1*: chr9:12680001-12720000. *SLC16A11*: chr17:6880001-6960000. *MUC19*: chr12:40800001-40840000.

**Figure S52:**
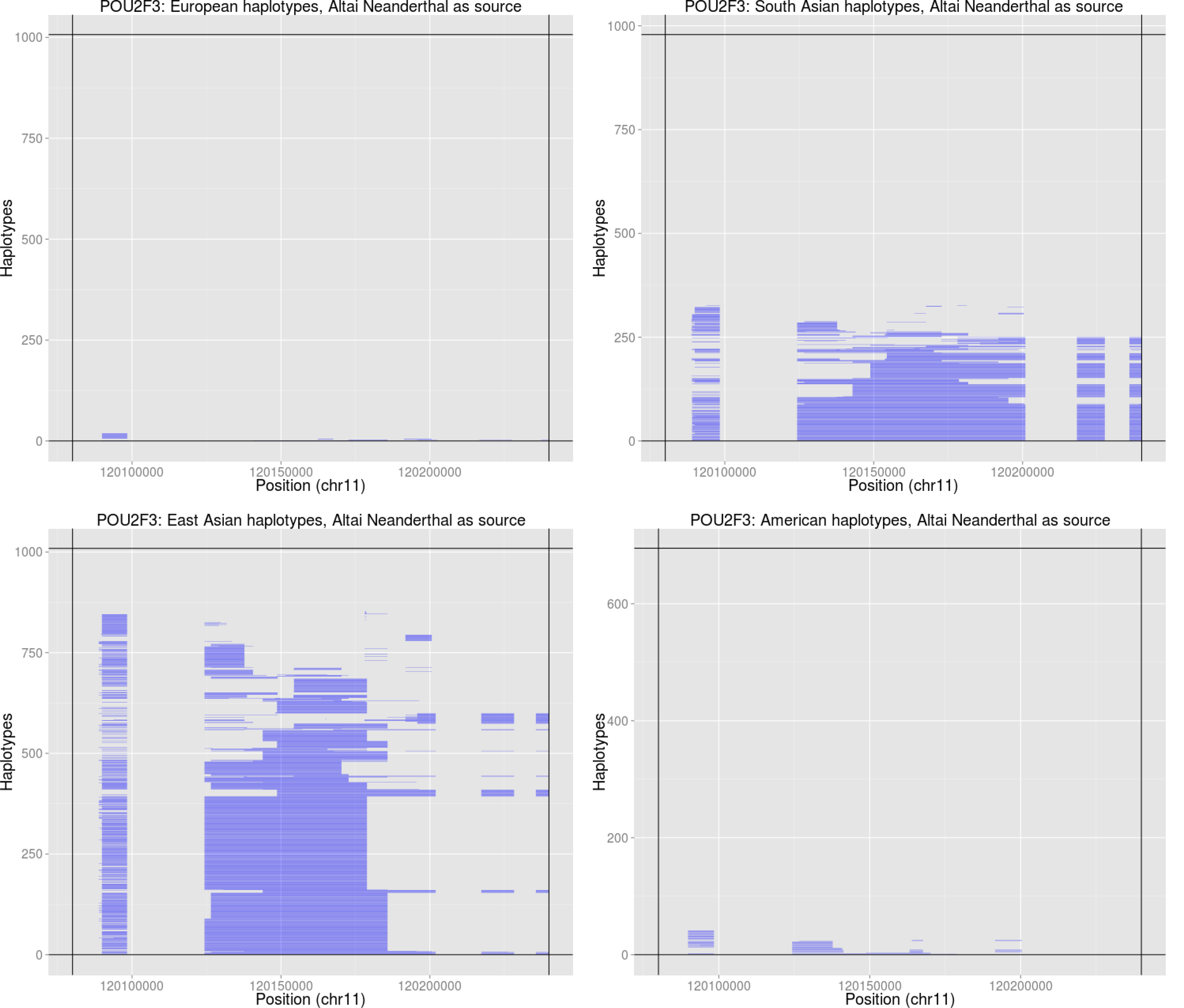
Introgressed tracks inferred in the four Non-African 1000 Genomes continental panels by an HMM [21] in the *POU2F3* region, using the Altai Neanderthal genome as the archaic source.

**Figure S53:**
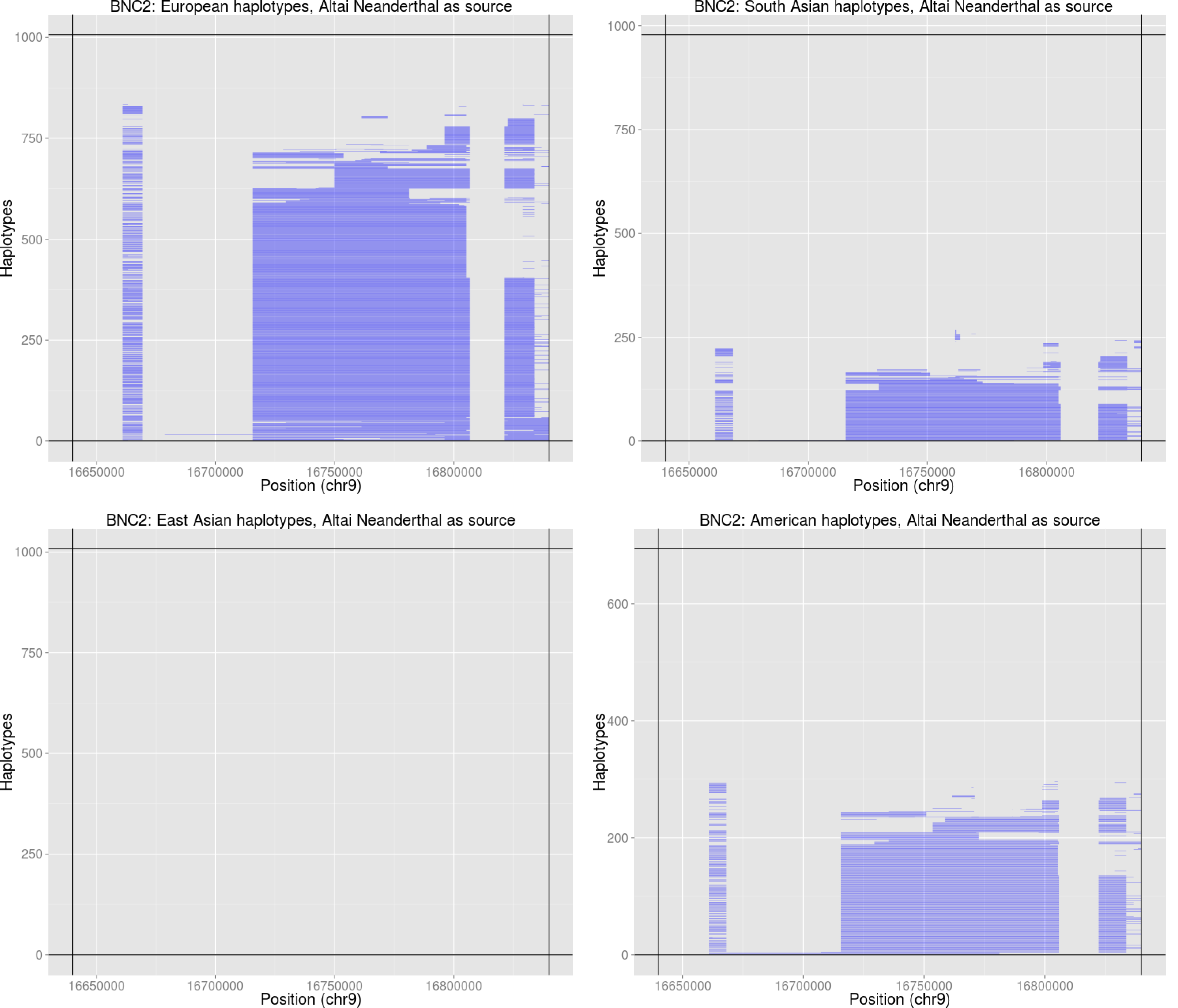
Introgressed tracks inferred in the four Non-African 1000 Genomes continental panels by an HMM [21] in the *BNC2* region, using the Altai Neanderthal genome as the archaic source.

**Figure S54:**
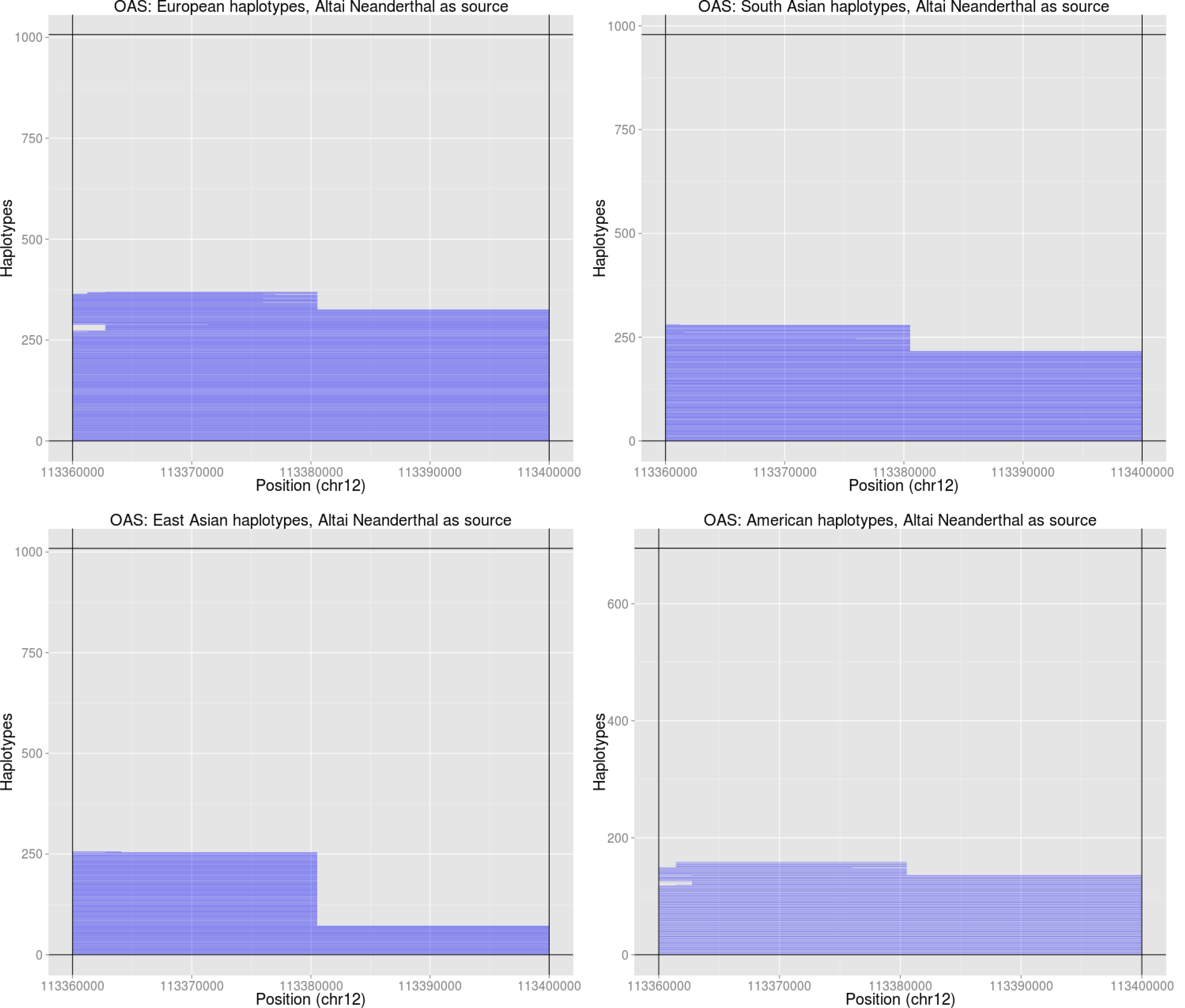
Introgressed tracks inferred in the four Non-African 1000 Genomes continental panels by an HMM [21] in the *OAS* region, using the Altai Neanderthal genome as the archaic source.

**Figure S55:**
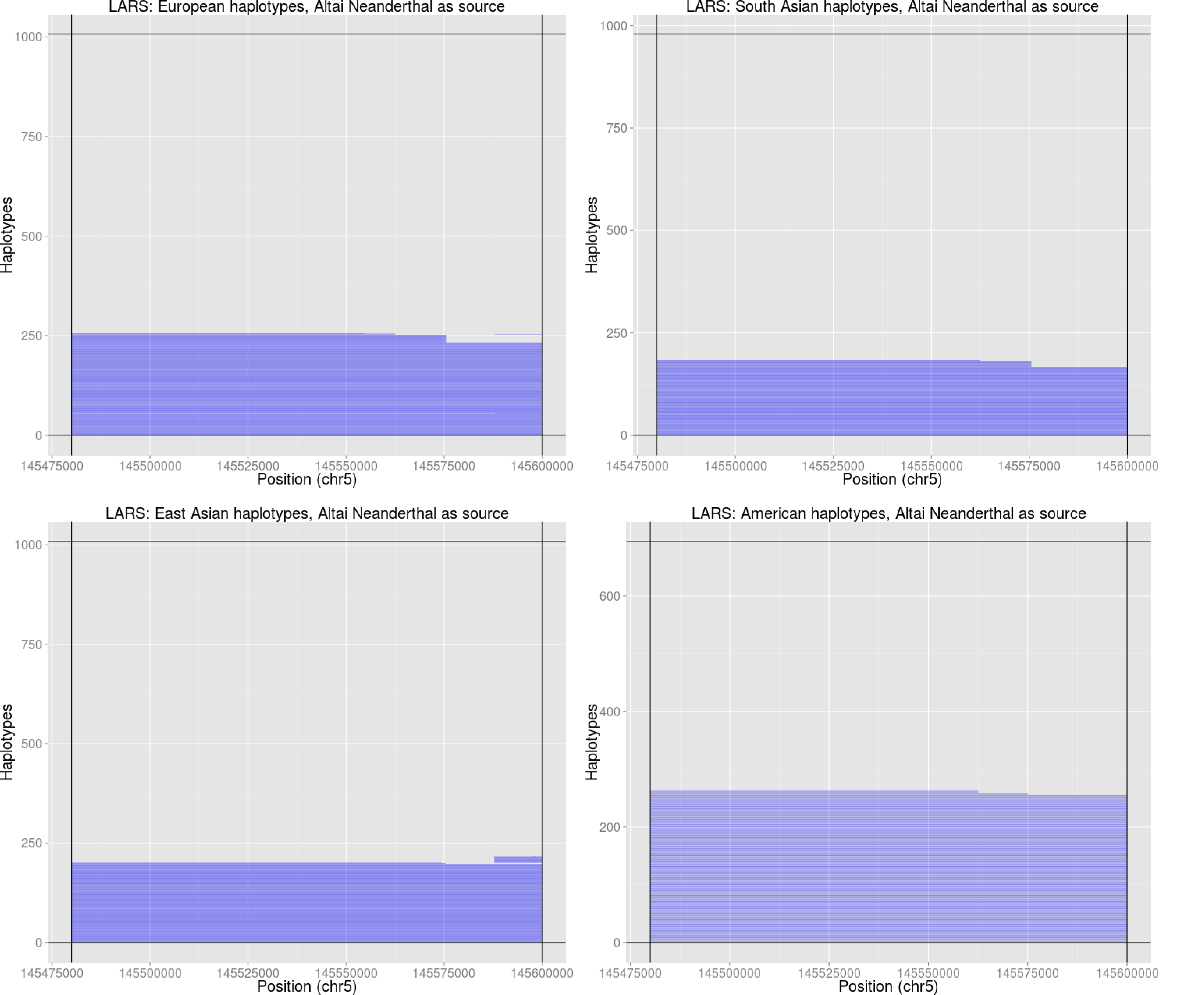
Introgressed tracks inferred in the four Non-African 1000 Genomes continental panels by an HMM [21] in the *LARS* region, using the Altai Neanderthal genome as the archaic source.

**Figure S56:**
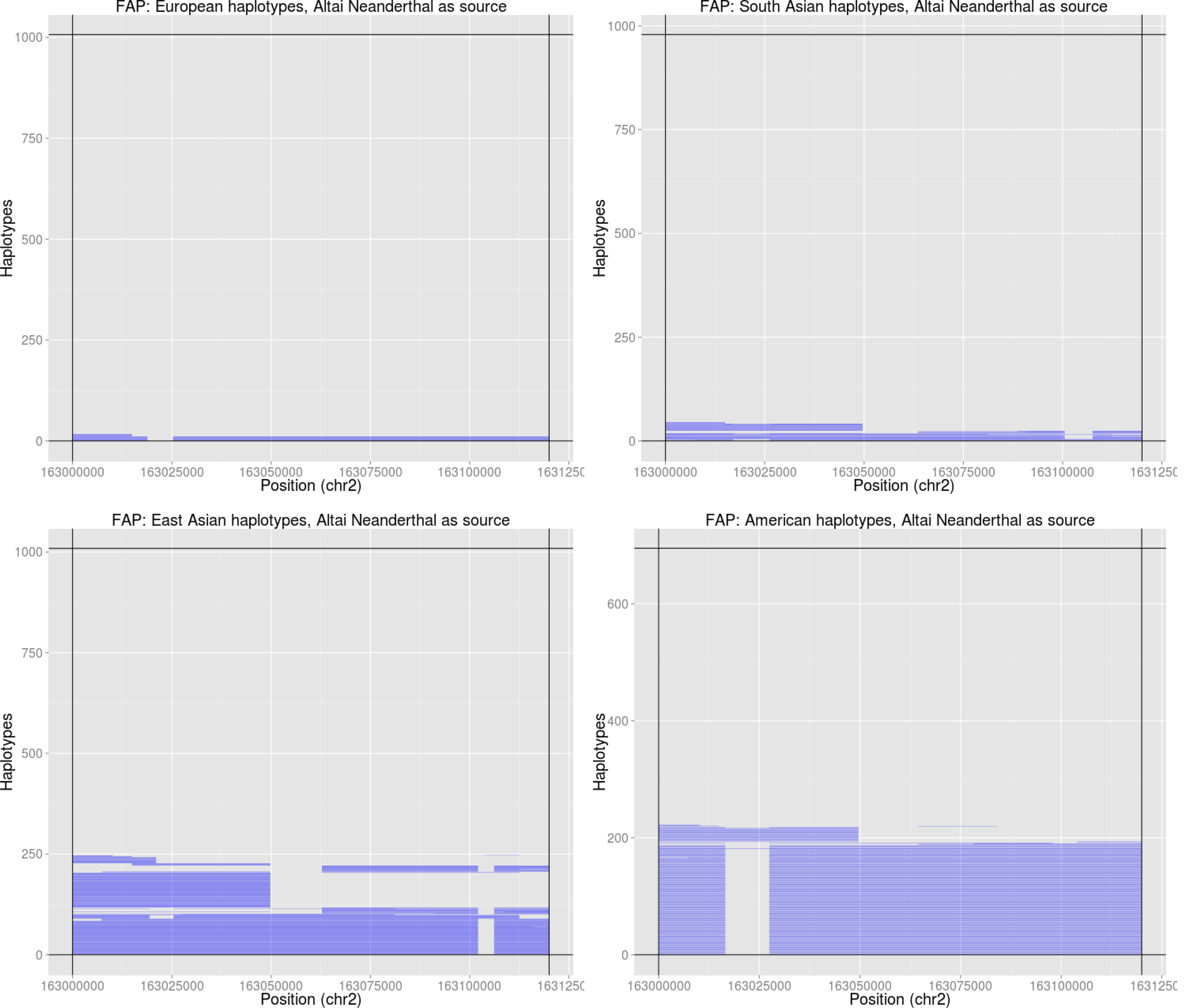
Introgressed tracks inferred in the four Non-African 1000 Genomes continental panels by an HMM [21] in the *FAP/IFIH1* region, using the Altai Neanderthal as the archaic source.

**Figure S57:**
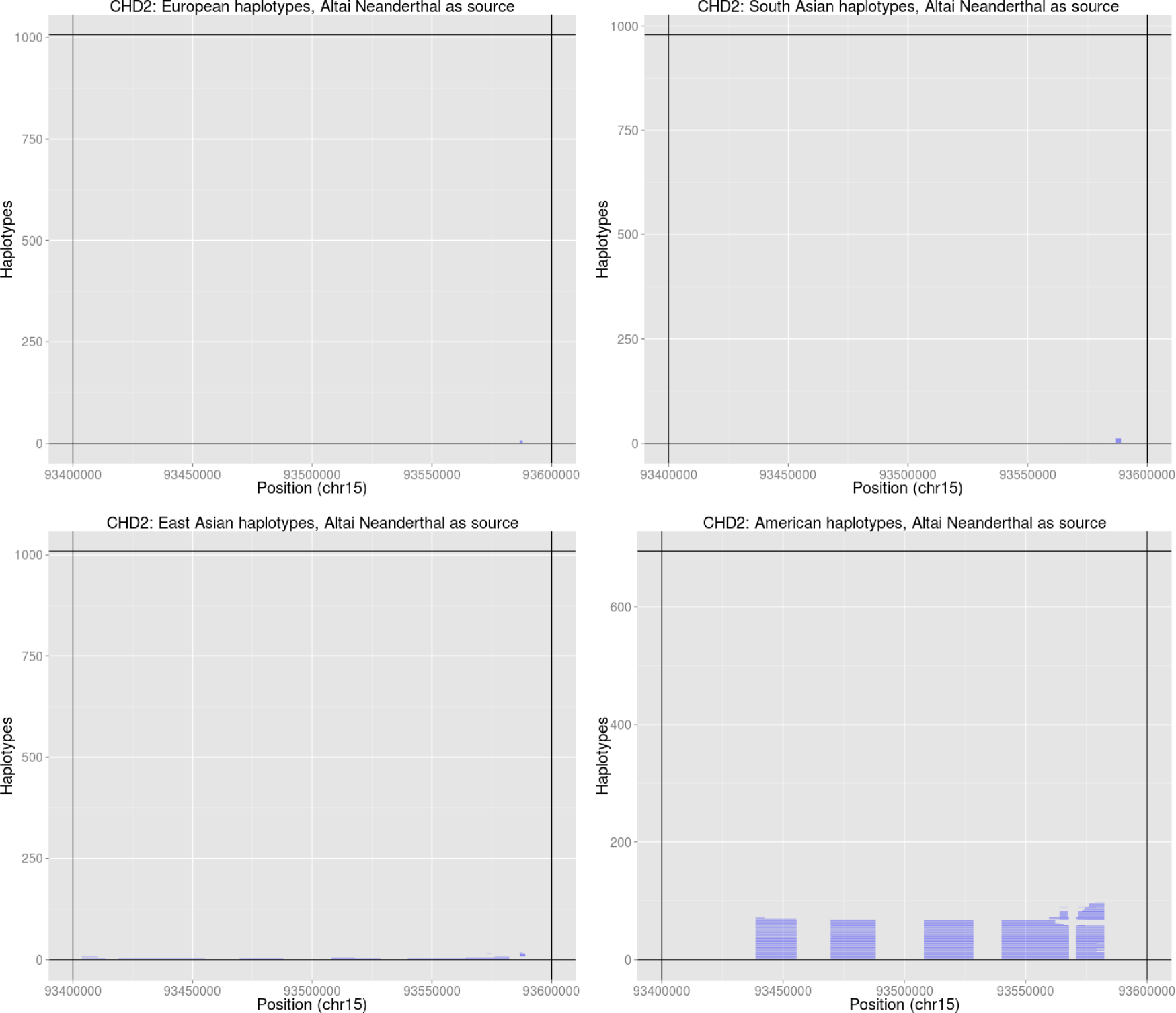
Introgressed tracks inferred in the four Non-African 1000 Genomes continental panels by an HMM [21] in the *CHD2* region, using the Altai Neanderthal genome as the archaic source.

**Figure S58:**
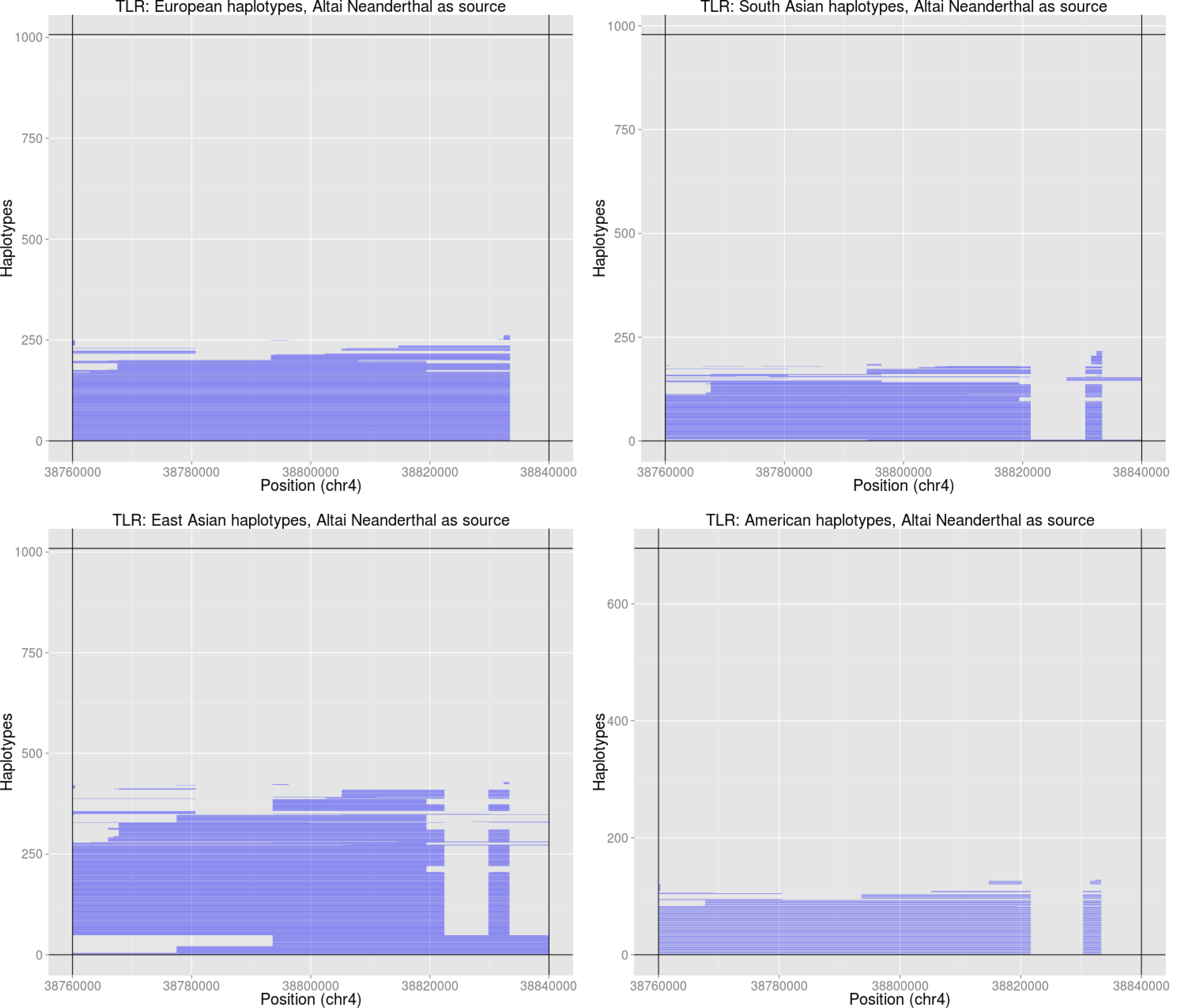
Introgressed tracks inferred in the four Non-African 1000 Genomes continental panels by an HMM [21] in the *TLR1-6* region, using the Altai Neanderthal genome as the archaic source.

**Figure S59:**
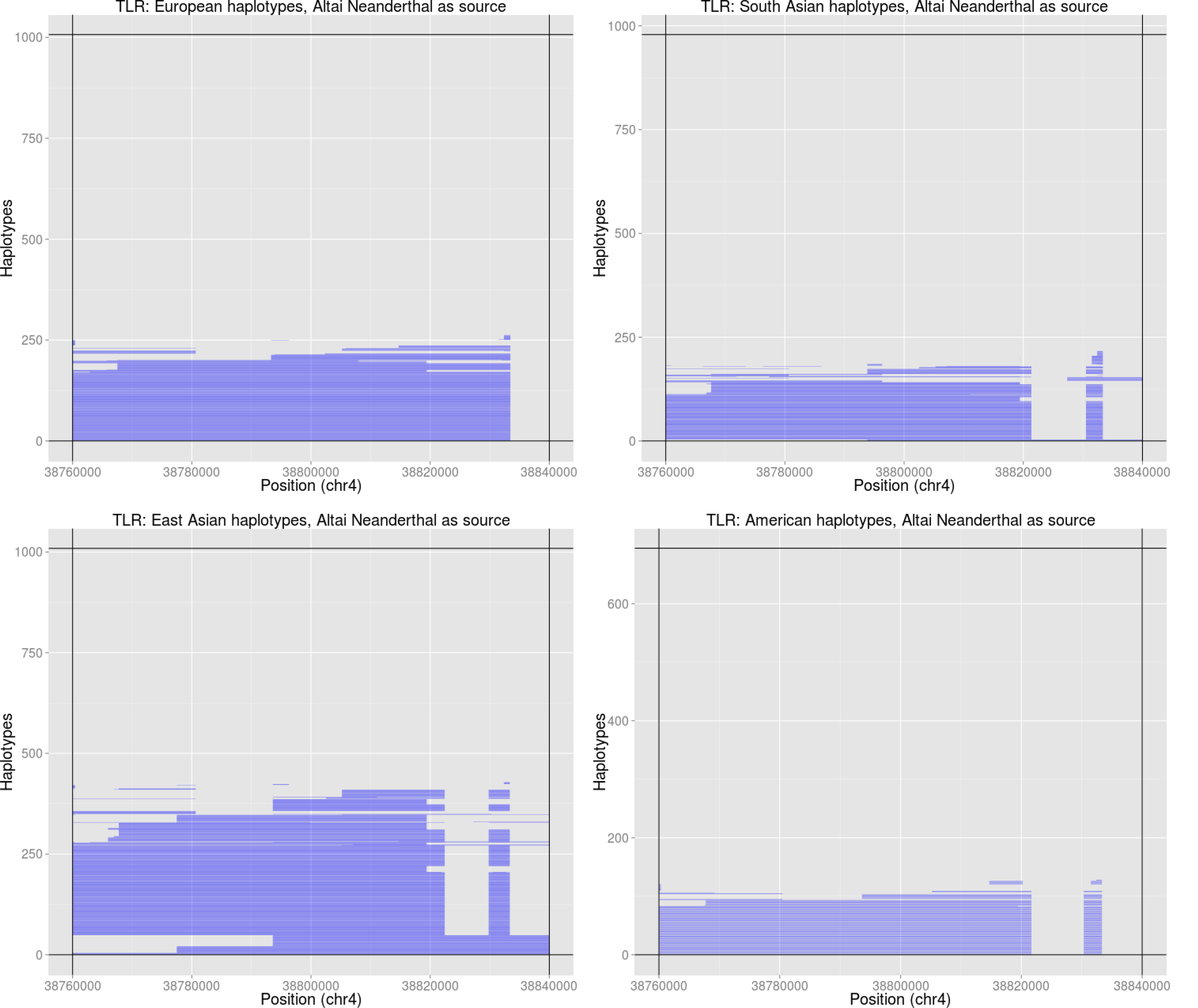
Introgressed tracks inferred in the four Non-African 1000 Genomes continental panels by an HMM [21] in the *ZFHX3* region, using the Altai Neanderthal genome as the archaic source.

**Figure S60:**
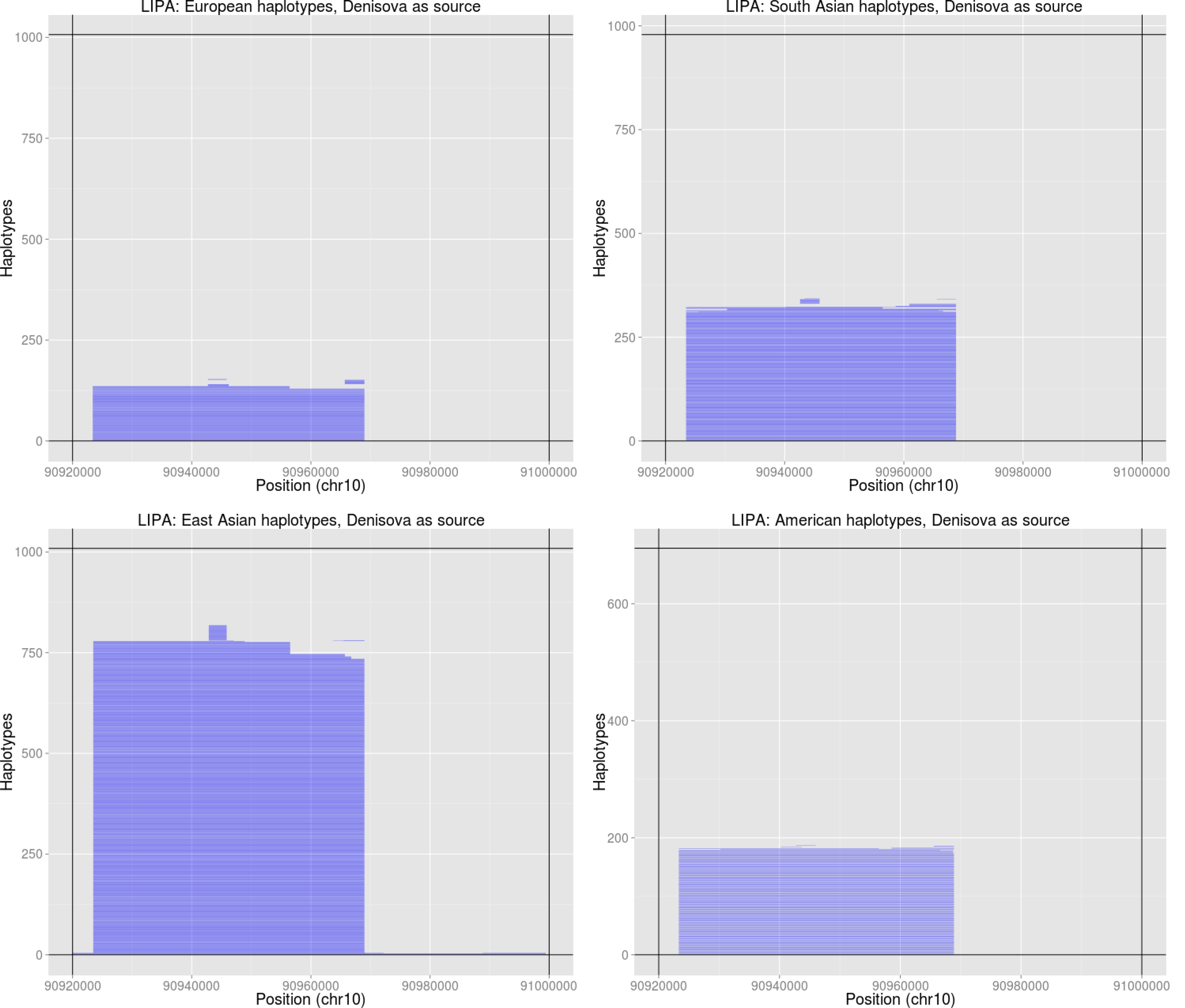
Introgressed tracks inferred in the four Non-African 1000 Genomes continental panels by an HMM [21] in the *LIPA* region, using the Denisova genome as the archaic source.

**Figure S61:**
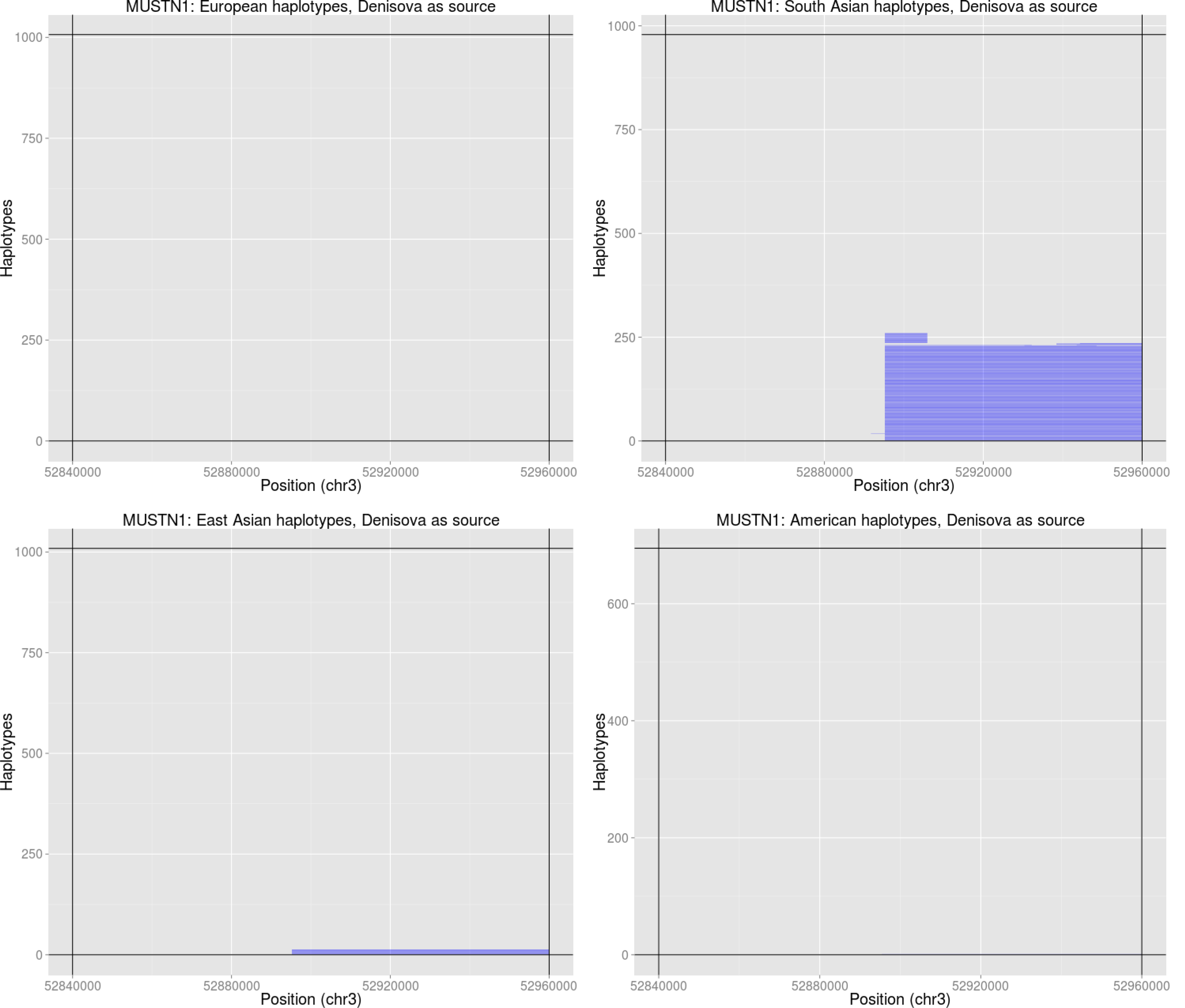
Introgressed tracks inferred in the four Non-African 1000 Genomes continental panels by an HMM [21] in the *MUSTN1* region, using the Denisova genome as the archaic source.

